# A spatially resolved single cell genomic atlas of the adult human breast

**DOI:** 10.1101/2023.04.22.537946

**Authors:** Tapsi Kumar, Kevin Nee, Runmin Wei, Siyuan He, Quy H. Nguyen, Shanshan Bai, Kerrigan Blake, Yanwen Gong, Maren Pein, Emi Sei, Min Hu, Anna Casasent, Aatish Thennavan, Jianzhuo Li, Tuan Tran, Ken Chen, Benedikt Nilges, Nachiket Kashikar, Oliver Braubach, Bassem Ben Cheikh, Nadya Nikulina, Hui Chen, Mediget Teshome, Brian Menegaz, Huma Javaid, Chandandeep Nagi, Jessica Montalvan, Delia F. Tifrea, Robert Edwards, Erin Lin, Ritesh Parajuli, Sebastian Winocour, Alastair Thompson, Bora Lim, Devon A. Lawson, Kai Kessenbrock, Nicholas Navin

## Abstract

The adult human breast comprises an intricate network of epithelial ducts and lobules that are embedded in connective and adipose tissue. While previous studies have mainly focused on the breast epithelial system, many of the non-epithelial cell types remain understudied. Here, we constructed a comprehensive Human Breast Cell Atlas (HBCA) at single-cell and spatial resolution. Our single-cell transcriptomics data profiled 535,941 cells from 62 women, and 120,024 nuclei from 20 women, identifying 11 major cell types and 53 cell states. These data revealed abundant pericyte, endothelial and immune cell populations, and highly diverse luminal epithelial cell states. Our spatial mapping using three technologies revealed an unexpectedly rich ecosystem of tissue-resident immune cells in the ducts and lobules, as well as distinct molecular differences between ductal and lobular regions. Collectively, these data provide an unprecedented reference of adult normal breast tissue for studying mammary biology and disease states such as breast cancer.

## Introduction

The human breast is an apocrine organ that plays an important physiological role in producing milk to nourish an infant after pregnancy^1^. This glandular function is mediated by an epithelial system consisting of highly branched lobular units producing milk that is transported via an intricate ductal network. The mammary epithelial system is embedded into an adipose-rich tissue and surrounded by a dense web of vasculature and lymphatic vessels that are directly connected to the regional lymph nodes (Fig. 1a). The human breast tissue is composed of four major spatial regions: 1) terminal ductal lobular units (TDLUs) and lobules of densely packed, branched epithelium, 2) tubular ducts of mostly bi-layered epithelium, 3) extracellular matrix (ECM)-rich connective tissue, and 4) adipose-rich regions. These areas consist of their own cellular neighborhoods and ecosystems that have been described in histopathological studies^1–3^. However, a comprehensive and systematic unbiased map of their cellular expression programs and spatial organizations in the breast remains lacking.

**Figure 1.**
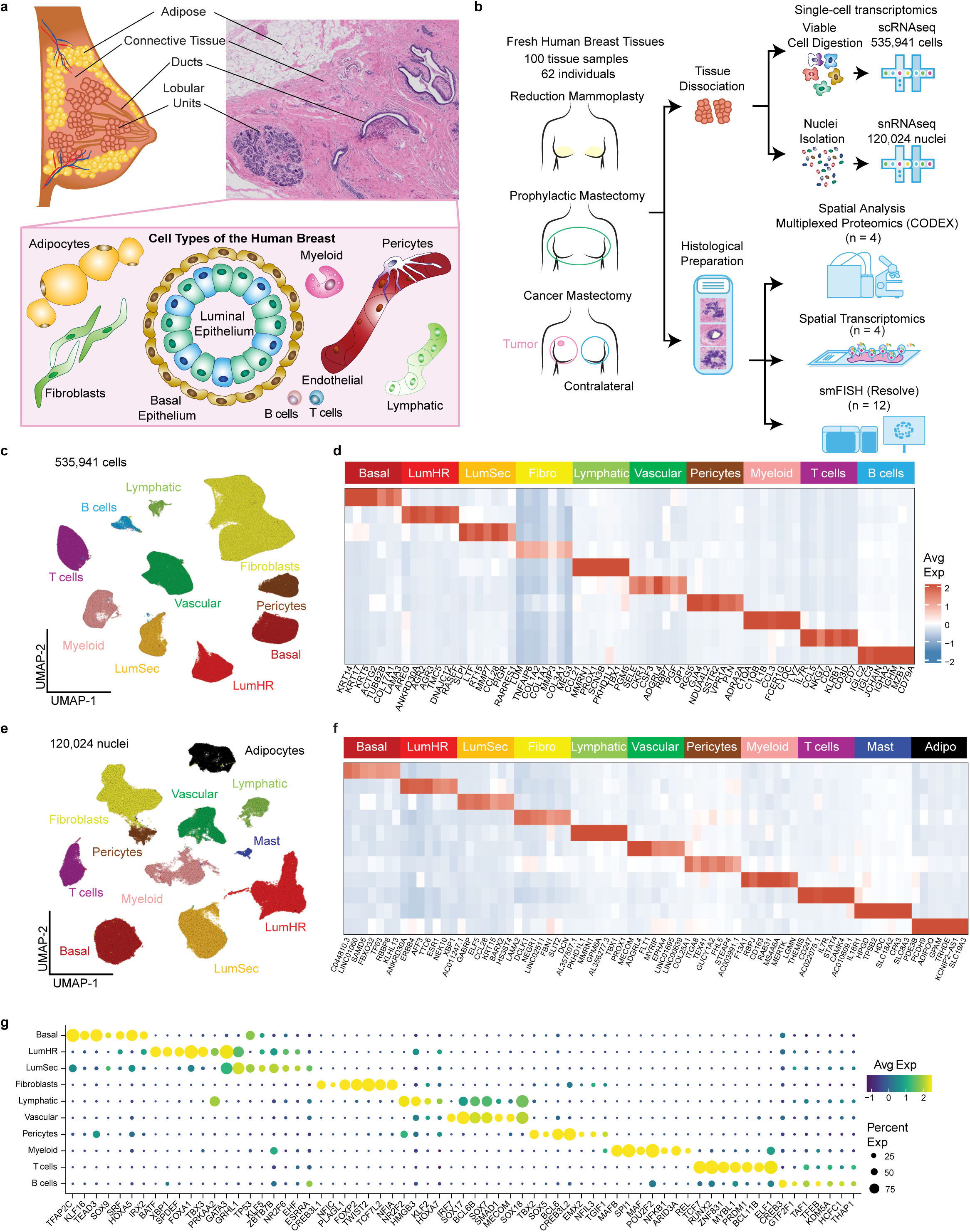
The Major Cell Types in the Adult Human Breast. **a,** Anatomy of the adult human breast and pathological H&E section, with illustrations showing the major cell types. **b**, Overview and workflow of the HBCA project, in which breast tissue samples were collected from different sources, followed by single cell and single nucleus RNA sequencing, as well as spatial analysis using CODEX, smFISH and ST. **c,** UMAP of scRNA-seq data from 535,941 cells integrated across 100 tissues from 62 women, showing 10 clusters that correspond to the major cell types. **d,** Consensus heatmap of the top 7 genes expressed in each cell type cluster from averaging scRNA-seq data. **e**, UMAP projection and clustering of 120,024 nuclei from snRNA-seq data across 20 tissues from 20 women, showing 11 clusters that correspond to the major cell types. **f**, Consensus heatmap of top 7 genes expressed from each cell type, averaged across the snRNA-seq data. **g,** Dot plot of the top 7 transcription factors identified from scRNA-seq data for each cell type cluster.

Previous studies have focused mainly on characterizing the epithelial cells, which comprise the inner layer of luminal cells and the outer layer of basal/myoepithelial cells within ducts and lobules^4–6^ (Fig. 1b). This focus on epithelium is mainly due to its implication in breast cancer^7, 8^, mammary stem cell and progenitor functions^9, 10^ and its dynamic changes during menstruation, pregnancy, and lactation^11, 12^. Previous studies using single-cell RNA sequencing (scRNA-seq) of normal human breast tissues identified three major mammary epithelial cell types, laying the groundwork for the present HBCA^4, 13, 6, 14, 15^. Increasing evidence also suggests that the microenvironment directly surrounding epithelial structures contains numerous stromal cells that actively crosstalk with the epithelial cells^16–19^. However, the intense focus on understanding epithelial cell biology has left a major gap in knowledge concerning the non epithelial cell types. The goal of the HBCA is to generate a comprehensive atlas using unbiased single-cell and spatial genomic methods, which is part of the larger Human Cell Atlas (HCA) project^20^.

### Major cell types in the adult human breast tissues

To identify the major cell types and their expression programs, we performed unbiased 3’ scRNA-seq (10X Genomics) from 100 tissue samples collected from 62 women under informed IRB consent (Fig. 1b, Supplementary Table 1). Fresh human breast tissue samples were obtained from disease-free women which included reduction mammoplasties (RM=30) and prophylactic mastectomies (PM=6), as well as breast cancer patients using the contralateral mastectomies (CM=26) from the other non-malignant breast. These fresh tissue samples were collected at four institutions within 1-2 hours after surgery using mirrored protocols, to generate viable cell suspensions from large tissue specimens (50-100g) (Methods). We sequenced an average of 9,143 cells per specimen at 50K average reads per cell (Supplementary Table 2).

After filtering, scRNA-seq data from 535,941 cells was integrated and clustered revealing 10 major breast cell types (Fig. 1c, Extended Data Fig. 1a, Methods). These cell types included three epithelial components: luminal hormone-responsive (LumHR), luminal secretory (LumSec) and basal/myoepithelial (Basal); two endothelial (lymphatic and vascular); three immune (T cells, B cells and myeloid cells) and two mesenchymal cell types (fibroblasts (Fibro) and Pericytes). While the frequency of the major cell types varied across women, all the cell types were detected in most women, irrespective of the tissue source (Extended Data Fig. 1b, 1g). Many cell types were consistent with histopathological^1, 2^ and molecular studies^13, 14^ of breast tissues, however the detection of a high number of pericytes (5.4% total cells) and immune cells (18.3% total cells) was unexpected. Many of the top expressed genes represented canonical cell type marker genes, thereby confirming their identities (Fig. 1d, Extended Data Fig. 1a). However, approximately half of the top marker genes have not previously been reported, providing a valuable resource for isolating or labelling specific breast cell types (Supplementary Table 3).

Notably, the scRNA-seq data did not identify any adipocytes, which represent a major component of breast tissue based on histological data^2, 21^. This issue was likely due to the large cell size of adipocytes (>50 microns), preventing their encapsulation on the scRNA-seq microdroplet platform^22^. To identify adipocytes and other challenging cell types, we also performed single nucleus RNA-seq (snRNA-seq) of 120,024 cells from 20 human breast tissue samples (Fig. 1e, Supplementary Table 1). Our snRNA-seq analysis detected all major cell types identified by scRNA-seq (except for B cells) and additionally included adipocytes and mast cells (Fig. 1f, Extended Data Fig. 1c, h). The frequency of cell types differed for some cell types (e.g., fibroblasts) in the snRNA-seq and scRNA-seq data, reflecting differences in cell proportions after tissue dissociation (Extended Data Fig. 1h). Additionally, the top cell type marker genes in snRNA-seq often differed from the scRNA-seq data, likely reflecting biological differences in the cytoplasmic and nuclear RNA pools (Supplementary Tables 3,4). To identify key transcription factors (TFs), we performed a regulon analysis^23^ using both the scRNA-seq data (Fig. 1g) and the snRNA-seq datasets (Extended Data Fig. 1d). These data identified many known^24^ and novel TFs that regulate breast cell type identities.

One potential concern was that sampling different spatial areas in the breast could potentially lead to differences in cell type compositions. To investigate this issue, we compared the cell type frequencies from matched (left/right) breasts from 22 women, which showed no significant differences based on Procrustes analysis (R=0.83, p=1.5e-6) (Extended Data Fig. 1e,f, Methods). We also compared cell type frequencies across the three main tissue sources, which showed only minor differences (Extended Data Fig. 1g). Collectively, these data identified 11 major cell types in adult mammary breast tissues.

### Spatial mapping reveals cellular neighborhoods in breast tissues

By histopathology human breast tissue can be divided into four major topographic regions: adipose tissue (A), connective tissue (C), and epithelial-rich regions, which are further subclassified into terminal ductal lobular units (TDLU), herein referred to as lobules (L), and ductal (D) regions^2, 3^ (Fig.1a). We utilized three orthogonal technologies to investigate the spatial organization of cell types *in situ*, including unbiased spatial transcriptomics (ST, 10X Genomics)^25^, targeted single molecule RNA FISH (smFISH, Resolve Biosciences)^26^ and co-detection by indexing (CODEX, Akoya Biosciences) for proteomic analysis^27^.

ST analysis was performed on tissues from 4 patients and the data was integrated and clustered, revealing 10 major ST clusters (Fig. 2a-b, Extended Data Fig. 2a, c-e, Supplementary Table 5). Although ST spots are 55 microns and could potentially be mixtures of cell types, a direct comparison to the scRNA-seq clusters showed that most of the ST clusters corresponded to a single dominant breast cell type cluster, except for ST03 and ST04 (Extended Data Fig. 2b). Importantly, ST cluster signatures showed a high concordance with the top cell type marker genes defined by scRNA-seq, validating markers *in situ* (Extended Data Fig. 2f). Notably, the abundance and distribution of ST clusters corresponded to the four different tissue areas (A, C, D, L) that were annotated by histopathology. Regions A and C had a high proportion of ST06 adipocyte and ST05 fibroblast clusters, respectively, while region D contained higher proportions of the ST04 LumSec and ST01 myoepithelial clusters and region L contained a higher proportion of the ST03 and ST02 epithelial clusters (Fig. 2c).

**Figure 2.**
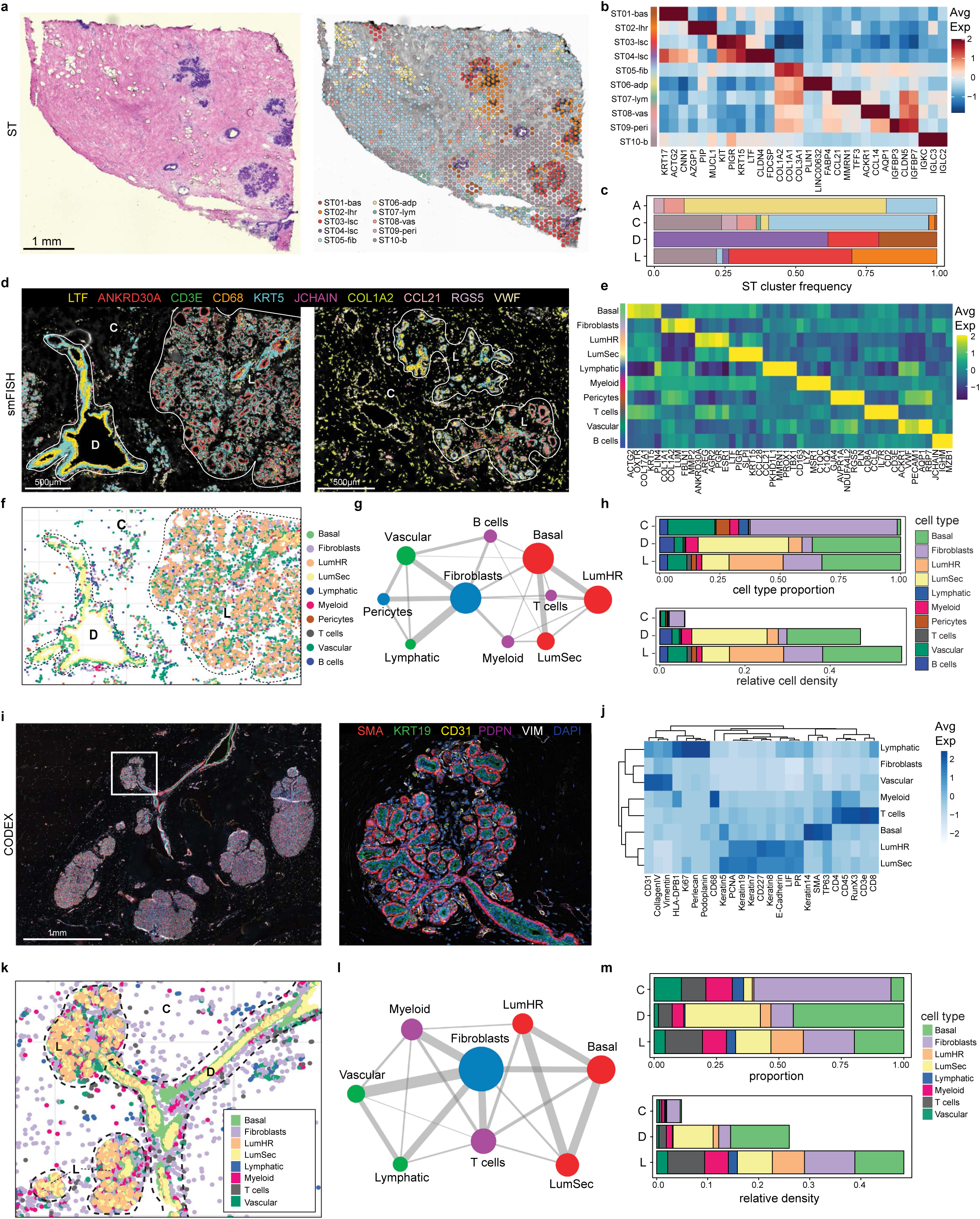
Spatial Analysis of Cell Types in Human Breast Tissues. **a,** ST experiment from P35, showing the unbiased clustering and histopathological image with topographic regions annotated. **b,** Heatmap of top 3 marker genes in each ST cluster for 4 tissue samples, showing the dominant cell type identity assigned. **c,** Frequencies of the ST clusters across the 4 topographic tissue areas in 4 combined tissue samples. **d,** smFISH experiments (Resolve Biosciences) using a custom 100-gene panel, showing a subset of 10 genes that mark different cell types in two women (P46-S1 and P47-S1). **e,** Heatmap of the top 5 targeted maker genes for each cell type in the smFISH data from 12 combined tissue samples. **f,** smFISH data after cell segmentation using combinations of markers. **g,** Spatial colocalization of cell types in smFISH data from 12 tissues, where node size represents the cell number and edge width represents the probability of colocalization. **h,** Frequencies of cell types and cell densities across three topographic areas using 12 tissues profiled by smFISH. **i**, CODEX data from P66 showing ductal-lobular structure with 6 protein markers, with an expanded region of one lobule in the right panel. **j**, Heatmap showing protein levels for markers that were used to identify different cell types. **k**, Cell segmentation using combinations of markers to identify cell types in the CODEX data from one tissue sample, with topographic areas annotated. **l**, Spatial colocalization graph of segmented cell types in the CODEX data from 4 tissues. **m**, Cell type and cell density frequencies from the CODEX data summarized across 4 tissue samples.

Since ST spots contain mixtures of cells, we also applied smFISH using a custom 100- gene panel based on the top expressed marker genes from the scRNA-seq data to distinguish cell types at single cell resolution (Supplementary Table 6, Methods). The smFISH analysis of 12 breast tissues from 5 women validated many top marker genes *in situ* (Fig. 2d-e, Supplementary Table 7). Most of the immune cell markers of T cells (*CD3E*), B cells (*JCHAIN*), and macrophages (*CD68*) were identified in the lobular and ductal regions, confirming their abundance in the scRNA-seq data (Fig. 2d). We further performed cell segmentation using combinations of the top markers of each cell type (Fig. 2f, Extended Data Fig. 3a, Methods). Using this data we computed a cell neighborhood proximity graph, which showed that the three epithelial cell types co-localized with B cells and T cells, while fibroblasts co-localized with vascular and lymphatic cells (Fig. 2g, Methods). We also quantified the cell type frequency in three major tissue areas (C, L, D), which showed that C was composed mainly of fibroblasts and vascular endothelial cells, while D consisted mainly of basal cells and high levels of LumSec epithelial cells, and L was composed of basal cells and high levels of LumHR epithelial cells, as well as fibroblasts (Fig. 2f, h, Extended Data Fig. 3a-b). The smFISH data also showed that overall cell density was low in connective tissue regions, and high in ductal and lobular regions.

We also investigated the spatial distribution of breast cell types in 4 patients using CODEX with a 34 antibody panel, which resolved 8 major cell types including T cells and myeloid cells (Supplementary Tables 8-9). One advantage of CODEX is that large tissue areas (approximately 1cm^2^) can be imaged, as well single cells in larger ducts and TDLUs (Fig. 2i). To perform a quantitative analysis, we performed cell segmentation and unsupervised clustering, followed by label transfer from the scRNA-seq data (Fig. 2j,k, Methods). This analysis showed that most cell types were consistent between the 4 tissue specimens in the CODEX data (Extended Data Fig. 3c). We also performed a proximity analysis, which was consistent with the smFISH data, but placed T cells closer to the fibroblasts (Fig. 2l). This data was used to define the cellular composition of the topographic regions at the protein level, which was consistent with the smFISH data (Fig. 2m).

### Epithelial cell diversity in the ducts and lobules

By histopathology, the ductal-lobular system of the breast consists of bi-layered epithelial cells, with an outer layer of basal cells and an inner layer of luminal cells^1, 2^ (Fig. 3a). However, additional epithelial subsets including stem and progenitor populations have been proposed within both the basal and luminal compartments^16, 10, 13, 14^. Our unbiased clustering of 114,967 epithelial cells and 56,280 epithelial nuclei identified three major epithelial cell types: Basal, LumSec, LumHR, which comprised a large proportion of the breast tissue (21.5% of cells, 45.4% of nuclei) and were consistent with previous scRNA-seq studies of human breast tissues^4, 6, 13, 14^ (Fig. 3b,c). As expected, nuclear expression of genes encoding hormone receptors (*ESR1*, *AR* and *PGR*) was specific to LumHR cells (Fig. 3d). We also performed an analysis of cytokeratin gene expression, which revealed basal specific, LumSec specific, and LumHR specific cytokeratins (Fig. 3e, Extended Data Fig. 4a-b).

**Figure 3.**
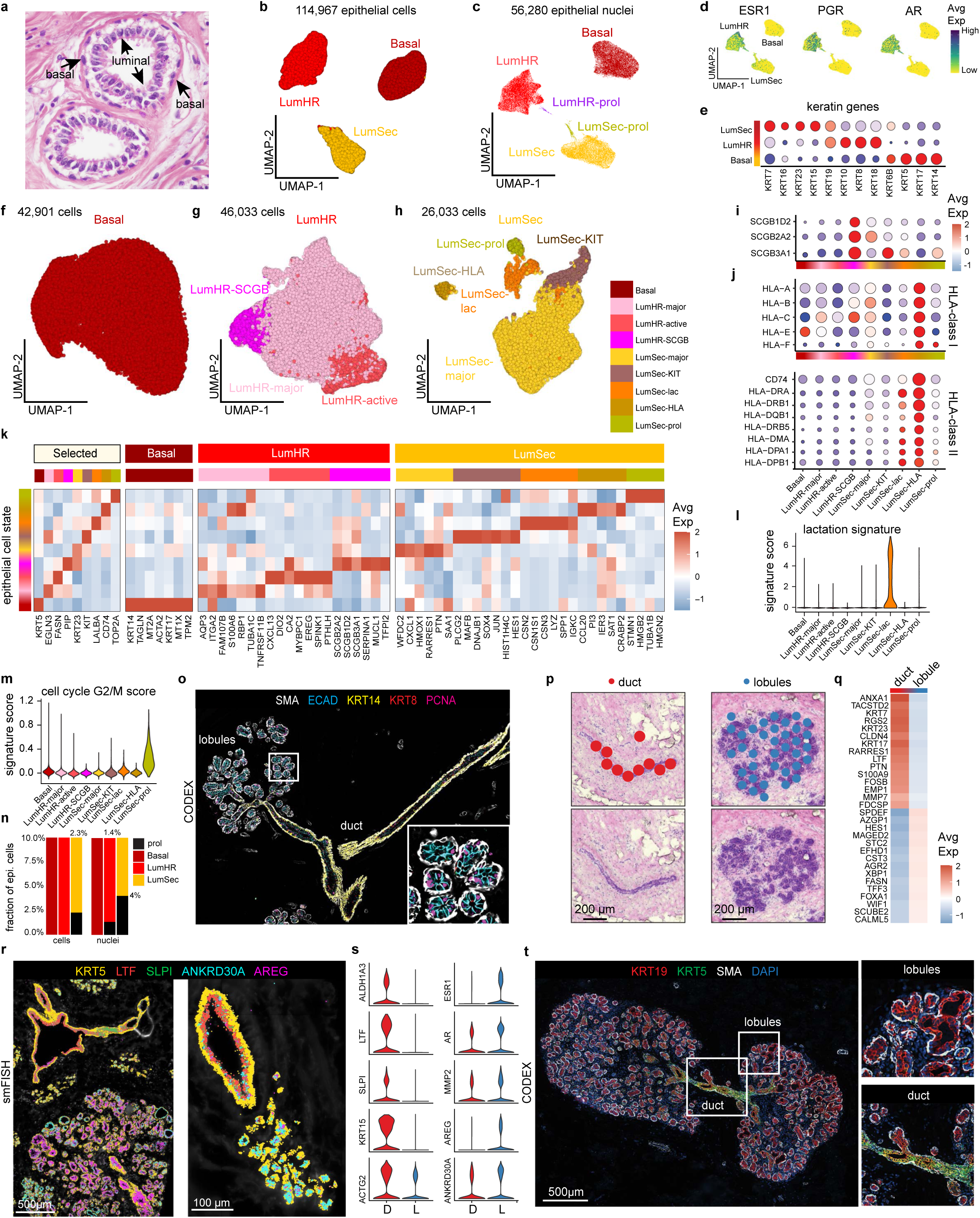
Epithelial cells of the Human Breast. **a,** Histopathological section of breast tissue showing the epithelial bilayer of a duct. **b,** UMAP of 114,967 epithelial cells, showing three major epithelial types. **c,** UMAP of 56,280 epithelial nuclei, showing 3 major epithelial types and two clusters of proliferating cells. **d,** UMAPs of snRNA-seq data showing the expression of hormone receptor genes. **e,** Dot plot of Keratin genes expressed across the 3 major epithelial cell types. **f,** UMAP of 42,901 basal epithelial cells. **g,** UMAP of 46,033 Luminal Hormone Responsive (LumHR) epithelial cells, showing 3 cell states. **h,** UMAP of 26,033 Luminal Secretory (LumSec) epithelial cells, showing 6 cell states. **i,** Expression of secretoglobin genes across the epithelial cell states. **j,** Expression of HLA-class I and HLA-Class II genes for the epithelial cell states **k,** Heatmap of top genes expressed for each epithelial cell state averaged across the scRNA-seq data. **l**, Lactation gene signature scores for the epithelial cell states. **m,** G2/M cell cycle scores across different epithelial cell states. **n,**Stacked barplots showing the fraction of proliferating epithelial cells in the scRNA-seq and snRNA-seq data among all epithelial cells. **o,** CODEX data from P66 showing proliferating cells in ducts and lobules labelled with PCNA. **p,** ST clusters for ductal and lobular regions labelled in H&E images **q,** Heatmap of ST top differentially expression genes between ducts versus lobules combined from 4 tissue samples. **r,** smFISH data (P46-S1 and P46-S4) showing genes that are expressed specifically in ductal and lobular regions. **s,** smFISH data showing *in situ* validation of genes that are expressed uniquely in the ducts and lobules in the scRNA-seq and ST data. **t**, CODEX data from P67 showing KRT5 in ducts and KRT19 in lobules/TDLU regions, with enlarged panels of the right.

To further resolve epithelial diversity, we clustered the scRNA-seq data from each epithelial cell type (Fig. 3f-h, k; Extended Data Fig. 4a). This analysis revealed that basal cells were remarkably homogenous, expressing *ACTA2* and *TP63*, as well as low levels of *EPCAM*, consistent with their role in basement membrane production and myoepithelial functions (Fig. 3f, k, Extended Data Fig. 4b,d). In contrast, a larger number of known and previously unrealized cell states were identified in the luminal cells (Fig. 3g,k). Within LumHR cells, three distinct cell states were identified, including one major cluster representing the canonical hormone responsive cell state (LumHR-major) marked by *EGLN3* expression, and two smaller clusters, one marked by secreted factors such as *MUCL1,* Prolactin-inducible protein *(PIP),* and secretoglobins (LumHR-SCGB), and the other cluster marked by genes downstream of hormone signaling such as *FASN*^28^ (LumHR-active) (Fig. 3g,i,k, Extended Data Fig. 4e). The LumHR-active cluster also displayed expression of genes that may relay pro-proliferative signals on neighboring epithelial cells upon hormone sensing, such as epiregulin (*EREG*)^29^ (Fig. 3g,i,k).

The highest cell state diversity was detected within LumSec cells, harboring five distinct cell states (Fig. 3h). The major canonical cell state marked by *KRT23* expression (LumSec major) in addition to four smaller cell states (Fig. 3h,k Extended Data Fig. 4f). The LumSec HLA cluster expressed genes encoding MHC-I and MHC-II molecules, as well as the chemokine *CCL20*, suggesting a role in immune cell signaling (Fig. 3j,k, Extended Data Fig. 4f). The LumSec-lac cell state was marked by genes involved in lactation such as caseins (*CSN2*, *CSN1S1*) and showed a higher lactation signature score (Fig. 3l, Extended Data Fig. 4f). The LumSec-prol cell state was characterized by genes involved in cell cycle and proliferation and showed an elevated G2/M score (Fig. 3m, Extended Data Fig. 4f). The LumSec-KIT cell state showed expression of the proto-oncogenes *KIT* and *JUN* as well as transcription factors including *SOX4*, *HES1* and *MAFB* (Fig. 3k, Extended Data Fig. 4f). Altogether, nine epithelial cell states were discovered, which varied across the 62 women (Extended Data Fig. 4c, Supplemental Table 10).

Interestingly, epithelial cell proliferation was restricted to LumSec and LumHR clusters, whereas basal cell proliferation was not detected. Small percentages of proliferating cells were identified in the scRNA-seq and snRNA-seq data within LumSec (2.3% of cells, 4% of nuclei) and LumHR (1.4% of nuclei) clusters (Fig. 3c,n). Consistent with S-phase activity, high levels of PCNA mRNA and protein expression were detected in luminal cells (Fig. 3o, Extended Data Fig. 4f). Consistently, the S-phase signature scoring revealed highest S-phase scores within the LumSec-prol cluster in scRNA-seq data, and within LumSec-prol and LumHR-prol clusters in snRNA-seq data, respectively (Extended Data Fig. 4g,h). Luminal cell proliferation occurred in both ductal and lobular regions, as revealed by *in situ* anti-PCNA immunofluorescence and *MKI67* transcript detection using smFISH (Fig. 3o, Extended Data Fig. 5d). Altogether, this data indicates that most proliferation occurs within the luminal compartment of the breast epithelium.

Next, we compared ductal and lobular regions by leveraging three spatial technologies. Using ST, we performed DE analysis between histopathological breast regions annotated as ducts and lobules/TDLUs (Fig. 3p, Extended Data Fig. 5a). This analysis identified many genes enriched in ducts that were associated with the LumSec cell type (e.g. *KRT17*, *LTF*, *KRT23*), while lobules showed increased expression of LumHR specific genes (e.g. *AZGP1*, *AGR2*) (Fig. 3q). In line with these findings, smFISH revealed increased expression of LumSec related genes in the ducts (e.g. *LTF, SLPI, KRT15*), while lobules were enriched for LumHR associated genes (*ANKRD30A, AR, ESR1*, *PGR*) (Fig. 3r,s, Extended Data Fig. 5c,d). On protein level, we found elevated levels of *KRT5* and *KRT14* in ducts versus lobules using CODEX (Fig. 3t). *KRT5* staining patterns were of particular interest, since *KRT5* is considered a basal cell marker, yet a significant enrichment of *KRT5+* cells was also found within the KRT19-positive luminal cell layer in ducts (Fig. 3t). Presence of KRT5/KRT19-double positive cells in ducts indicated an enrichment of cells with an intermediate basal-luminal expression profile within ductal structures, which have been linked to loss of lineage fidelity and cancer initiation^30^. Previous studies referred to the two luminal cell types (LumHR and LumSec) as alveolar and ductal^14, 16^. However, our analysis indicates that while the abundance of LumHR and LumSec cells differs between ducts and lobules, both cell types exist within ducts and alveolar structures of lobules (Fig. 2h, m, Extended Data Fig. 5b,e,f).

### Immune Ecosystem of the Normal Breast

The scRNA-seq data of 63,815 cells and 17,289 nuclei from 62 women showed that immune cells were organized into three major populations (myeloid, NK/T and B cells) and were unexpectedly abundant (11.9% total cells, 13.9% total nuclei) in all three different breast tissue sources (CM, PM, RM) (Fig. 4a,b). The abundance of the immune cells was also validated in the spatial CODEX and smFISH data (Fig. 4c-e), as well as our histological images (Extended Data Fig 7a). To further investigate immune cell diversity, we clustered cells within the myeloid, NK/T and B cell clusters and annotated the cell states using both unbiased and canonical marker genes^31–33^ (Fig.4f-k, Extended Data Fig. 6a-c).

**Figure 4.**
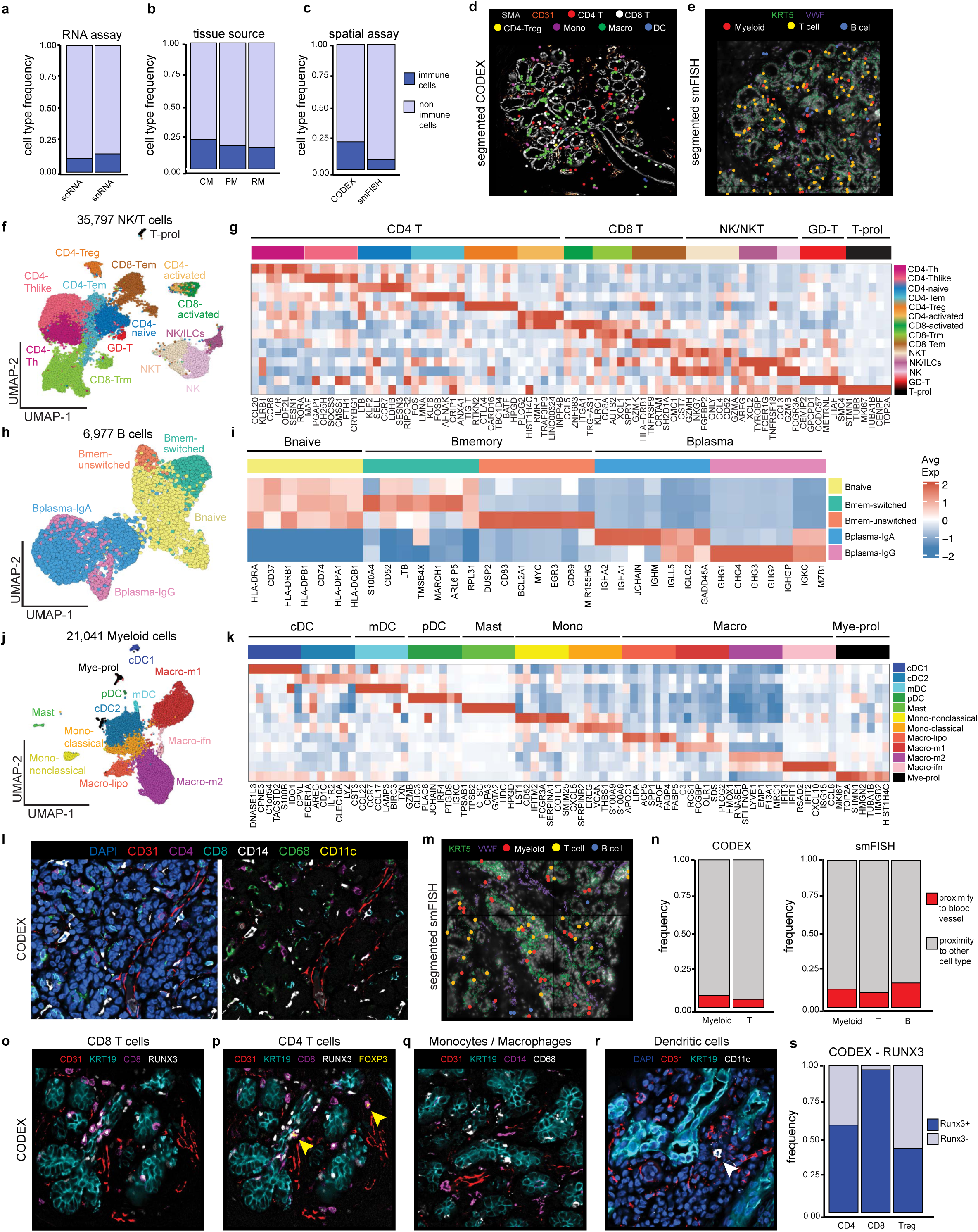
Immune Cell Abundance and Diversity in the Human Breast. **a,** immune and non-immune cell type frequencies among total cells in the scRNA and snRNA- seq data **b**, immune and non-immune cell type frequencies by tissue source in the scRNA-seq data. **c,** immune cell type frequencies quantified from the CODEX N=4 and Resolve smFISH N=12 tissue samples. **d,** CODEX data from P66 showing TDLU region with localization of 6 immune cell types, where segmented cells are shown as colored dots overlaid on image of IF staining of SMA (myoepithelial) and CD31 (vessel) for spatial reference. **e,** smFISH data (P46- S1) showing TDLU region with localization of 3 immune cell types, where segmented cells are shown as colored dots overlaid on image of IF staining of KRT5 (basal epithelial) and VWF (vessel) for spatial reference. **f,** UMAP of scRNA-seq data showing clustering of 35,797 NK/T cells into 14 cell states. **g,** heatmap showing top genes expressed for each NK/T cell cluster using average values across single cells. **h,** UMAP of 6,977 B cells scRNA-seq data showing five cell states. **i,** heatmap showing top genes expressed for each B cell state using average values across single cells. **j,** UMAP of 21,041 myeloid cell scRNA-seq data showing clustering of 12 cell types and states. **k,** heatmap showing top genes expressed for each myeloid cell cluster using averaged scRNA-seq values. **l,** CODEX data from P66 showing co-localization of immune cells and vascular marker (CD31). **m,** smFISH segmented data (P46-S1) showing co localization of immune cells and vascular marker (VWF). **n,** frequency of immune cells that are in the proximity of vessel markers versus other cell type markers by neighborhood analysis in CODEX and smFISH data. o-r, CODEX data from P66 and P67 showing co-localization of different immune cells with epithelial marker KRT19 and vascular marker CD31. Yellow and white arrows indicate CD4 Tregs and DCs, respectively. **s,** frequency of T cells expressing Runx3 tissue residency marker in CODEX data.

Within the NK/T cells, the scRNA-seq data of 35,797 cells identified 14 subsets (Fig 4f,g, Extended Data Fig 6a). The CD4 T cells included naïve T cells (*SELL*), Th and Thlike (*IL7R, CCR6, CCL20*), T effector memory (Tem (*LMNA*) and T regulatory (Treg) (*FOXP3*) cells. We identified two populations of CD8 T cells, T resident memory (Trm) (*ITGA1*) and Tem (*GZMK*), as well as a cluster of gamma delta (GD) T (*TRGC1*) cells. We also detected clusters of CD4 and CD8 T cells that displayed upregulation of genes associated with TCR activation (*PLCG2, ZNF683*). The subclustering further resolved populations of natural killer (NK), NKT, and innate lymphoid cells (ILC) that were discriminated by their combinatorial expression of NK (*GNLY*), T cell (*CD3D*), and ILC (*IL7R*) markers. We detected a small proliferating cluster (*MKI67*) that contained cells from many NK and T cell subsets. Notably, most subsets expressed the tissue resident marker *RUNX3*^34^ and did not express high levels of checkpoint or exhaustion markers^35^, which is consistent with a homeostatic rather than disease-associated phenotype (Extended Data Fig. 6d). Clustering of the 6,977 integrated scRNA-seq data of the B cells showed populations of naïve, memory, and plasma B cells, which were distinguished by their expression of *CD27* (memory), *CD38* (plasma), or neither (naïve) (Fig. 4h,i, Extended Data Fig. 6b). Among the memory B cells, we identified both switched (*MYC*) and unswitched (no *MYC* expression) cells. We also identified two populations of plasma B cells that separated based on their preferential expression of IgG or IgA immunoglobulins.

Clustering of the 21,041 scRNA-seq myeloid cells resolved distinct subsets of dendritic cells (DC), monocytes, macrophages, and mast cells (Fig. 4j,k, Extended Data Fig. 6c). We identified four subpopulations of dendritic cells, including mature (mDC) (*LAMP3*), plasmacytoid (pDC) (*LILRA4*), and two conventional cell states (cDC1 (*CLEC9A*), and cDC2 (*CLEC10A*)). Among the macrophages, we detected classically (Macro-m1, *IL1B*) and alternatively (Macro-m2, *MRC1*) activated subsets, a population defined by the expression of interferon response genes (Macro-ifn, *IFIT1*, *IFIT2*), and a lipid-associated macrophage subcluster (Macro-lipo, *APOC, LPL, TREM2*). We also identified populations of classical (*EREG*) and non-classical (*FCGR3A*) monocytes, as well as mast cells defined by their expression of *TPSAB1*. This analysis also identified a heterogenous cluster of proliferating myeloid cells (Mye-prol, *MKI67*). Although the NK/T, B and myeloid cell states showed variable frequencies, they were consistently identified across the 50 women, suggesting they play a ubiquitous function in normal breast homeostasis (Extended Data Fig. 6e-h).

The spatial organization of seven major immune cell types (monocytes, macrophages, CD4 T, CD8 T, CD4 Treg, dendritic and B cells) was delineated using a total of 18 markers on the smFISH platform and 8 markers on the CODEX (Extended Data Fig. 7b,c, Supplementary Table 6, 9, respectively). Spatial analyses showed that all seven cell types were present in three major tissue regions (C, D, L), excluding adipose which could not be assessed (Extended Data Fig. 7c-f). The highest density of immune cells was observed in the epithelial D and L regions, whereas the connective tissue showed only very sparse immune cell density (Extended Data Fig. 7c-f). Importantly, most immune cells were found in the breast tissue parenchyma rather than in vessels, consistent with a tissue resident phenotype (Fig. 4l-n). Furthermore, many immune cells were embedded within the epithelial layers and had elevated levels of the *RUNX3* protein, further indicative of a tissue resident phenotype^34^ (Fig. 4o,p,s). Collectively, these data show that normal breast tissues harbor an abundant and diverse milieu of immune cell types.

### Fibroblast Cell Diversity in the Breast

Fibroblasts are mesenchymal cells responsible for the production of the extracellular matrix (ECM), which supports both the epithelial structures, as well as the connective tissue of the breast^1^. In our data, fibroblasts represented an abundant cell type in the breast (21.1% of cells, 16.5% of nuclei). Previous histopathological studies have annotated breast fibroblasts as ‘intralobular’ or ‘interlobular’ based on their tissue localization (Fig. 5a)^1, 2^. We reclustered scRNA-seq data from 113,157 fibroblast cells across 62 women, which identified three distinct fibroblast cell states (Fig. 5b,c, Extended Data Fig. 8a-b). Unbiased signature analysis suggested that two cell states were enriched for integrin, ECM, and collagen gene signatures, while the third had signatures of antioxidant and chemokine activity (Fig. 5d). The ‘fibro-matrix’ cells showed high transcription of collagen genes and scored highest for a ‘collagen’ gene signature (Fig. 5d-e, Extended Data Fig. 8c,d). The ‘Fibro-prematrix’ cells displayed elevated pre-collagen genes such as *PCOLCE2*, and genes involved in fatty acid metabolism (*FABP4, CD36, PPARG*). The ‘Fibro-immune’ cell state displayed elevated expression of cytokines and chemokines and therefore likely plays a role in immune cell regulation (Fig. 5c). We further investigated the expression of *FAP,* an activated fibroblast marker often reported in cancer associated fibroblasts (CAFs), which showed very low expression in the three cell states (Fig. 5f).

**Figure 5.**
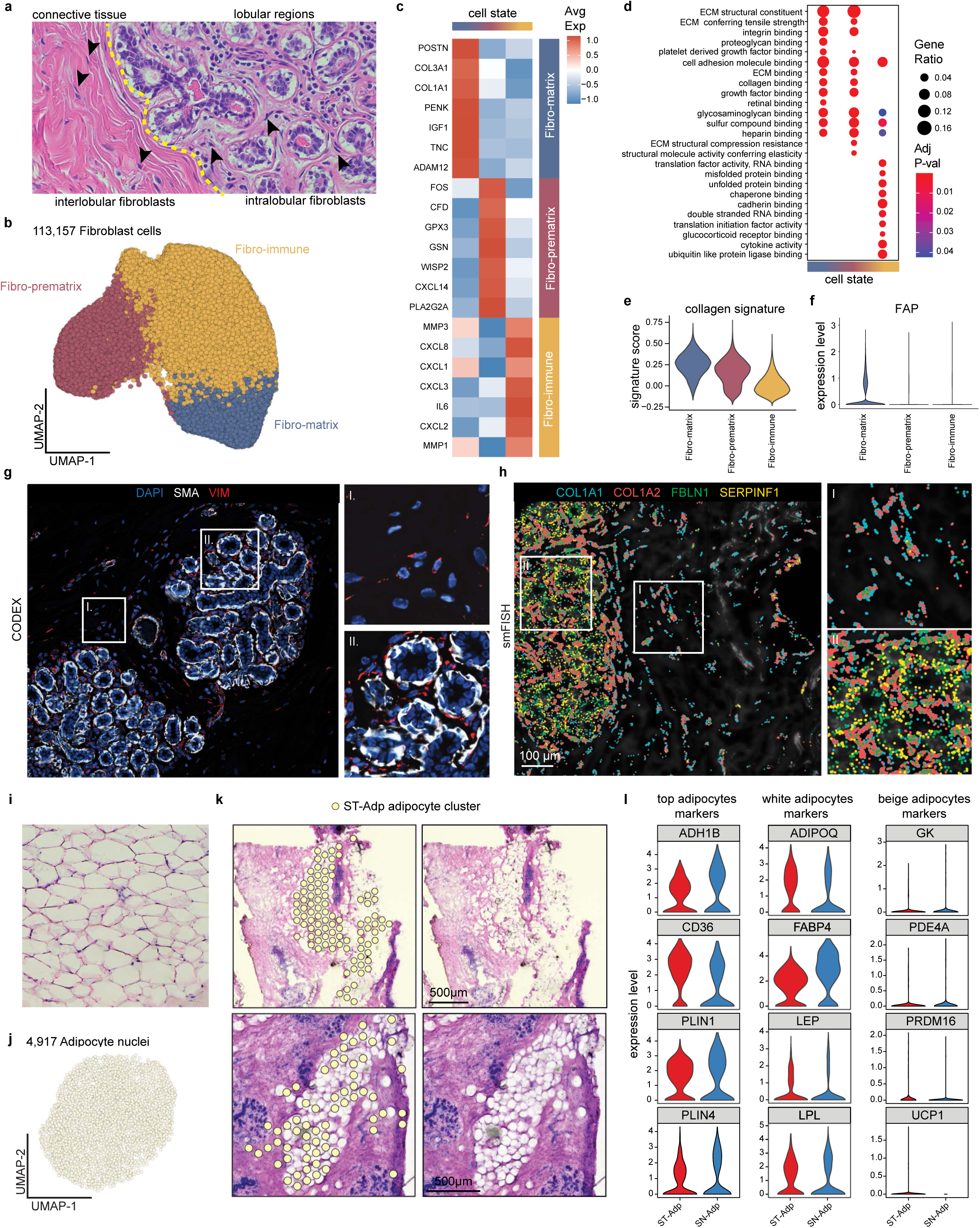
Breast Fibroblasts and Adipocytes. **a,** Histopathological sections showing regions with intralobular and interlobular fibroblasts in the breast. **b,** UMAP of 113,157 fibroblast cells, showing 3 major cell states. **c,** Heatmap of top genes expressed for each fibroblast cell state, averaged from the scRNA-seq data. **d,** Unbiased gene set enrichment analysis of the top pathways and signatures associated with each cell state. **e,** Collagen gene signature scores of the fibroblast cell states. **f**, expression of *FAP* across the fibroblast cell states. **g,** CODEX data from P68 showing fibroblasts marked by VIM in the connective tissue (I) and interlobular (II) regions. **h,** smFISH data (P64-S3) showing a subset of four fibroblast genes and their distribution in the connective tissue (I) and intralobular (II) areas. **i,** Histopathological section of breast adipose tissue. **j,** UMAP of 4,917 adipocyte snRNA-seq data. **k,** ST showing adipocyte clusters identified and matching H&E images from two women (P10 and P46). **l,** Expression of top adipocyte marker genes in the ST data and snRNA data (left panels), top white adipocyte markers (middle panels), and top beige adipocyte markers (right panels).

To investigate the spatial distribution of the fibroblasts, we first used the pan-fibroblast marker vimentin (VIM) in our CODEX analysis, which revealed two prominent locations of fibroblasts in the interlobular and intralobular regions (Fig. 5g). To further distinguish the three cell states, we utilized four genes from our spatial smFISH panel, including *COL1A1* and *COL1A2* (Fibro matrix), as well as *FBLN1* and *SERPINF1* (Fibro-prematrix). This data showed that the pre matrix markers were expressed predominantly in the lobular regions (Fig. 5h, Extended Data Fig. 8f). To further quantify this finding, we classified the smFISH genes into three groups (epi proximal, epi-middle and epi-distal), which confirmed that *SERPINF1* and *FBLN1* was elevated in the intralobular regions (Extended Data Fig. 8f,g,h). We also used RNAscope to investigate the expression of *MMP3* in the Fibro-immune cell state, which showed higher levels in the fibroblasts proximal to lobular epithelial regions, consistent with the smFISH data (Extended Data Fig. 8i, Methods).

### Adipose Tissues of the Breast

Adipose tissue represents a large proportion of the human breast and plays important roles as a source of energy and hormones^12, 22, 36^. Adipocytes are the main cell type in adipose tissues and are readily identified by histopathology (Fig. 5i). However, adipocytes have been notoriously difficult to profile with single-cell genomics methods due to their large cell size, high lipid content and fragile nature^6^. Indeed, adipocytes were not captured by our scRNA-seq methods (Fig. 1c). Therefore, we leveraged snRNA-seq to capture the transcriptomic profiles of 4,917 breast adipocytes (4.1% of nuclei) and spatial transcriptomics (ST) data from 4 women (Fig 5j,k). Collectively, these methods identified *ADH1B*, *CD36*, *PLIN1*, *PLIN4*, *ADIPOQ*, *FABP*4, *LEP* and *LPL* as the top genes expressed in breast adipocytes, which was consistent across the two orthogonal platforms (Fig 5l). Both the snRNA-seq and ST data showed that most adipocyte markers were expressed uniformly across all adipocytes, with limited cell state heterogeneity. We further interrogated our dataset for brown/beige and white adipocyte markers, which showed that breast adipocytes exclusively corresponded to white adipocytes (Fig. 5l).

### Vascular and Lymphatic Endothelial Cells

The human breast is a highly vascularized organ containing a network of veins and arteries that diffuse into the ducts and lobules via capillaries and are often detected in histopathological sections (Fig. 6a). Our scRNA-seq analysis of 40,824 endothelial cells across 62 women showed that vascular endothelial cells (expressing *PECAM1* and *VWF)* represent an abundant cell type (7.6% of cells, 7.3% of nuclei) in the normal breast (Fig. 6b). Re-clustering of scRNA-seq vascular endothelial cells identified 3 major cell states that corresponded to arterial endothelial (*SOX17*, *GJA4*), venous endothelial (*ACKR1*, *SELP*) and capillary endothelial (*RGCC*, *CA4*) cells based on canonical markers^37, 38^, and further identified many new top marker genes (Fig. 6b,c, Extended Data Fig. 9b,c). In addition to the vascular system, the lymphatic network represents a ‘passive system’ for removing cellular waste and is often identified in histopathological sections (Fig. 6d). Our data shows that lymphatic endothelial cells (expressing *PROX1* and *PDPN)* occur at low frequencies (1.4% of cells, 3.9% of nuclei) in breast tissue. Clustering of 7,530 lymphatic cells from the scRNA-seq data identified three major cell states that varied across women (Fig. 6e,f, Extended Data Fig. 9e). The Lym-major cell state expressed *LYVE1* and *CCL21* and represented the most abundant component of the lymphatic vessels (Extended Data Fig. 9f). The Lym-immune cell state expressed *ACKR4* and *NTS* which resembles cells on the ceiling of subcapsular sinus in human lymph nodes^38^, as well as several chemokine ligands and signatures, suggesting a role in immune cell signaling (Extended Data Fig. 9g). The third cell states are the Lym-valve cells which expressed *CLDN11* and play an important role in preventing lymphatic fluid backflow^39^.

**Figure 6.**
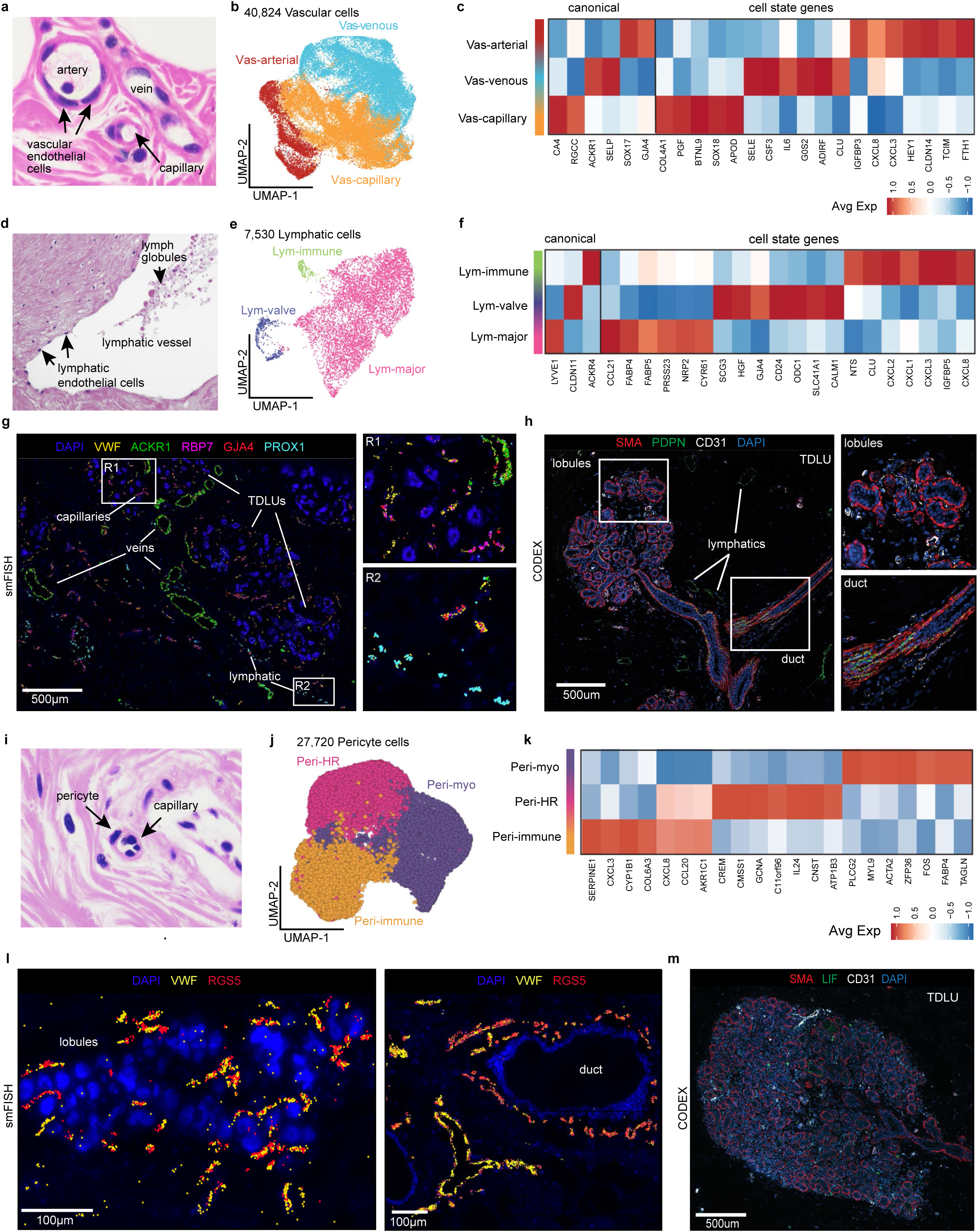
Vascular, Lymphatic and Pericyte Cells in the Human Breast. **a,** Histopathological section showing veins and capillaries in breast tissue. **b,** UMAP of 40,824 vascular endothelial cells, showing 3 major cell states. **c,** Heatmap of top genes expressed for each vascular endothelial cell state, using averaged values from scRNA-seq data. **d,** Histopathological section showing a lymphatic duct in breast tissue. **e,** UMAP of 7,530 lymphatic endothelial cells, showing 3 major cell states. **f,** Heatmap showing expression of top genes for each lymphatic cell state, averaged from the scRNA-seq data. **g,** smFISH data (P47-S1) showing a subset of vascular gene markers (*VWF*, *ACKR1*, *RBP7* and *GJA4*) and lymphatic markers (*PROX1*), with enlarged right panels showing vascular (R1) and lymphatic cell (R2) regions **h,** CODEX data from P66 of a TDLU region showing vascular cells (anti-CD31) and lymphatic cells (anti-PDPN) cells, with myoepthelial cells labelled (anti-SMA). The two right panels show zoomed-in regions with vascular cells near lobules and a duct. **i,** Illustration of pericytes and histopathological section showing capillaries with pericytes attached. **j,** UMAP projection and clustering of 27,720 pericyte cells, showing 3 cell states. **k,** Heatmap of top genes expressed for each pericyte cell state from averaged scRNA-seq data. **l,** smFISH data showing vascular cells (*VWF*) and pericyte (*RGS5*) labelled in two different breast tissue samples (P46-S3 and P35-S1). **m,** CODEX results from P67 showing vascular cells (anti-CD31) and pericytes (anti-LIF) in a TDLU region.

Using the three spatial platforms, we investigated the localization of the vascular and lymphatic cell states. The ST data showed two distinct clusters for vascular and lymphatic cells that corresponded to the histopathological vessel structures and validated many scRNA-seq cell type markers *in situ* (Extended Data Fig. 9h). The smFISH data showed that larger venous structures expressing *ACKR1* and the vascular marker *VWF* are typically found in the connective tissues, while smaller capillary structures expressing *RBP7* were closely integrated within epithelial lobular or ductal structures (Fig. 6g, R1 inset panel, Extended Data Fig. 9i). Spatial analysis using smFISH showed that the lymphatic cells (*PROX1* positive) are located predominantly in connective tissues regions (Fig. 6g). This was also reflected in the CODEX analysis using anti-*PDPN* (Fig. 6h). Collectively, these data provide a molecular definition of the vascular and lymphatic cell states in the human breast.

### Pericytes of the Breast

Pericytes have traditionally been described as ‘vascular accessory cells’ that play a role in regulating vasoconstriction in capillaries to regulate blood-flow into tissues^40, 41^ (Fig. 6i). While pericytes have been studied extensively in the brain^28^, there is limited data on their role in normal breast tissues. Unexpectedly, our scRNAseq data from 27,720 pericytes across 62 women showed that pericytes are an abundant cell type (5.2% of cells, 2.1% of nuclei) in the human breast. Clustering of the scRNA-seq pericyte data identified 3 major cell states: myopericytes (Peri-myo), immun-pericytes (Peri-immune) and hormone-responsive pericytes (Peri-HR) (Fig. 6j-k, Extended Data Fig.10a,b). These pericyte cell states were present at variable frequencies in most of the women in our study (Extended Data Fig. 10d). The Peri-myo were consistent with their classical role in constricting blood vessels to regulate blood flow^41^ (Fig. 6k, Extended Data Fig. 10c). The Peri-immune expressed gene signatures involved in chemokine and cytokine functions, suggesting they are involved in immune cell signaling (Extended Data Fig.10c). The Peri-HR cells expressed genes such as *CREM*, *CMSS2* and *GCNA*, and showed gene signatures associated with hormone and gluococorticoid receptor binding (Extended Data Fig. 10c). Spatial analysis using smFISH showed that pericytes (*RGS5* positive) were highly abundant in normal breast tissues in lobular regions, where they often co-localized with vascular cells (*VWF* positive) (Fig. 6l, Extended Data Fig. 10g-i). Similarly, CODEX proteomic analysis showed that pericytes (LIF-positive) were often located in lobular regions and co-localized with vascular markers (CD31-positive) (Fig. 6m). This data suggests that large numbers of pericytes congregate in lobular regions and may have diverse functions in breast tissues.

### Metadata Correlations with Cell Types and States

We investigated the association of the breast cell type and state frequencies with clinical parameters, including age, menopausal status, body mass index (BMI) and parity (pregnancy) status (Extended Data Fig. 11). Menopause status and older age (>50 years) both correlated with lower frequencies of cell types including the basal epithelial (p = 0.004, p = 0.019, Fisher Exact) and B cells (p = 0.002, p = 0.003, Fisher Exact) and decreased LumSec epithelial cells (p= 0.049 in menopause, N.S. with old age, Fisher Exact) (Extended Data Fig 11a,c). Menopause and old age were also associated with many cell state changes, including the epithelial cell states (LumHR-active, LumSec-major, LumSec-KIT, LumSec-lac, LumSec-prol), a fibroblast cell state (Fibro-matrix) and many immune cell state changes, including B cells (Bnaive, Bmem-switched, Bplasma-IgA), T cells (CD4 naïve, CD4 Th1, CD4, Trm, CD8-Trm) and myeloid cells (Macro m1) (Extended Data Fig 11b). Additionally, this analysis revealed significant associations between obesity (overweight: > 25 BMI) and decreased lymphatic endothelial cells, as well as epithelial cell state changes (LumSec-KIT) and T cell changes (CD8-ZNF683) (Extended Data Fig 11e,f). Notably, this analysis did not identify any significant cell type changes associated with parity status, however, it identified significant associations with parity and increased CD4- Tregs (Extended Data Fig 11g-h).

## Discussion

Here, we report an unbiased atlas of the adult human breast tissues that comprises 11 major cell types and 53 unique cell states that are organized into 4 major spatial tissue domains (Supplementary Fig. 12). In the epithelial cells, our data show limited basal cell state diversity, while the two luminal epithelial cell types comprise 7 distinct cell states reflecting diverse biological functions. Our data estimate that only 2-4% of the breast epithelial cells are proliferative, and most are associated with the LumSec populations consistent with their previous annotation as ‘luminal progenitors’. However, we also identified a small number (1.4%) of proliferating cells in the LumHR population snRNA-seq data. Notably, no proliferating cells were detected in the basal cells, which may prompt the field to revisit the concept of a basal stem cell fueling epithelial homeostasis^10^. Our detailed spatial comparison between epithelial ducts and lobules provided insights including the presence of luminal cells with basal-like features (e.g., KRT5 expression) as well as increased LumSec cells within ducts, while lobules and TDLUs were enriched in LumHR cells.

In the non-epithelial compartment, we identified an unexpectedly abundant (∼12%) and diverse milieu of tissue-resident immune cells in the normal breast tissues. Our spatial analysis shows that both lobules and ducts are immune-rich ecosystems, where many of the T cells, B cells and myeloid cells congregate. Only a small number of immune cells overlapped with vascular structures, and most express the tissue resident marker *RUNX3*. Understanding the variation and diversity of the immune cells is very important for breast cancer, where immunotherapy has recently become the standard of care for some subtypes^42^. Additionally, our data shows that pericytes are an abundant cell type that is integrated in the lobular epithelium and may have additional biological functions beyond their classical role in vascular constriction^40, 41^. The genomic references of lymphatic endothelial cells are of particular relevance, due to their wide use in the clinical evaluation in lymph-node positive breast cancers^43^. Additionally, the snRNA-seq and ST provide one of the first genomic references of the genes expressed in breast adipocytes and show that they are exclusively white adipocytes^44^.

Our comprehensive metadata identified significant changes in the breast tissue architecture that corresponded to menopause, age, and BMI, consistent with some previous pathological studies^45–47^. However, we did not find any significant associations with parity status, where previous studies have suggested differences in the epithelial populations^14, 48, 49^. This may reflect that our cohort did not include any pregnant or lactating women, which represents an important future direction for building this atlas^15^. A notable drawback of our HBCA is the lack of ethnic and ancestral diversity, which mainly included Caucasian and African American women (Supplementary Fig 11i). This bias should also be addressed in future studies to advance our understanding of diseases and improve outcomes for women of all backgrounds. In closing, this human breast cell atlas significantly advances our knowledge of the epithelial and non epithelial cell types in adult human breast tissues, providing a comprehensive reference for studying mammary biology, development, and breast cancer.

## Data Availability

HBCA website: http://www.breastatlas.org

## Software Availability

Code is available under: https://github.com/navinlabcode/HumanBreastCellAtlas

## Author Contributions

Data analysis was performed by TK, RW, SH, KB, MP, YG, MH, AC, BN, NK, KC. Experiments were performed by KN, QN, SB, TK, ES, JL, AT, MP, TR. Tissue samples and clinical coordination was performed by OB, BBC, AW, JW, HC, AC, MT, BM, HJ, JM, RE, DT, CN, EL, RP, SW, AT, BL. Project management and manuscript writing was performed by BL, DL, NN and KK.

## Acknowledgements

This work was supported by major funding support from the Chan-Zuckerberg Initiative (CZI) SEED Network Grant (CZF2019-002432). This work was supported by grants to N.E.N. from the NIH National Cancer Institute (RO1CA240526, RO1CA236864, 1R01CA234496, F30CA243419), the CPRIT Single Cell Genomics Center (RP180684), the American Cancer Society (132551-RSG-18-194-01-DDC). N.E.N. is an AAAS Fellow, AAAS Wachtel Scholar, Damon-Runyon Rachleff Innovator, Andrew Sabin Fellow, and Jack & Beverly Randall Innovator. TK is funded by the NCI T32 Translational Genomics Fellowship and Rosalie B. Hite fellowship. This study was supported by the MD Anderson Sequencing Core Facility Grant (CA016672). MP is supported by a fellowship from the CIRM Training Grant (EDUC4-12822). The work was also supported by a Damon-Runyon Quantitative Biology Postdoctoral Fellow to R.W. We thank Bailey Marshall, Norbert Tavares and Jonah Cool at CZI for their guidance and support. We thank Jill Waters, Sam Stingley and Lewis Vann from Resolve Biosciences for their support on this project. We are grateful to Jie Wiley, Anita Wood, Angela Alexander and Alejandro Contreras for clinical support. We are thankful to Aaron Longworth, Sharmila Mallya and Michael Curran for their advice. We thank Yiyun Lin and Rui Ye at MD Anderson for help with experiments. Finally, we are extremely grateful to all of the women who participated in the HBCA and donated their breast tissue to this project. This publication is part of the Human Cell Atlas (www.humancellatlas.org/publications).

## EXPERIMENTAL METHODS

### Protocol Availability

The breast tissue dissociation protocols for preparing cell suspensions and nuclear suspensions that were developed for the HBCA project have been deposited to protocols.IO (http://www.protocols.io) under the following accessions:

*Dissociation of viable cell suspensions from human breast tissues* https://www.protocols.io/view/dissociation-of-single-cell-suspensions-from-human-bp2l641bkvqe/v1

*Dissociation of nuclear suspensions from human breast tissues* https://www.protocols.io/view/dissociation-of-nuclear-suspensions-from-human-bre-x54v98ym4l3e/v1

### Collection of Normal Breast Tissue Samples

Fresh breast tissue samples were collected from the University of California, Irvine, Baylor College of Medicine, St. Luke’s Medical Center and the Cooperative Human Tissue Network (CHTN). The study was approved by the Institutional Review Boards at the respective institutions using mirror protocols, including MD Anderson Cancer Center (PA17-0503), Baylor College of Medicine (H-46622), UC Irvine (HS-2017-3552). Reduction mammoplasty tissues were collected mainly at Baylor St. Luke’s Medical Center, while prophylactic mastectomies and contralateral mastectomies from the other breast of cancer patients were collected at MD Anderson and UC Irvine. With the exception of the CHTN samples, all of the fresh breast tissue samples were collected 1-2 hours after the surgical procedures and dissociated into viable cell suspensions using 1 hour, 6 hour or 24 hour dissociation protocols. All of the tissue samples at the respective institutions were analyzed for normal pathology at the time of collection and any women within incidental tissues with pre-cancer diagnosis (eg. ADH or DCIS) were excluded.

### Breast tissue dissociation for single-cell RNAseq

Detailed protocol for overnight breast tissue mechanical and enzymatic digestion for single-cell RNA sequencing (scRNAseq) were developed and optimized for the HBCA project and can be found with step-by-step instructions at Protocols.io (www.protocols.io). Surgical tissue was transported in sterile DMEM medium (Sigma #D5796) on ice. Excess adipose tissue was removed prior to dissociation. Large breast tissue pieces are divided into individual 1-2g preparations, which will be subjected to dissociation solution consisting of collagenase A (1mg/ml working solution, Sigma #11088793001) dissolved in DMEM F12/HEPES media (Gibco #113300) and BSA fraction V solutions (Gibco# 15260037) mixed at a 3:1 ratio, respectively or, 20ml of 4mg/ml Collagenase Type 1 (in 5% FBS DMEM). For each preparation, a 10cm dish with 2ml dissociation solution was used to mince tissue into homogenous suspension with paste-like consistency. Minced tissue was transferred into a 50ml conical tube with 40ml of dissociation solution in a rotating hybridization oven for 2 to 6 hours at 37°C until completely digested. Cell suspension was centrifuged at 500g for 5 minutes and supernatant was removed. The pellet was resuspended in 5ml trypsin (Corning #25053CI) at room temperature and incubated in a rotating hybridization oven at 37°C for 5 minutes. Trypsin was neutralized with 10ml DMEM containing 10% heat inactivated fetal bovine serum (FBS) (Sigma #F0926). The solution was mixed by pipetting up and down, and then filtered through a 70μm strainer (Falcon # 352350). A sterile syringe plunger flange was used to grind the leftover unfiltered tissue and DMEM was used to wash the remaining single cells off the filter. The flow-through was centrifuged at 500g for 5 minutes and the supernatant was removed. Resulting pellet was nutated at room temperature for 10min in 20ml 1x MACS RBC lysis buffer (MACS #130-094- 183) to remove red blood cells (RBCs). To stop RBC lysis, 20ml DMEM was added and then centrifuged at 500g for 5min. The cell pellet was washed in 10ml of cold DMEM and centrifuged at 500g for 5 minutes. Pellet was then resuspended in cold PBS (Sigma #D8537) +0.04%BSA solution (Ambion #AM2616) and filtered through a 40μm flowmi (Bel-Art #h13680-0040). Trypan blue stained cells were counted in the Countess II FL automated cell counter (Thermo Fisher) and their concentration was adjusted to 700-1200 cells/µl.

For overnight digestions, after digestion the enzymatic tissue digestion mixture was centrifuged at 400g for 5mins. Supernatant was removed and the tissue pellet was washed with 50mls of PBS. Supernatant was removed and 2ml of 0.05% trypsin was used to break up tissues into single cell suspensions in a 15ml conical and placed into a 37C water bath. Dissociation was accelerated by pipetting with a p1000 set at 1ml, pipetting up and down 10 times every 2 minutes. 10mls of 10%FBS+DMEM was used to neutralize the enzymatic digestion and the sample was centrifuged for 5mins at 400g. The resulting pellet was resuspended in 100ul in 20 U/mL DNase I (Sigma-Aldrich, D4263-5VL) and incubated at 37C for 5mins to liberate cells from DNA. 10mls of 10% FBS+DMEM was added and the tissue was centrifuged for 400g for 5mins. The resulting single-cell suspension was passed through a 100um strainer filter. Cells were then stained for FACS using fluorescently labeled antibodies for CD31 (eBiosciences, 48- 0319-42), CD45 (eBiosciences, 48-9459-42), EpCAM (eBiosciences, 50-9326-42), CD49f (eBiosciences, 12-0495-82), SytoxBlue (Life Technologies, S34857). Only samples with at least 80% viability as assessed using SytoxBlue with FACS were included in this study. For scRNAseq, we excluded doublets, and dead cells (SytoxBlue+), for FACS isolation. Flow cytometry sorted cells were washed with 0.04% BSA in PBS and suspended at approximately 1000 cells/µL.

### Single cell RNA Sequencing

Single-cell suspensions were immediately subjected to scRNAseq using the Chromium platform (10X Genomics). Single cell capture, barcoding and library preparation were performed by following the 10X Genomics Single Cell Chromium 3’ protocols (V2: CG00052, V3: CG000183, V3.1: CG000204). The final libraries were sequenced on the Novoseq 6000 system S2-100 flowcell (Illumina). Data were processed using the CASAVA 1.8.1 pipeline (Illumina Inc.), and sequence reads were converted to FASTQ files and UMI read counts using the CellRanger software (10X Genomics).

### Single-nucleus RNA sequencing

Detailed protocol for overnight breast tissue mechanical isolation for single-nucleus RNA sequencing (snRNAseq) can be found at protocols.io (www.protocols.io). To isolate single nuclei, 0.5-1g fresh breast tissue were placed in a 10cm dish with 2ml lysis buffer. Nuclei lysis buffer consists of NST-DAPI buffer with 0.1U/μl RNase Inhibitor (NEB #M0314L)^1, 2^. Tissue was minced until tissue chunks are no longer visible. The suspension was filtered through a 40μm cell strainer (Falcon #352340). A sterile syringe plunger flange was used to gently grind the leftover tissue on the filter and then rinsed with 3ml of lysis buffer. The flow-through was transferred into 5ml DNA LoBind tubes and incubated on ice for 10min. The tube was centrifuged at 500xg for 5min at 4°C. The supernatant was removed, and nuclei were washed with 1ml cold lysis buffer and centrifuged again. The nuclei pellet was resuspended in 1% BSA (Sigma #SRE0036) in PBS supplemented with 0.2U/μl RNase Inhibitor. Nuclei were filtered through a 40μm Flowmi cell strainer, counted by hemocytometer under DAPI channel and concentration was adjusted to 700-1200 nuclei/µl. 10X Genomics RNA experiments proceeded immediately to avoid nuclei aggregation. Single cell capture, barcoding, library preparation and sequencing were the same as detailed above. For nuclei preparations sorted using flow cytometry, a 10ml dounce tissue homogenizer was placed on ice, and 40g of breast tissue was placed in a 10cm tissue culture dish on ice. Approximately 10g of tissue was minced into fine (∼2mm x ∼2mm) pieces, which was then added to the dounce homogenizer. 10ml of nuclear isolation buffer (400ul 1M Tris-HCL ph =7.5, 80ul 5M NaCL, 120 ul 1M MgCL_2_, 400ul 10% NP-40, 39mls DNase/RNase free sterile H_2_O) was pipetted over the tissue into the dounce homogenizer. Tissue was dounce-homogenized with the piston until running smoothly.

Homogenization was repeated until all 40g of tissue was digested. Nuclei suspension is then centrifuged at 500g for 5 mins at 4C, washed in 1%BSA in PBS, and is stained with Hoechst for flow cytometry-based sorting of high-quality nuclei to be subjected to snRNAseq.

### Spatial Transcriptomic Profiling of Normal Breast Tissues

Spatial Transcriptomics experiments were performed using the Visium Platform (10X Genomics), with the following modifications to the manufacturer’s protocols.

Fresh breast tissues from four patients were embedded in cryomolds with OCT compound (Fisher #NC9542860, #1437365) over dry ice. The tissue blocks were stored at −80°C in sealed bags. 12μm sections were sections in a cryomicrotome (Cryostar NX70, Thermo Scientific) with chuck and blade temperatures set at −17 °C and −15 °C, respectively. The tissue section was placed within the capture area of the Visium spatial slide (10X Genomics PN-1000184). Protocol was optimized for normal breast tissue according to manufacturer’s Tissue Optimization protocol (10X protocol #CG000238) and the slides were permeabilized for 12 minutes. Sectioned slides were fixed and stained as detailed by manufacturer (10X protocol #CG000160). Imaging was conducted on the Nikon Eclipse Ti2 system following imaging guidelines (10X protocol #CG000241). The final libraries were constructed by following the user guide (10X protocol #CG000239) and sequenced on the Illumina Novoseq 6000 system S1-200 flowcell.

### Resolve Highly Multiplexed In situ RNA Profiling using smFISH

Resolve Biosciences probes were designed to target 100 genes based on the top expressed genes in each of the breast cell types from the scRNA-seq data and are listed in Supplementary Table 6. To prepare the tissue for Resolve smFISH analysis, OCT embedded tissues were cut to 12μm sections in a microtome with chuck and blade temperatures set at −17 °C and −15 °C, respectively. Tissue sections were thawed and fixed with 4% v/v Formaldehyde (Sigma-Aldrich F8775) in 1x PBS for 30 min at 4 °C. After fixation, sections were washed for one minute in washed 50% Ethanol and 70% Ethanol at room temperature. Fixed samples were used for Molecular Cartography according to the manufacturer’s instructions (protocol 3.0; www.resolvebiosciences.com), starting with the aspiration of ethanol and the addition of buffer BST1 (step 6 and 7 of the tissue priming protocol). Briefly, tissues were primed followed by overnight hybridization of all probes specific for the target genes (see below for probe design details and target list). Samples were washed the next day to remove excess probes and fluorescently tagged in a two-step color development process. Regions of interest were imaged as described below and fluorescent signals removed during decolorization. Color development, imaging and decolorization were repeated for cycles to build a unique combinatorial code for every target gene that was derived from raw images as described below. Samples were imaged on a Zeiss Celldiscoverer 7, using the 50x Plan Apochromat water immersion objective with an NA of 1.2 and the 0.5x magnification changer, resulting in a 25x final magnification. Standard CD7 LED excitation light source, filters, and dichroic mirrors were used together with customized emission filters optimized for detecting specific signals. Excitation time per image was 1000 ms for each channel (DAPI was 20 ms). A z-stack was taken at each region with a distance per z-slice according to the Nyquist-Shannon sampling theorem. The custom CD7 CMOS camera (Zeiss Axiocam Mono 712, 3.45 µm pixel size) was used.

For each region, a z-stack per fluorescent color (two colors) was imaged per imaging round. A total of 8 imaging rounds were done for each position, resulting in 16 z-stacks per region. The completely automated imaging process per round (including water immersion generation and precise relocation of regions to image in all three dimensions) was realized by a custom python script using the scripting API of the Zeiss ZEN software (Open application development).

### Highly-multiplexed immunostaining using co-detection by indexing (CODEX)

Formalin fixed paraffin embedded human breast tissue was analyzed using COdetection by inDEXing (CODEX®/PhenoCycler - Akoya Biosciences, Marlborough, MA). The experiments were performed following manufacturer’s protocols. Briefly, the tissue was sectioned at 5-7 µm and mounted onto 22×22 mm glass coverslips, previously coated with 0.1% poly-L-lysine. The tissue section was dewaxed and stained with a mixture of oligonucleotide-barcoded PhenoCycler antibodies and post-fixed according to the PhenoCycler user manual. The tissue was then imaged on the PhenoCycler-Open platform, whereby three fluorescent oligo reporters with spectrally distinct dyes are applied to the tissue in iterative imaging cycles. Imaging data were acquired with a Keyence BZ-X800 fluorescent microscope at 20x magnification. The tissue was stained with a 34-antibody panel targeting proteins listed in Supplementary Table 10.

### RNAscope *in situ* hybridization combined with immunofluorescence

To simultaneously detect MMP3 mRNA and Vimentin and Pan-Cytokeratin (PanCK) protein *in situ* in human breast FFPE tissue sections, RNAscope® Multiplex Fluorescent Reagent Kit V2 (ACD Biotechne, Cat. 323100) was combined with immunofluorescence (IF). Manufacturer’s instructions were followed for RNAscope *in situ* hybridization unless otherwise inducated using a probe targeting human MMP3 gene (Hs-MMP3 RNAscope® Probe, Cat. 403421). 5μm FFPE tissue sections were baked at 60⁰C for 1h20min, followed by deparaffinization using Histoclear (10min, 2x) and 100% ethanol (2min, 2x). After pretreatment, hydrogen peroxide incubation, and target retrieval for 15min, a barrier was created with a hydrophobic pen and dried at room temperature (RT) for 40min. Following MMP3 probe hybridization for 2h at 40⁰C and washing, three signal amplification steps were performed and HRP signal was developed using Opal™ 570 at 1:1500 dilution (Akoya Biosciences, Cat. FP1488001KT). IF was performed after the HRP blocker step with all steps conducted in the dark. Tissue was washed twice in TBST and blocked in 10% FBS in TBS + 0.1% BSA at 4⁰C overnight. Anti-Vimentin antibody (R&D, raised in goat, Cat. AF2105) and anti-PanCK antibody (GeneTex, raised in mouse, Cat. GTX26401) were used at 1:200 and 1:500 dilution, respectively, in TBS + 0.1% BSA for 2h at RT. Following three TBS washes, donkey anti-goat-AF488 (for Vim) and donkey anti-mouse AF647 (for PanCK) were used as secondary antibodies at 1:500 dilution in TBS for 2h at RT. Tissues were washed 3x in TBS and mounted with VECTASHIELD® Antifade Mounting Medium with DAPI (Vector laboratories, Cat. H-1200). Images were acquired on a Keyence BZ- X700 using DAPI, Cy3, Cy5 and GFP filter sets.

## COMPUTATIONAL METHODS

### Code Availability

The scripts associated with the analysis are available at https://github.com/navinlabcode/HumanBreastCellAtlas.

### Single cell RNA and nuclei RNA data preprocessing and filtering

Sequencing reads from single cells and single nuclei from the 10X Genomics Chromium were demultiplexed, aligned to the GRCh38.p12 human genome reference ^3, 4^ using the default parameters of the Cell Ranger pipeline (version v3.1.0, 10x Genomics). Count matrices were generated for both datasets that were further analyzed using Seurat^5^(v. 3.2.3)^5^. Cells from each sample were further filtered for low quality by removing cells with fewer than 500 UMIs or 200 genes detected. Potential doublets and multiplets were classified as cells that express more than 20,000 UMIs or 5000 genes and were removed. Cells with higher than 10% mitochondrial or 50% ribosomal transcripts were also filtered since they represented low quality or dying cells. Similarly, for single cell nuclei, the same filtering metrics were used for the single cell data, except the min number of genes used for filtering cells was 150, since nuclei data express fewer genes.

### Clustering of Major Cell Types in scRNA-seq and snRNA-seq Data

The major cell types in the scRNA-seq data and nuclei in the snRNA-seq data clustering was done by integrating all samples together using Canonical Correlation Analysis (CCA) based integration from the Seurat package. The filtered gene matrices from each sample were normalized using *NormalizeData* function. To identify highly variable genes, we used the *FindVariableFeatures* that models the mean-variance relationship of the normalized counts of each gene across cells and identified 5000 genes per sample. We further identified anchors using *FindIntegrationAnchors* to integrate all patients using following parameters – dims=20, k.filter=30, anchor.features = 3000 and k.score = 30 which were used for the *IntegrateData* function with dims=20. The integrated dataset was then used for downstream analysis which included scaling and centering the data using *ScaleData*, finding the most significant principal components (PC) using *RunPCA* and utilizing the *ElbowPlot* to determine the number of PCs used for clustering. Different resolution parameters for unsupervised clustering were then examined to determine the optimal number of clusters. For the major cell type and nuclei clustering, the first 20 PCs were used for unsupervised clustering with a resolution = 0.2 yielding a total of 21 cell clusters and for nuclei the resolution = 0.3 yielding 21 nuclei clusters using the *FindNeighbours* and *FindClusters* functions. For visualization, the dimensionality was further reduced using eitherUMAP methods with Seurat function *RunUMAP*. The PC’s used to calculate the UMAP embedding were the same as those used for clustering. Each resulting cluster was further analyzed for potential doublets or low quality cells using a 3 step process – 1) Calculation the quality metrics such as nCount_RNA and mitochondrial content and removed clusters with any outlier values (greater or less than 2 sd than the average of all clusters), 2) Checked the top 15 differentially expressed genes of each cluster and removed the clusters where genes were predominantly mitochondrial, ribosomal or hemoglobin genes, 3) using the canonical cell type markers for each cell types, we determined if any cluster had cells expressing canonical markers from a different cell type, suggesting they are doublets with another cell types. Based on the above criteria, we identified 10 major cell type clusters and 12 nuclei clusters that were well separated in UMAP space.

### Assignment of Cell Type Annotations to Clusters

To annotate the major cell type of each single cell or nuclei, *FindAllMarkers* was used to find differentially expressed genes in each cluster using the Wilcoxon rank sum test statistical framework. The top 12 most significant DEGs (ranked by average log fold change; adjusted *P* values < 0.05) were then carefully reviewed. Further, we further checked each cluster using the known canonical markers such as EPCAM for epithelial cells, PTPRC for immune cells, CD3D/E/G for T cells, CD19/MS4A1/CD79A for B cells, LUM/DCN/COL6A1 for fibroblasts, PECAM1 for endothelial cells and RGS5 for pericytes. We also applied SingleR6 to annotate the clusters. The three approaches were combined to infer major cell types for each cell and nuclei cluster according to the resulting annotation designated by SingleR and the enrichment of canonical marker genes and top-ranked DEGs in each cell cluster.

### Identification of Cell States by Reclustering of Cell Type Data

Each cell cluster was further extracted and underwent clustering and filtering as described above, however with different parameters. The different parameters used for clustering the expression states of major cells are as follows - B cells (dims = 12; k.param = 20, scaled by nCount_RNA, resolution = 0.3), T cells (dims = 20; k.param = 20, scaled by nCount_RNA, resolution = 0.4), Myeloid cells (dims = 30; k.param = 20, scaled by nCount_RNA, resolution = 0.4), Fibroblasts (dims = 30, k.param = 20, scaled by nCount_RNA, resolution = 0.4), LumHR (dims = 35; k.param = 20, resolution = 0.075, scaled by nCount_RNA), LumSec (dims = 35; k.param = 20, resolution = 0.2 scaled by nCount_RNA), Pericytes (dims = 25; k.param = 20, scaled by nCount_RNA, resolution = 0.4), Lymphatic (dims = 30; resolution = 0.05) and Vascular Endothelial cells (dims = 30; resolution = 0.1. For vascular cells, one cluster was only from overnight digestion. To avoid potential dissociation artifact, we removed this cluster OD from further downstream analysis). Each round of clustering was followed by filtering for low quality and doublets cells. Differentially expressed genes were calculated for each cell cluster relative to other cells within its cell type compartment using the “FindMarkers” function in Seurat with the Wilcoxon rank sum test for statistical significance. Expression states were further annotated by investigating the top 200 genes of each cluster and performing pathway enrichment on the cell states as described in the Pathway Enrichment section. For each cell type, we showed top genes of each cell state in the heatmaps based on the average log fold change.

### Cell cycle analysis

We utilized the “CellCycleScoring” function from the Seurat package that is based on the cell cycle phase genes from the paper by Tirosh et al ^7^. Each cell and nuclei were given a quantitative score for G1, G2/M and S scores based on scoring of marker genes at each stage of the cell cycle.

### Pathway enrichment analysis

For Gene set enrichment analysis, ranked genes were selected based on the above test filtered for an adjusted p-value <=0.05 and arranged by average log foldchange values between each cluster and fed into the ‘fgsea’ R-package^8^ using 1000 permutations. Curated gene sets of KEGG, Biological Processes and Reactome were downloaded from the Molecular Signature Database (MSigDB, http://software.broadinstitute.org/gsea/msigdb/index.jsp) and were used to calculate enrichment scores. Significantly enriched gene sets were identified with a Benjamini-Hochberg adjusted P value <= 0.05. For the identified cell states, we also selected top 200 genes and performed GO and KEGG enrichment analysis using the clusterProfiler package^9^.

### Regulatory network analysis

On the raw count matrix, we applied SCENIC^10^ to inference the regulatory networks following the instructions from https://scenic.aertslab.org/. On the regulon score matrix, we performed differential expression (DE) analysis following the similar approach in DE gene analysis and identified top regulons for each cell type.

### Spatial analysis of smFISH Resolve data

We applied QuPath (version 0.3.0)^11^ to segment cells based on their DAPI images, then applied ImageJ (version 1.52n)^12^ and the Molecular Cartography plug-in (Resolve Biosciences) to count genes in each cell. For the DAPI image, we also manually annotated different regions (duct, lobule, connective tissue and fibrocysts) by matched pathology H&E sections using ImageJ. The cell-gene count matrix was then input into Seurat (v. 3.2.3)^5^ for downstream analysis. For the 12 samples of spatial resolve data, cells with less than 10 gene counts were filtered. Counts data were then normalized using the *NormalizeData* with default *LogNormalize* method. Afterwards, normalized counts were scaled and centered using *ScaleData* function. All genes were used for principal component analysis (PCA) using *RunPCA* with default parameters. *ElbowPlot* was used to determine the number of PCs for the downstream analyses and *RunUMAP* was applied to reduce data to a 2D space. We applied a two-steps approach to annotate cells. First, we curated a marker list of each cell type and used *AddModuleScore* from *Seurat* to calculate the cell type scores of each cell (Supplementary Table 6). By comparing cell type scores, we took the largest score to assign the cell types and assigned cells with all scores less than 0.5 as low confident cells. Then, a random forest machine learning model with a default of 500 trees was trained on the data while setting the cell type assignment as output and top 20 PCs as predictors using *randomForest* package^13^ (CRAN). Out-of-bag predictions were used as our final cell type annotation while cells with largest voting rate less than 0.5 were assigned to the low confident group and were filtered for the downstream analysis. Cell type differentially expressed (DE) genes were identified using *FindAllMarkers*. Cell spatial colocalization graph was calculated using *scoloc* function with *DT* method from *CellTrek* package^14^.

### Spatial transcriptomics data analysis

Sequencing reads from Visium ST (10X Genomics) experiments were first preprocessed with *Space Ranger* v1.2.0 (10X Genomics) and mapped to the *GRCh38* reference genome. The count matrices were subsequently analyzed using Seurat v3. We filtered out spots with a total count less than 100. The UMI counts were normalized using *SCTransform*. Similar to the scRNA-seq analysis, we then used Seurat anchor-based integration with the default parameters for the four samples. After integration, dimensionality reduction was performed using the *RunPCA* function. Clustering was done using the *FindClusters* function. Differentially expressed genes for each ST cluster were identified using *FindAllMarkers* with two-sided Wilcoxon rank sum test. P values were adjusted by Bonferroni correction and genes with adjusted *P* < 0.05 were retained.

### Procrustes analysis between the left and right breast

In contralateral samples, we calculated Bray-Curtis dissimilarity between samples using the *vegdist* function from the vegan package (2.5-6) using R studio [https://cran.rproject.org/web/packages/vegan/index.html] based on the cell type count matrix and then performed a multidimensional scaling using *cmdscale.* To compare the cell type composition between the left and right breast, we applied Procrustes analysis using the *protest* function from the vegan package and performed a 9,999 times of permutation test. Pearson’s correlation test was used to measure the similarities on top-2 dimensions between the left and right breast on after Procrustes rotations.

### Statistics and reproducibility

The Wilcoxon signed-rank test was used to evaluate associations between clinical metavariables for comparing celltype frequencies. Fisher exact test was used to compare counts of patients having a minimum of 20 cells of each cellstate. The p values from the 2-tailed tests are reported for each comparison and only the significant ones are shown in the Extended Data Fig 11.

### Computational analysis of CODEX data

StarDist trained on TissueNet dataset (https://datasets.deepcell.org/data) was used for cell segmentation. Then, the average intensity of each protein was calculated for individual cells using the segmentation masks and the protein images. If the protein was localized in nuclei, e.g., Ki67, PCNA, FoxP3, then the average intensity was calculated from the nuclear mask obtained with StarDist. Otherwise, if the protein was localized in the cell membrane, e.g., CD4, CD3, E-Cadherin, then the average intensity was calculated from the membrane mask. The average protein intensities were then z-scored across all cells. Unsupervised clustering using Leiden algorithm was performed based on the normalized average protein intensities to assign cluster labels for all cells in each sample individually. The average intensity of each protein was then re calculated for each cluster and displayed on a heatmap to identify cell types manually based on marker expression, e.g., basal, luminal, fibroblast, T cells, myeloid, endothelial cells. Tissue regions were manually annotated into lobules, ducts, and connective tissue based on their histological structure and morphology. The number of cell types in each annotated region was counted, and the relative cell percentages and densities (cell count per area unit) were compared between the different regions. For the immune cell-specific analysis, we took out the previously identified myeloid and T cells clusters and increased the clustering resolution and identified CD8 T cells, CD4 T cells, T regulatory cells, monocytes, macrophages, and dendritic cells. RunX3 positive cells were defined by examining the gene expression distribution.

