## supplementary figures and tables for "A spatially resolved single cell genomic atlas of the adult human breast"

### Supplementary Materials

Kumar, Nee, Wei, He, Nguyen et al.

Extended Data Fig. 1 – Variation of Major Breast Cell Types Across Samples and Women ……………………………3

Extended Data Fig. 2 – Spatial Transcriptomic Analysis of Breast Cell Types …………………………………………….5

Extended Data Fig. 3 – Spatial In Situ Analysis of Breast Cell Types with CODEX and smFISH ……………………….7

Extended Data Fig. 4 – Analysis of Single Cell and Nuclei Epithelial Data …………………………………………………9

Extended Data Fig. 5 – Spatial analysis of epithelial cells in ductal and lobular structures ……………………………..11

Extended Data Fig. 6 – Canonical marker expression and frequency of immune cell subsets of the human breast ...13

Extended Data Fig. 7 – Spatial analysis of immune cells of the human breast …………………………………………..15

Supplementary Table 4 - Top Marker Genes Expressed in the Breast Cell Types from Nuclei data …………………..33

Supplementary Table 5 - Quality Control Metrics for Spatial Transcriptomics Data …………………………………….35

Supplementary Table 6 - Custom Targeted Gene Panel for smFISH ……………………………………………………..36

Supplementary Table 7 - Quality Control Metrics for smFISH …………………………………………………………...37

Supplementary Table 8 - Samples for CODEX ………………………………………………………………………………38

Supplementary Table 9 - CODEX 34-Antibody Panel Design …………………………………………………………...39

Supplementary Table 10 - Top Gene Markers expressed for Breast Cell States ………………………………………...41

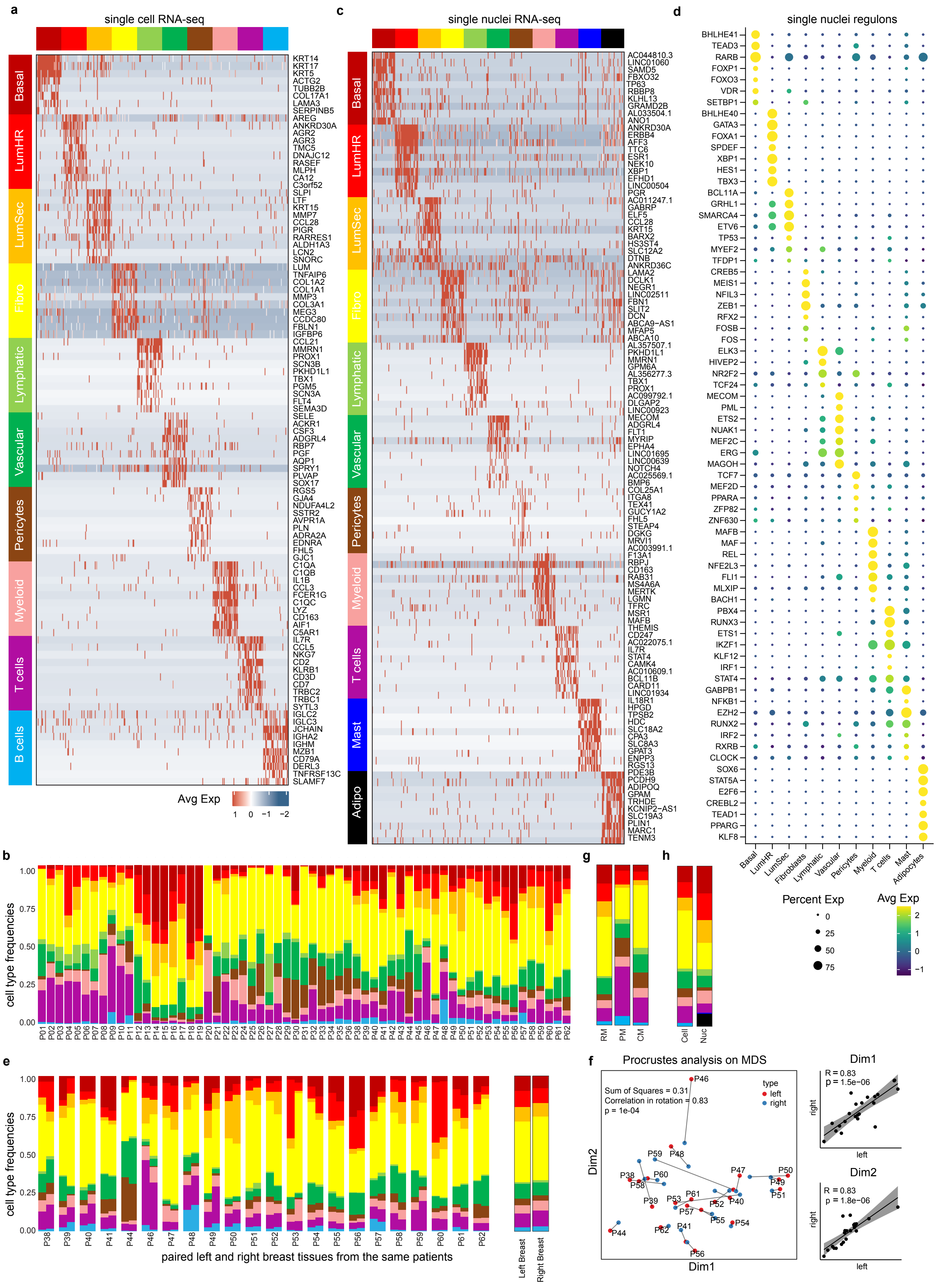
a
single cell RNA-seq
Basal
LumHR
LumSec
Fibro
Lymphatic
Vascular
Pericytes
Myeloid
T cells
B cells
KRT14 KRT17 KRT5 ACTG2 TUBB2B COL17A1 LAMA3 SERPINB5 AREG ANKRD30A AGR2 AGR3 TMC5 DNAJC12 RASEF MLPH CA12 C3orf52 SLPI LTF KRT15 MMP7 CCL28 PIGR RARRES1 ALDH1A3 LCN2 SNORC LUM TNFAIP6 COL1A2 COL1A1 MMP3 COL3A1 MEG3 CCDC80 FBLN1 IGFBP6 CCL21 MMRN1 PROX1 SCN3B PKHD1L1 TBX1 PGM5 SCN3A FLT4 SEMA3D SELE ACKR1 CSF3 ADGRL4 RBP7 PGF AQP1 SPRY1 PLVAP SOX17 RGS5 GJA4 NDUFA4L2 SSTR2 AVPR1A PLN ADRA2A EDNRA FHL5 GJC1 C1QA C1QB IL1B CCL3 FCER1G C1QC LYZ CD163 AIF1 C5AR1 IL7R CCL5 NKG7 CD2 KLRB1 CD3D CD7 TRBC2 TRBC1 SYTL3 IGLC2 IGLC3 JCHAIN IGHA2 IGHM MZB1 CD79A DERL3 TNFRSF13C SLAMF7
Avg Exp
1 0 −1 −2
c
single nuclei RNA-seq
Basal
LumHR
LumSec
Fibro
Lymphatic
Vascular
Pericytes
Myeloid
T cells
Mast
Adipo
AC044810.3 LINC01060 SAMD5 FBXO32 TP63 RBBP8 KLHL13 GRAMD2B AL033504.1 ANO1 ANKRD30A ERBB4 AFF3 TTC6 ESR1 NEK10 XBP1 EFHD1 LINC00504 PGR AC011247.1 GABRP ELF5 CCL28 KRT15 BARX2 HS3ST4 SLC12A2 DTNB ANKRD36C LAMA2 DCLK1 NEGR1 LINC02511 FBN1 SLIT2 DCN ABCA9-AS1 MFAP5 ABCA10 AL357507.1 PKHD1L1 MMRN1 GPM6A AL356277.3 TBX1 PROX1 AC099792.1 DLGAP2 LINC00923 MECOM ADGRL4 FLT1 MYRIP EPHA4 LINC01695 LINC00639 NOTCH4 AC025569.1 BMP6 COL25A1 ITGA8 TEX41 GUCY1A2 FHL5 STEAP4 DGKG MRVI1 AC003991.1 F13A1 RBPJ CD163 RAB31 MS4A6A MERTK LGMN TFRC MSR1 MAFB THEMIS CD247 AC022075.1 IL7R STAT4 CAMK4 AC010609.1 BCL11B CARD11 LINC01934 IL18R1 HPGD TPSB2 HDC SLC18A2 CPA3 SLC8A3 GPAT3 ENPP3 RGS13 PDE3B PCDH9 ADIPOQ GPAM TRHDE KCNIP2-AS1 SLC19A3 PLIN1 MARC1 TENM3
d
single nuclei regulons
BHLHE41 TEAD3 RARB FOXP1 FOXO3 VDR SETBP1 BHLHE40 GATA3 FOXA1 SPDEF XBP1 HES1 TBX3 BCL11A GRHL1 SMARCA4 ETV6 TP53 MYEF2 TFDP1 CREB5 MEIS1 NFIL3 ZEB1 RFX2 FOSB FOS ELK3 HIVEP2 NR2F2 TCF24 MECOM PML ETS2 NUAK1 MEF2C ERG MAGOH TCF7 MEF2D PPARA ZFP82 ZNF630 MAFB MAF REL NFE2L3 FLI1 MLXIP BACH1 PBX4 RUNX3 ETS1 IKZF1 KLF12 IRF1 STAT4 GABPB1 NFKB1 EZH2 RUNX2 IRF2 RXRB CLOCK SOX6 STAT5A E2F6 CREBL2 TEAD1 PPARG KLF8
Basal LumHR LumSec Fibroblasts Lymphatic Vascular Pericytes Myeloid T cells Mast Adipocytes
Percent Exp
0
25
50
75
Avg Exp
2
1
0
−1
b
cell type frequencies
1.00 0.75 0.50 0.25 0.00
P01 P02 P03 P04 P05 P06 P07 P08 P09 P10 P11 P12 P13 P14 P15 P16 P17 P18 P19 P20 P21 P22 P23 P24 P25 P26 P27 P28 P29 P30 P31 P32 P33 P34 P35 P36 P37 P38 P39 P40 P41 P42 P43 P44 P45 P46 P47 P48 P49 P50 P51 P52 P53 P54 P55 P56 P57 P58 P59 P60 P61 P62
g
RM PM CM
h
Cell Nuc
e
cell type frequencies
1.00 0.75 0.50 0.25 0.00
P38 P39 P40 P41 P44 P46 P47 P48 P49 P50 P51 P52 P53 P54 P55 P56 P57 P58 P59 P60 P61 P62
paired left and right breast tissues from the same patients
Left Breast
Right Breast
f
Procrustes analysis on MDS
Sum of Squares = 0.31
Correlation in rotation = 0.83
p = 1e-04
type
left
right
P46 P59 P48 P38 P60 P58 P47 P50 P49 P51 P39 P53 P61 P52 P40 P57 P55 P54 P44 P62 P41 P56
Dim2
Dim1
Dim1
R = 0.83
p = 1.5e−06
right
left
Dim2
R = 0.83
p = 1.8e−06
right
left

**Extended Data Fig. 1 – Variation of Major Breast Cell Types Across Samples and Women**
**a,** Heatmap of scRNA-seq data clustered by cell type, showing the top marker genes for 100 randomly sampled cells per cluster. **b,** Stacked barplots showing the variation of cell type frequencies across the 62 women. **c,** Heatmap of snRNA-seq data clustered by cell type, showing the top marker genes for 100 randomly sampled cells per cluster. **d,** Top transcription factors and regulons identified with SCENIC for each cell type cluster from the snRNA-seq data **e,** Major cell type frequencies of matched left and right breasts from 22 women (left panel) and averages across all left and all right breasts (right panel). **f,** Multi-dimensional scaling Procrustes analysis to determine the concordance of left and right breast cell type frequencies and Pearson correlations for the 22 women with matched breast tissue samples **g,** Cell type frequencies for different tissue sources, including reduction mammoplasties (RM), prophylactic mastectomies (PM) and contralateral mastectomies (CM) **h,** Cell type frequencies for scRNA-seq cell data and snRNA-seq nuclei data showing differences in compositions.

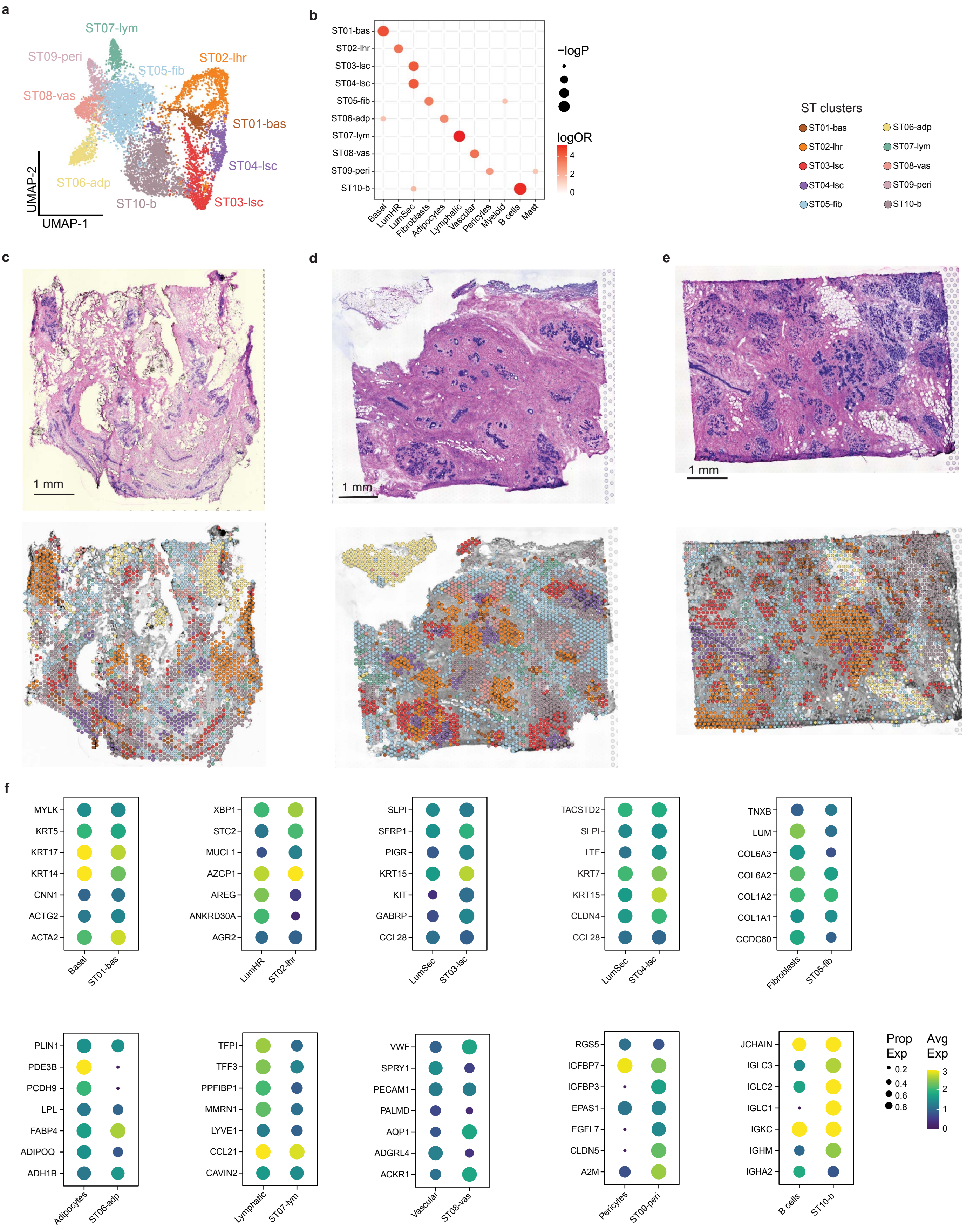
a
ST07-lym
ST09-peri
ST05-fib
ST02-lhr
ST08-vas
ST01-bas
ST04-lsc
ST06-adp
ST10-b
ST03-lsc
UMAP-2
UMAP-1
b
ST01-bas
ST02-lhr
ST03-lsc
ST04-lsc
ST05-fib
ST06-adp
ST07-lym
ST08-vas
ST09-peri
ST10-b
Basal
LumHR
LumSec
Fibroblasts
Adipocytes
Lymphatic
Vascular
Pericytes
Myeloid
B cells
Mast
−logP
logOR
4
2
0
ST clusters
ST01-bas
ST02-lhr
ST03-lsc
ST04-lsc
ST05-fib
ST06-adp
ST07-lym
ST08-vas
ST09-peri
ST10-b
c
d
e
1 mm
1 mm
1 mm
f
MYLK
KRT5
KRT17
KRT14
CNN1
ACTG2
ACTA2
Basal
ST01-bas
XBP1
STC2
MUCL1
AZGP1
AREG
ANKRD30A
AGR2
LumHR
ST02-lhr
SLPI
SFRP1
PIGR
KRT15
KIT
GABRP
CCL28
LumSec
ST03-lsc
TACSTD2
SLPI
LTF
KRT7
KRT15
CLDN4
CCL28
LumSec
ST04-lsc
TNXB
LUM
COL6A3
COL6A2
COL1A2
COL1A1
CCDC80
Fibroblasts
ST05-fib
PLIN1
PDE3B
PCDH9
LPL
FABP4
ADIPOQ
ADH1B
Adipocytes
ST06-adp
TFPI
TFF3
PPFIBP1
MMRN1
LYVE1
CCL21
CAVIN2
Lymphatic
ST07-lym
VWF
SPRY1
PECAM1
PALMD
AQP1
ADGRL4
ACKR1
Vascular
ST08-vas
RGS5
IGFBP7
IGFBP3
EPAS1
EGFL7
CLDN5
A2M
Pericytes
ST09-peri
JCHAIN
IGLC3
IGLC2
IGLC1
IGKC
IGHM
IGHA2
B cells
ST10-b
Prop Exp
0.2
0.4
0.6
0.8
Avg Exp
3
2
1
0

**Extended Data Fig. 2 – Spatial Transcriptomic Analysis of Breast Cell Types**
**a,** Integrated UMAP and unbiased clustering of ST data from 4 breast samples, showing 10 major cell type clusters. **b,** Concordance of ST cluster marker genes and the scRNA-seq data clusters of the major cell types. **c-e,** Histopathological images, and spatial distribution of the 10 ST clusters in the ST data from the breast tissues of three women (P10, P47 and P46). **f,** *in situ* validation and concordance of top marker gene expression levels between the ST clusters and the scRNA-seq or snRNA-seq data for different cell types.

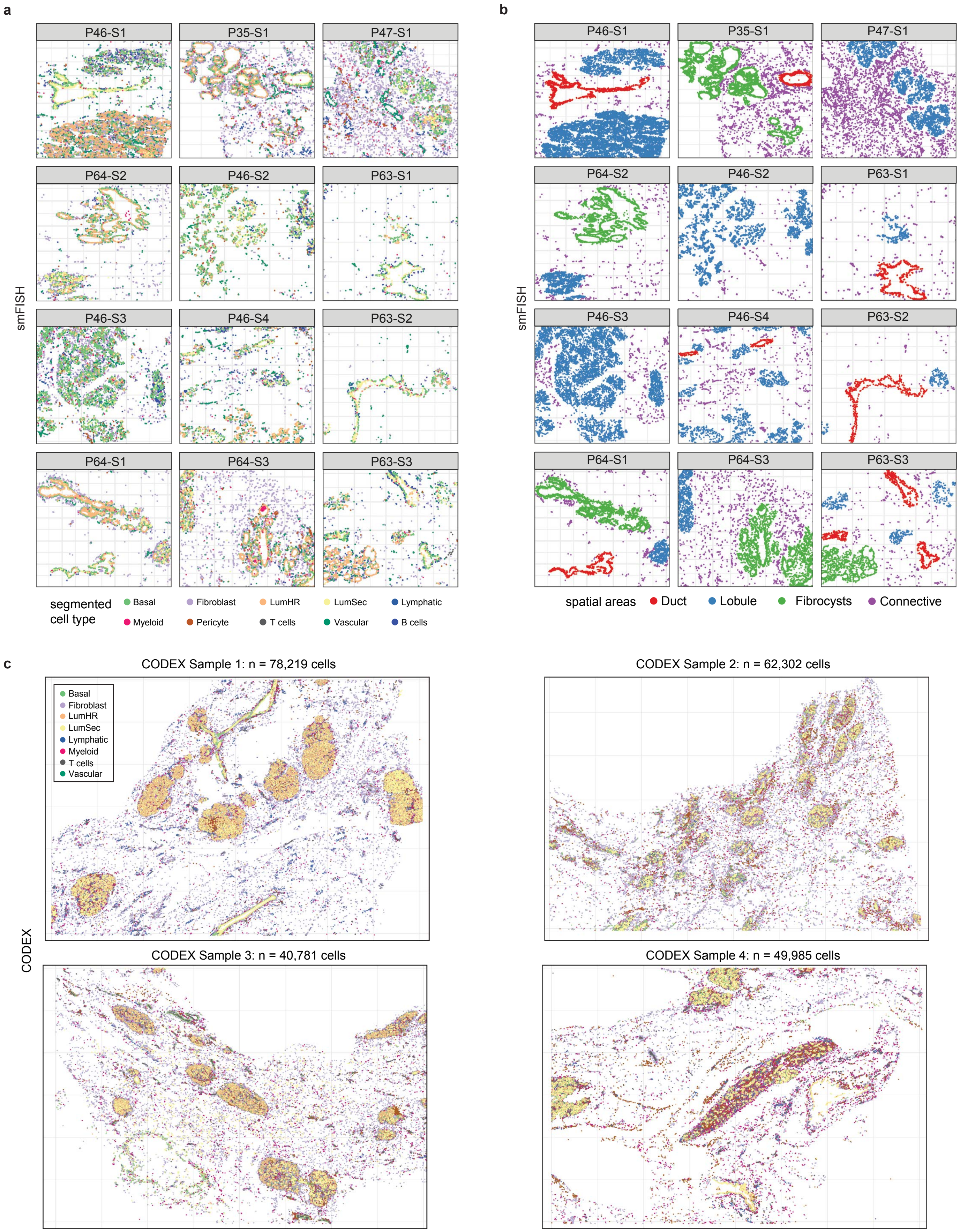
a
P46-S1
P35-S1
P47-S1
P64-S2
P46-S2
P63-S1
P46-S3
P46-S4
P63-S2
P64-S1
P64-S3
P63-S3
smFISH
segmented cell type
Basal
Fibroblast
LumHR
LumSec
Lymphatic
Myeloid
Pericyte
T cells
Vascular
B cells
b
spatial areas
Duct
Lobule
Fibrocysts
Connective
c
CODEX Sample 1: n = 78,219 cells
CODEX Sample 2: n = 62,302 cells
CODEX Sample 3: n = 40,781 cells
CODEX Sample 4: n = 49,985 cells
CODEX

**Extended Data Fig. 3 – Spatial *In Situ* Analysis of Breast Cell Types with CODEX and smFISH**

**a,** Cell segmented smFISH spatial *in situ* data from 12 tissue samples profiled from 4 different women. Cells were segmented based on combinations of markers for each cell type. **b,** Pathologist annotated anatomical regions in the breast tissue labeled as ducts, lobules, fibrocysts or connective areas. **c,** Cell segmentation results of CODEX data from 4 different women using combinations or single protein markers to identify different cell types.

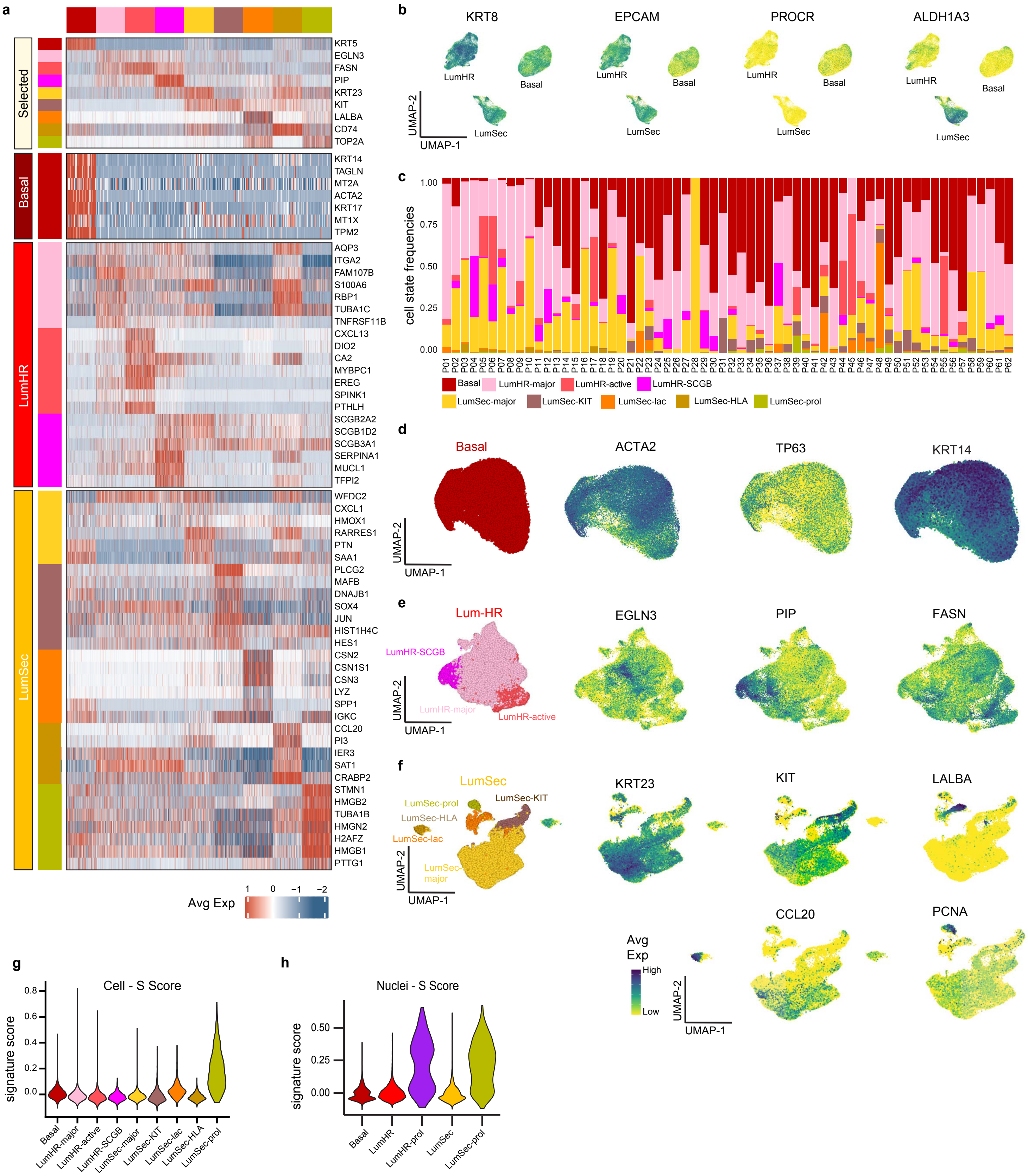
a
Selected
Basal
LumHR
LumSec
KRT5
EGLN3
FASN
PIP
KRT23
KIT
LALBA
CD74
TOP2A
KRT14
TAGLN
MT2A
ACTA2
KRT17
MT1X
TPM2
AQP3
ITGA2
FAM107B
S100A6
RBP1
TUBA1C
TNFRSF11B
CXCL13
DIO2
CA2
MYBPC1
EREG
SPINK1
PTHLH
SCGB2A2
SCGB1D2
SCGB3A1
SERPINA1
MUCL1
TFPI2
WFDC2
CXCL1
HMOX1
RARRES1
PTN
SAA1
PLCG2
MAFB
DNAJB1
SOX4
JUN
HIST1H4C
HES1
CSN2
CSN1S1
CSN3
LYZ
SPP1
IGKC
CCL20
PI3
IER3
SAT1
CRABP2
STMN1
HMGB2
TUBA1B
HMGN2
H2AFZ
HMGB1
PTTG1
Avg Exp
1 0 −1 −2
b
KRT8
EPCAM
PROCR
ALDH1A3
LumHR
Basal
LumSec
UMAP-2
UMAP-1
c
cell state frequencies
1.00
0.75
0.50
0.25
0.00
P01 P02 P03 P04 P05 P06 P07 P08 P09 P10 P11 P12 P13 P14 P15 P16 P17 P18 P19 P20 P21 P22 P23 P24 P25 P26 P27 P28 P29 P30 P31 P32 P33 P34 P35 P36 P37 P38 P39 P40 P41 P42 P43 P44 P45 P46 P47 P48 P49 P50 P51 P52 P53 P54 P55 P56 P57 P58 P59 P60 P61 P62
Basal
LumHR-major
LumHR-active
LumHR-SCGB
LumSec-major
LumSec-KIT
LumSec-lac
LumSec-HLA
LumSec-prol
d
Basal
ACTA2
TP63
KRT14
UMAP-2
UMAP-1
e
Lum-HR
LumHR-SCGB
LumHR-major
LumHR-active
EGLN3
PIP
FASN
UMAP-2
UMAP-1
f
LumSec
LumSec-prol
LumSec-KIT
LumSec-HLA
LumSec-lac
LumSec-major
KRT23
KIT
LALBA
CCL20
PCNA
Avg Exp
High
Low
UMAP-2
UMAP-1
g
Cell - S Score
signature score
0.8
0.6
0.4
0.2
0.0
Basal
LumHR-major
LumHR-active
LumHR-SCGB
LumSec-major
LumSec-KIT
LumSec-lac
LumSec-HLA
LumSec-prol
h
Nuclei - S Score
signature score
0.50
0.25
0.00
Basal
LumHR
LumHR-prol
LumSec
LumSec-prol

**Extended Data Fig. 4 – Analysis of Single Cell and Nuclei Epithelial Data**
**a,** Heatmap of epithelial scRNA-seq data clustered by cell states showing the top expressed genes for 100 randomly sampled cells per cluster. **b,** UMAP feature plots showing the expression of *KRT8*, *EPCAM*, *PROCR* and *ALDH1A3* genes in epithelial cell types. **c,** Epithelial cell state frequencies and their variation across the 62 women. **d,** UMAP feature plots showing the expression of canonical marker genes in basal epithelial cells. **e,** UMAP feature plots showing the expression of top cell state marker genes for the LumHR cells. **f,** UMAP feature plots showing the expression of top cell state genes of each LumSec epithelial cells. **g,** Cell cycle scoring of S-phase for different epithelial cell states detected in the scRNA-seq data, **j,** Cell cycle scoring for S-phase in the epithelial cell type clusters detected in the snRNA-seq data.

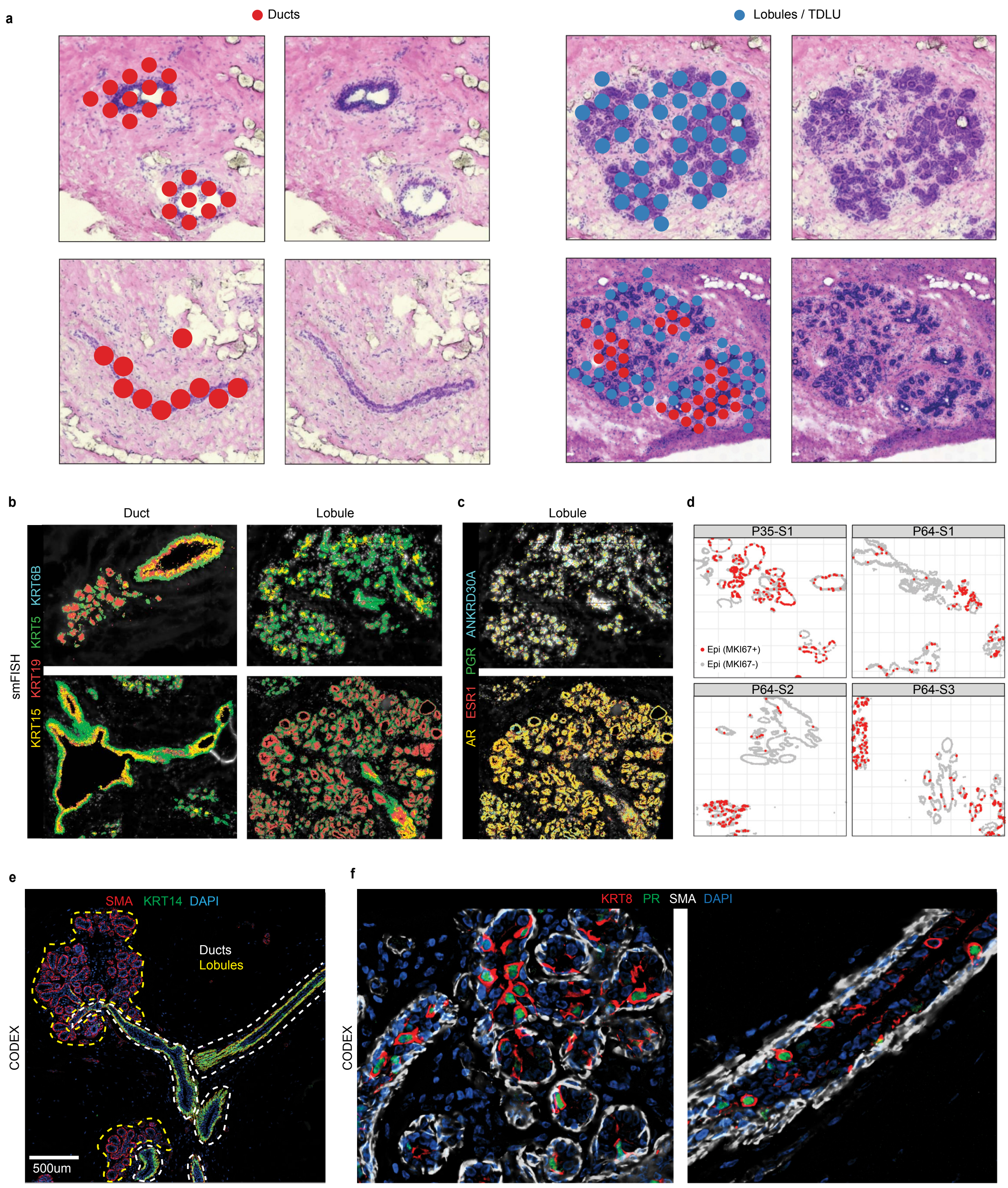
a
Ducts
Lobules / TDLU
b
Duct
Lobule
smFISH
KRT15 KRT19 KRT5 KRT6B
c
Lobule
AR ESR1 PGR ANKRD30A
d
P35-S1
P64-S1
P64-S2
P64-S3
Epi (MKI67+)
Epi (MKI67-)
e
SMA KRT14 DAPI
Ducts
Lobules
CODEX
500um
f
KRT8 PR SMA DAPI
CODEX

**Extended Data Fig. 5 – Spatial analysis of epithelial cells in ductal and lobular structures**
**a,** Spatial transcriptomic analysis showing clusters labelled as ducts or lobules/TDLUs based on pathological annotation from the histological staining of two breast tissues (P35 and P46). **b,** smFISH data (P46-S1 and P46-S4) showing a subset of Keratin markers and their localization to 4 different breast tissue regions annotated as either ducts or lobules/TDLUs. **c,** smFISH data (P46-S1) showing the expression hormone receptors in a lobular/TDLU region from one breast tissue. **d,** smFISH data showing the expression of the MKI67 proliferation marker in the epithelial cells of the ducts and lobules from 4 different breast tissues. **e,** CODEX analysis from P66 of ductal and lobular/TDLU regions, showing differences for KRT14 levels in ducts and lobules **f,** CODEX data from P67 showing protein levels of KRT8 and progesterone receptor (PR) in epithelial cells in the ducts and lobular/TDLU regions.

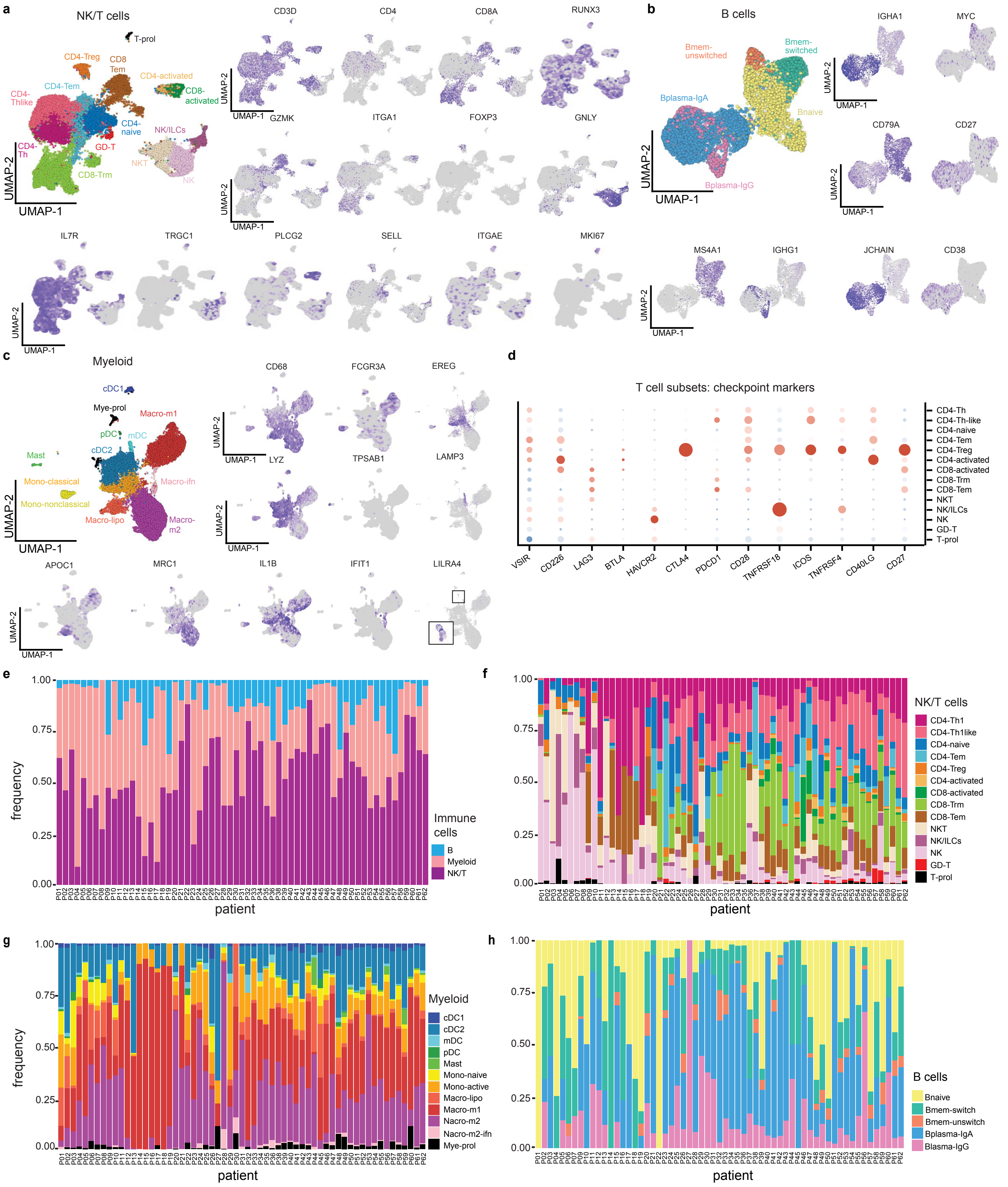

a
NK/T cells
T-prol
CD4-Treg
CD8 Tem
CD4-activated
CD8-activated
CD4-Tem
CD4-Thlike
CD4-naive
NK/ILCs
CD4-Th
GD-T
NKT
NK
CD8-Trm
UMAP-2
UMAP-1
CD3D
CD4
CD8A
RUNX3
GZMK
ITGA1
FOXP3
GNLY
IL7R
TRGC1
PLCG2
SELL
ITGAE
MKI67
b
B cells
Bmem-unswitched
Bmem-switched
Bplasma-IgA
Bnaive
Bplasma-IgG
IGHA1
MYC
CD79A
CD27
MS4A1
IGHG1
JCHAIN
CD38
c
Myeloid
cDC1
Mye-prol
Macro-m1
pDC
mDC
cDC2
Mast
Mono-classical
Macro-ifn
Mono-nonclassical
Macro-lipo
Macro-m2
CD68
FCGR3A
EREG
LYZ
TPSAB1
LAMP3
APOC1
MRC1
IL1B
IFIT1
LILRA4
d
T cell subsets: checkpoint markers
CD4-Th
CD4-Th-like
CD4-naive
CD4-Tem
CD4-Treg
CD4-activated
CD8-activated
CD8-Trm
CD8-Tem
NKT
NK/ILCs
NK
GD-T
T-prol
VSIR
CD226
LAG3
BTLA
HAVCR2
CTLA4
PDCD1
CD28
TNFRSF18
ICOS
TNFRSF4
CD40LG
CD27
e
frequency
patient
Immune cells
B
Myeloid
NK/T
f
NK/T cells
CD4-Th1
CD4-Th1like
CD4-naive
CD4-Tem
CD4-Treg
CD4-activated
CD8-activated
CD8-Trm
CD8-Tem
NKT
NK/ILCs
NK
GD-T
T-prol
g
Myeloid
cDC1
cDC2
mDC
pDC
Mast
Mono-naive
Mono-active
Macro-lipo
Macro-m1
Nacro-m2
Nacro-m2-ifn
Mye-prol
h
B cells
Bnaive
Bmem-switch
Bmem-unswitch
Bplasma-IgA
Blasma-IgG

**Extended Data Fig. 6 – Canonical marker expression and frequency of immune cell subsets of the human breast**

**a-c,** Feature plots show canonical marker expression in NK/T, B and myeloid cell subpopulations from scRNA-seq data of 62 women. **d,** Dot plot shows expression of checkpoint markers in NK and T cell subpopulations from scRNA-seq data of 62 women. **e-h,** stacked barplots showing the variation of NK/T, B and myeloid cell subpopulation frequencies across the 62 women.

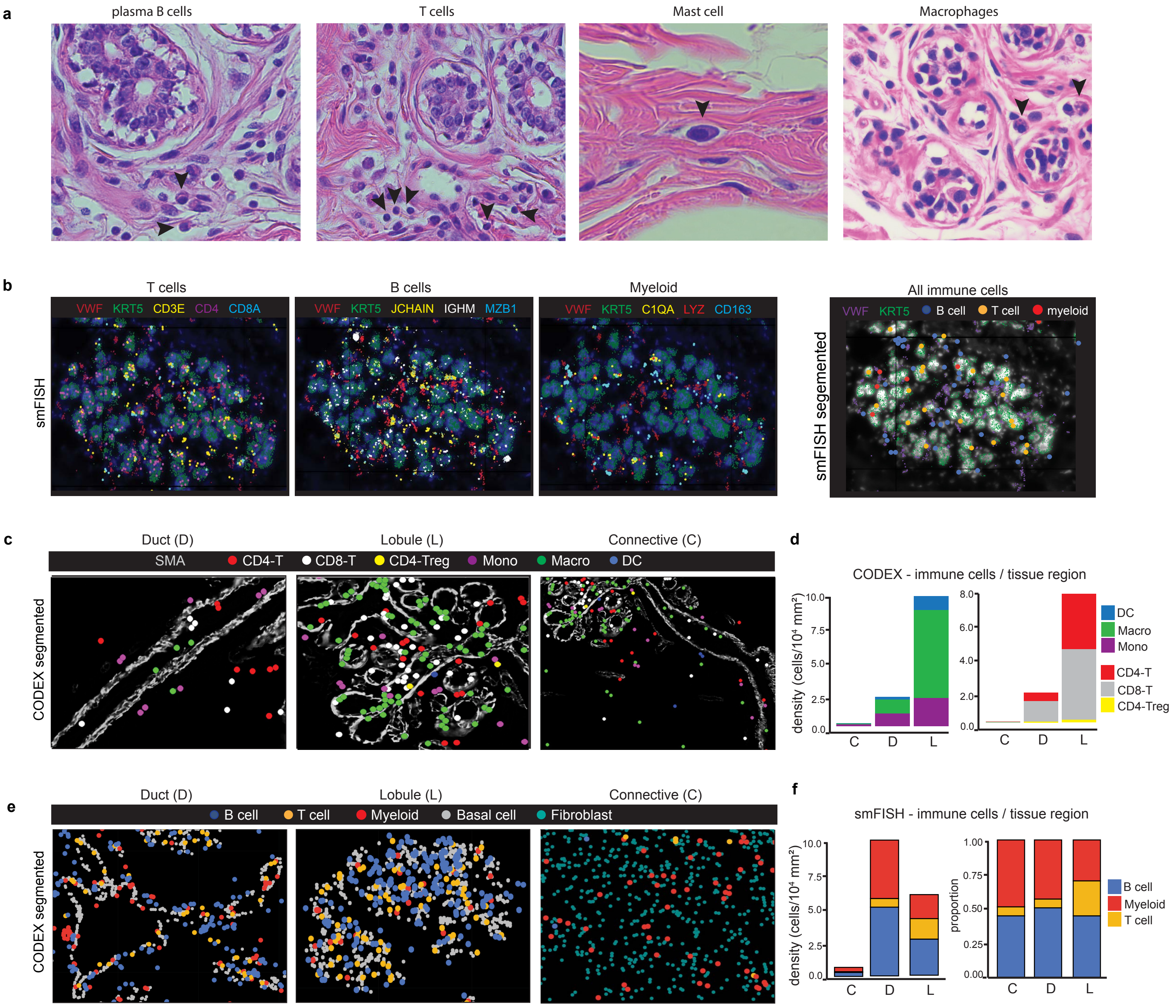

a
plasma B cells
T cells
Mast cell
Macrophages
b
T cells
B cells
Myeloid
All immune cells
VWF KRT5 CD3E CD4 CD8A
VWF KRT5 JCHAIN IGHM MZB1
VWF KRT5 C1QA LYZ CD163
VWF KRT5 B cell T cell myeloid
smFISH
smFISH segemented
c
Duct (D)
Lobule (L)
Connective (C)
SMA CD4-T CD8-T CD4-Treg Mono Macro DC
CODEX segmented
d
CODEX - immune cells / tissue region
density (cells/10⁴ mm²)
DC
Macro
Mono
CD4-T
CD8-T
CD4-Treg
C D L
e
Duct (D)
Lobule (L)
Connective (C)
B cell T cell Myeloid Basal cell Fibroblast
CODEX segmented
f
smFISH - immune cells / tissue region
density (cells/10⁴ mm²)
proportion
B cell
Myeloid
T cell
C D L

**Extended Data Fig. 7 – Spatial analysis of immune cells of the human breast**
**a,** H&E staining of immune cells (arrows) in human breast tissues. **b,** smFISH data (P46-S1) shows RNA localization (left panels) and segmentation (right panel) of B, T and myeloid cells using 9 RNA markers**. c,** CODEX data (P66) showing immune cells in C, D, and L regions. **d,** stacked bar graphs summarizing CODEX data showing the density of immune cell types in each spatial region from c, shown as cells per $10^4$ microns in n = 4 women. **e,** smFISH data (P46-S1 and P47-S1) showing immune cell types in C, D and L tissue regions. **f,** stacked bar graphs summarizing smFISH data showing density and frequency of immune cell types in each spatial region from e in n = 6 women.

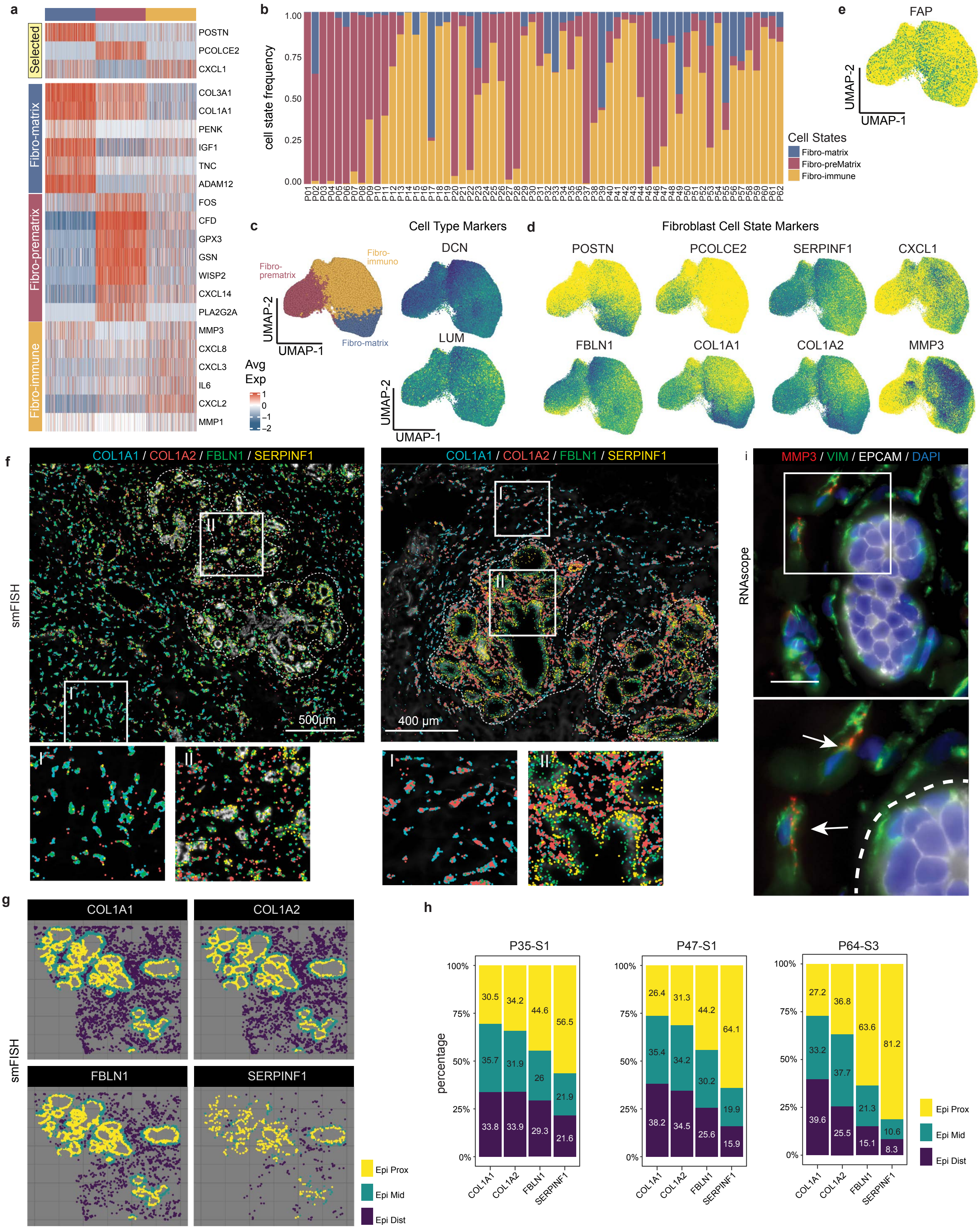
a
Selected
Fibro-matrix
Fibro-prematrix
Fibro-immune
POSTN
PCOLCE2
CXCL1
COL3A1
COL1A1
PENK
IGF1
TNC
ADAM12
FOS
CFD
GPX3
GSN
WISP2
CXCL14
PLA2G2A
MMP3
CXCL8
CXCL3
IL6
CXCL2
MMP1
Avg Exp
b
cell state frequency
Cell States
Fibro-matrix
Fibro-preMatrix
Fibro-immune
e
FAP
UMAP-2
UMAP-1
c
Cell Type Markers
Fibro-prematrix
Fibro-immuno
Fibro-matrix
DCN
LUM
d
Fibroblast Cell State Markers
POSTN
PCOLCE2
SERPINF1
CXCL1
FBLN1
COL1A1
COL1A2
MMP3
f
smFISH
COL1A1 / COL1A2 / FBLN1 / SERPINF1
500µm
400 µm
i
RNAscope
MMP3 / VIM / EPCAM / DAPI
g
COL1A1
COL1A2
FBLN1
SERPINF1
Epi Prox
Epi Mid
Epi Dist
h
P35-S1
P47-S1
P64-S3
percentage
COL1A1
COL1A2
FBLN1
SERPINF1

**Extended Data Fig. 8 – Fibroblasts in the Human Breast**
**a,** Heatmap of scRNA-seq data clustered by fibroblast cell states showing top expressed genes for 100 randomly sampled cells per cluster, with canonical markers shown. **b,** Fibroblast cell state frequencies in scRNA-seq data and variation in 62 women. **c,** UMAP feature plots of top fibroblast cell type markers **d,** UMAP feature plots of top fibroblast cell state markers.
**e,** UMAP feature plots showing the expression of FAP in scRNA-seq data of fibroblasts. **f,** smFISH data showing fibroblast cell state markers in areas of connective tissue regions (I) and epithelial regions (II) from two women (P47-S1 and P64-S3). **g**, smFISH data (P35-S1) indicating spatial proximity regions with epithelial-proximal (Epi-prox), epithelial-middle (Epi-mid) and epithelial-distant (Epi-Dist) regions for different fibroblast cell state marker genes. **h,** Percentages of fibroblast cell state markers that are proximal, middle or distant to the epithelial cells, quantified from the smFISH data **i,** RNAscope *in situ* hybridization of breast tissues using an *MMP3* probe in combination with anti-Vimentin and anti-PanCK immunofluorescent staining, with enlarged panel below.

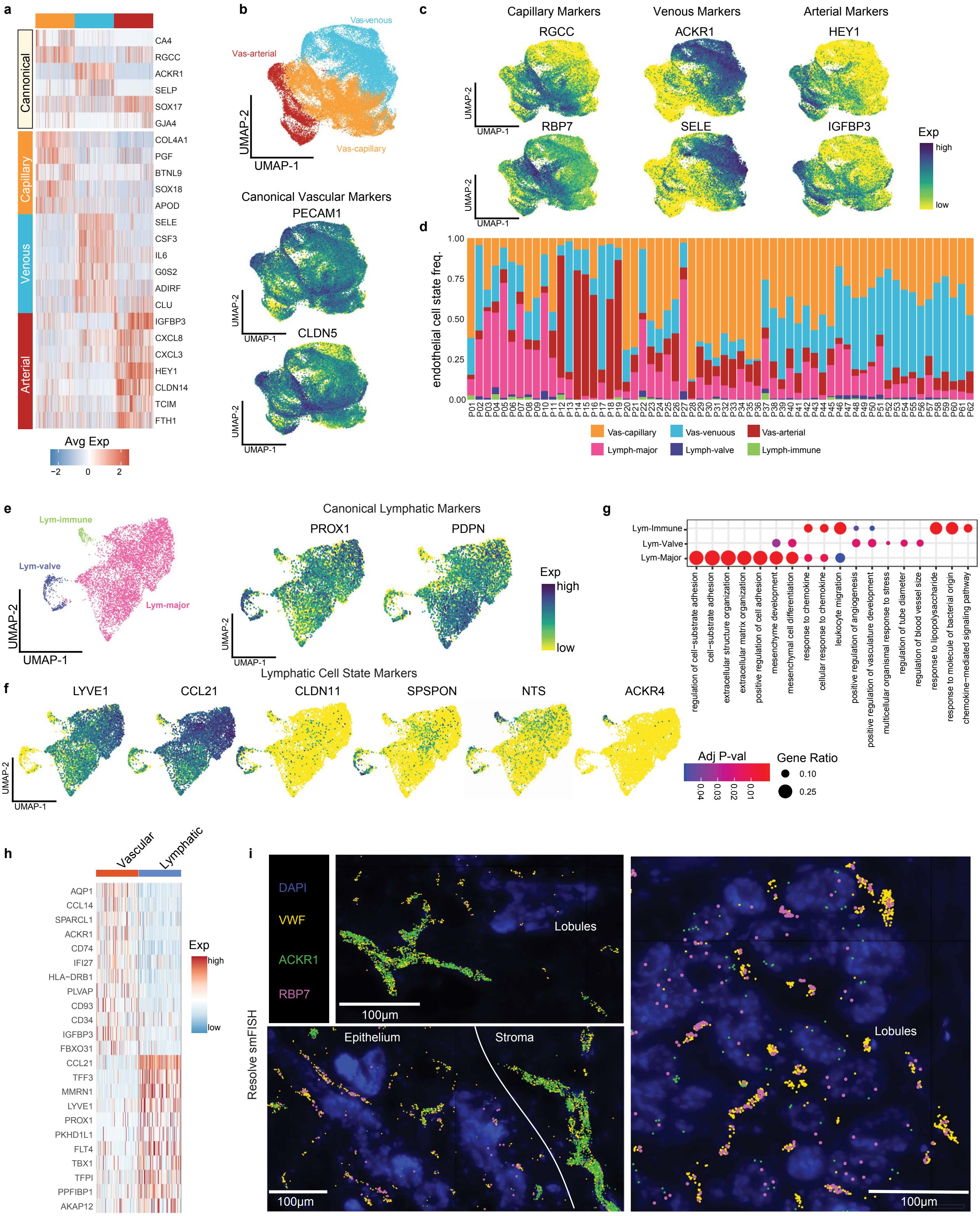
a
Cannonical
Capillary
Venous
Arterial
CA4
RGCC
ACKR1
SELP
SOX17
GJA4
COL4A1
PGF
BTNL9
SOX18
APOD
SELE
CSF3
IL6
G0S2
ADIRF
CLU
IGFBP3
CXCL8
CXCL3
HEY1
CLDN14
TCIM
FTH1
Avg Exp
−2
0
2
b
Vas-venous
Vas-arterial
Vas-capillary
UMAP-2
UMAP-1
Canonical Vascular Markers
PECAM1
CLDN5
c
Capillary Markers
Venous Markers
Arterial Markers
RGCC
ACKR1
HEY1
RBP7
SELE
IGFBP3
Exp
high
low
d
endothelial cell state freq.
1.00
0.75
0.50
0.25
0.00
Vas-capillary
Vas-venuous
Vas-arterial
Lymph-major
Lymph-valve
Lymph-immune
e
Lym-immune
Lym-valve
Lym-major
Canonical Lymphatic Markers
PROX1
PDPN
Lymphatic Cell State Markers
f
LYVE1
CCL21
CLDN11
SPSPON
NTS
ACKR4
g
Lym-Immune
Lym-Valve
Lym-Major
regulation of cell−substrate adhesion
cell−substrate adhesion
extracellular structure organization
extracellular matrix organization
positive regulation of cell adhesion
mesenchyme development
mesenchymal cell differentiation
response to chemokine
cellular response to chemokine
leukocyte migration
positive regulation of angiogenesis
positive regulation of vasculature development
multicellular organismal response to stress
regulation of tube diameter
regulation of blood vessel size
response to lipopolysaccharide
response to molecule of bacterial origin
chemokine−mediated signaling pathway
Adj P-val
0.04
0.03
0.02
0.01
Gene Ratio
0.10
0.25
h
Vascular
Lymphatic
AQP1
CCL14
SPARCL1
ACKR1
CD74
IFI27
HLA−DRB1
PLVAP
CD93
CD34
IGFBP3
FBXO31
CCL21
TFF3
MMRN1
LYVE1
PROX1
PKHD1L1
FLT4
TBX1
TFPI
PPFIBP1
AKAP12
i
Resolve smFISH
DAPI
VWF
ACKR1
RBP7
Lobules
100μm
Epithelium
Stroma

**Extended Data Fig. 9 – Endothelial cells in the Human Breast**

**a,** Heatmap of scRNA-seq data clustered by endothelial cell states, showing top expressed genes for 100 randomly sampled cells per cluster and canonical markers. **b,** UMAP of vascular cell state clusters with *PECAM1* and *CLDN5* canonical marker feature plots. **c,** UMAP feature plots of canonical markers for arterial, venous and capillary endothelial cell states. **d,** Endothelial cell state frequencies across 62 women. **e,** UMAP and feature plots showing the lymphatic cell state clusters and canonical marker genes **f,** UMAP feature plots showing lymphatic cell state marker gene expression. **g,** Dot plot of gene ontology enrichment results for lymphatic cell states. **h,** Heatmap showing top gene expression for vascular and lymphatic endothelial regions detected in the ST cluster data. **i,** smFISH data showing veins (*ACKR1*) and capillaries (*RBP7*), as well as a canonical vascular marker (*VWF*) in two different HBCA samples (P46-S3 and P64-S3).

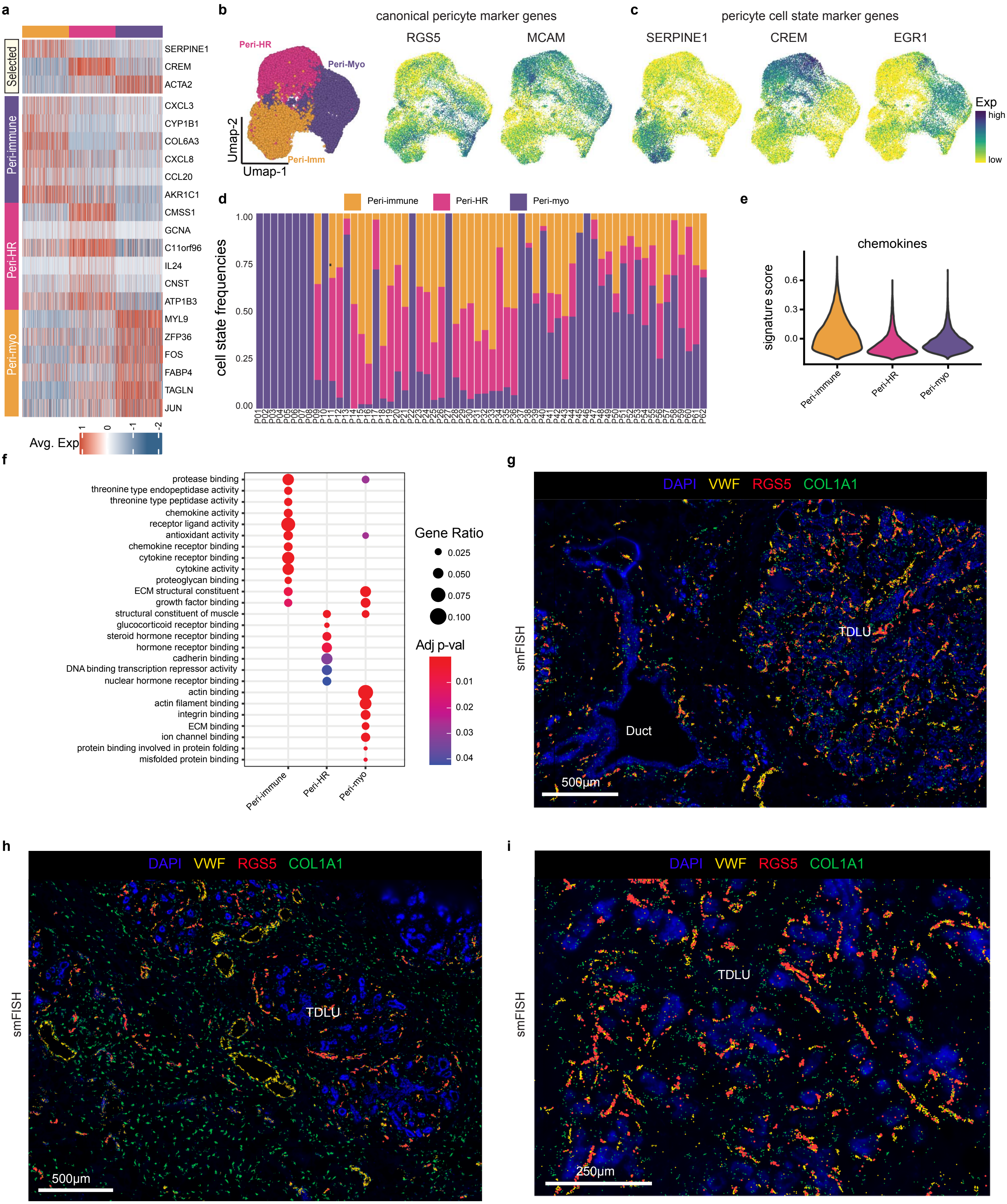
a
b
canonical pericyte marker genes
c
pericyte cell state marker genes
Selected
Peri-immune
Peri-HR
Peri-myo
SERPINE1
CREM
ACTA2
CXCL3
CYP1B1
COL6A3
CXCL8
CCL20
AKR1C1
CMSS1
GCNA
C11orf96
IL24
CNST
ATP1B3
MYL9
ZFP36
FOS
FABP4
TAGLN
JUN
Avg. Exp
Peri-HR
Peri-Myo
Peri-Imm
Umap-2
Umap-1
RGS5
MCAM
SERPINE1
CREM
EGR1
Exp
high
low
d
Peri-immune
Peri-HR
Peri-myo
cell state frequencies
e
chemokines
signature score
f
Gene Ratio
Adj p-val
protease binding
threonine type endopeptidase activity
threonine type peptidase activity
chemokine activity
receptor ligand activity
antioxidant activity
chemokine receptor binding
cytokine receptor binding
cytokine activity
proteoglycan binding
ECM structural constituent
growth factor binding
structural constituent of muscle
glucocorticoid receptor binding
steroid hormone receptor binding
hormone receptor binding
cadherin binding
DNA binding transcription repressor activity
nuclear hormone receptor binding
actin binding
actin filament binding
integrin binding
ECM binding
ion channel binding
protein binding involved in protein folding
misfolded protein binding
g
h
i
smFISH
DAPI VWF RGS5 COL1A1
TDLU
Duct
500µm
250µm

**Extended Data Fig. 10 – Pericytes in the Human Breast**
**a,** Heatmap of scRNA-seq data clustered by pericyte cell states showing top marker genes for 100 randomly sampled cells per cluster, with canonical markers above. **b,** UMAP feature plots showing the expression of canonical pericyte markers in the scRNA-seq data. **c**, UMAP feature plots showing pericyte cell state marker expression. **d,** Frequency of pericyte cell states and variation across the 62 women. **e,** Chemokine signature score across the pericyte cell states. **f,** Gene ontology enrichment results for the three pericyte cell states. **g-i,** smFISH data showing expression of pericyte marker *RGS5,* together with vascular marker *VWF* and fibroblast marker *COL1A1* in lobular and ductal regions from three different breast tissue samples (P46-S1, P47-S1 and P46-S3).

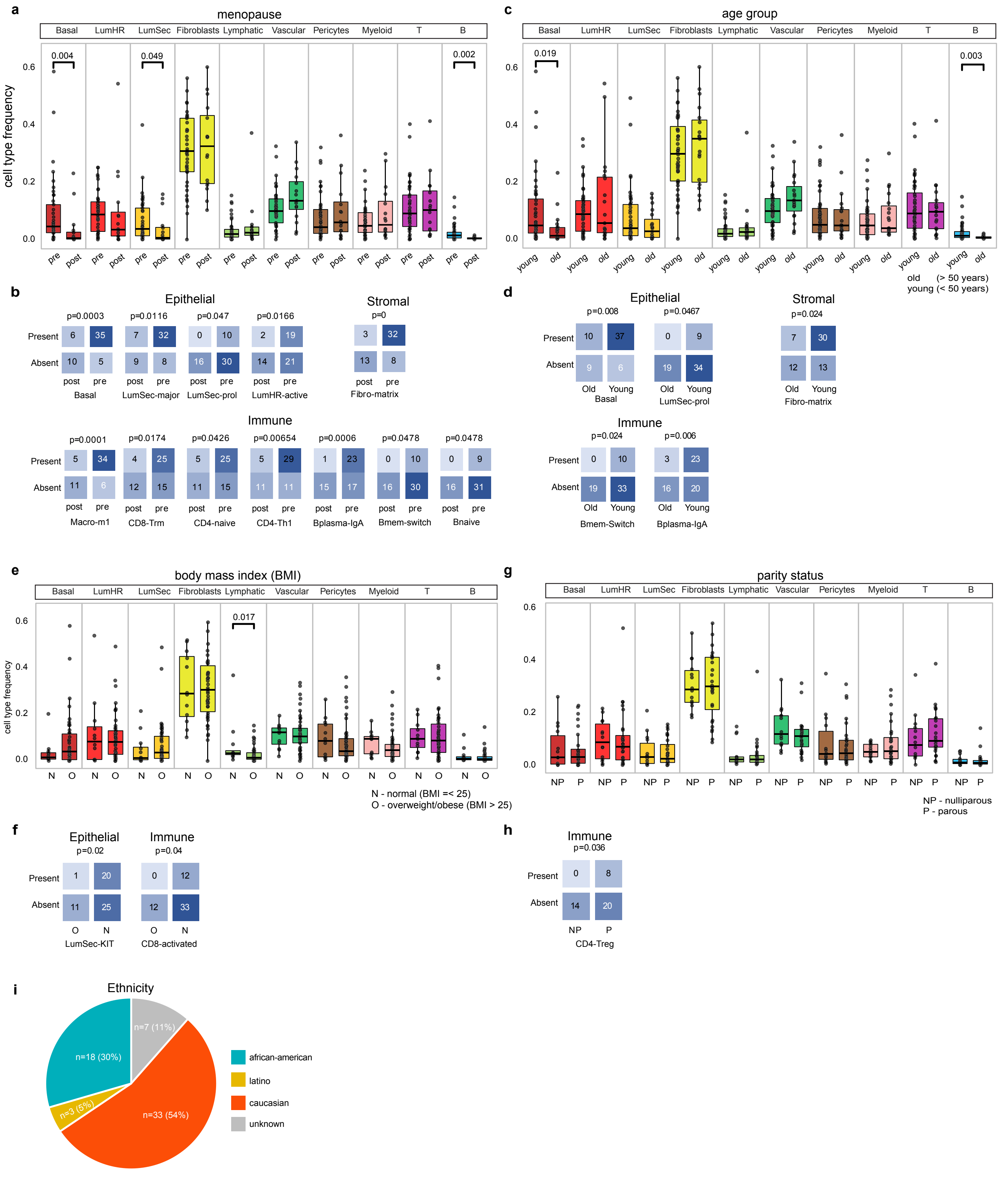
a
menopause
Basal LumHR LumSec Fibroblasts Lymphatic Vascular Pericytes Myeloid T B
cell type frequency
0.004
0.049
0.002
pre post
c
age group
0.019
0.003
young old
old (> 50 years)
young (< 50 years)
b
Epithelial
Stromal
p=0.0003 p=0.0116 p=0.047 p=0.0166 p=0
Present 6 35 7 32 0 10 2 19 3 32
Absent 10 5 9 8 16 30 14 21 13 8
post pre
Basal LumSec-major LumSec-prol LumHR-active Fibro-matrix
Immune
p=0.0001 p=0.0174 p=0.0426 p=0.00654 p=0.0006 p=0.0478 p=0.0478
Present 5 34 4 25 5 25 5 29 1 23 0 10 0 9
Absent 11 6 12 15 11 15 11 11 15 17 16 30 16 31
Macro-m1 CD8-Trm CD4-naive CD4-Th1 Bplasma-IgA Bmem-switch Bnaive
d
Epithelial
Stromal
p=0.008 p=0.0467 p=0.024
Present 10 37 0 9 7 30
Absent 9 6 19 34 12 13
Old Young
Basal LumSec-prol Fibro-matrix
Immune
p=0.024 p=0.006
Present 0 10 3 23
Absent 19 33 16 20
Bmem-Switch Bplasma-IgA
e
body mass index (BMI)
0.017
N O
N - normal (BMI =< 25)
O - overweight/obese (BMI > 25)
g
parity status
NP P
NP - nulliparous
P - parous
f
Epithelial p=0.02
Immune p=0.04
Present 1 20 0 12
Absent 11 25 12 33
O N
LumSec-KIT CD8-activated
h
Immune
p=0.036
Present 0 8
Absent 14 20
NP P
CD4-Treg
i
Ethnicity
n=7 (11%)
n=18 (30%)
n=3 (5%)
n=33 (54%)
african-american
latino
caucasian
unknown

**Extended Data Fig. 11 – Metadata correlations with breast cell types and states**
**a,** Association of the major cell type frequencies with pre- and post-menopause status in the n=62 women with significance using Wilcox test as indicated. **b,** Cell states significantly associated with menopause status determined by Fisher exact test. **c,** Association of major cell type frequencies with women's age classified into young (<50 years) and old (>50 years) groups for the n=62 women, with significance using Wilcox test indicated. **d,** Cell states significantly associated with age determined by Fisher exact test. **e,** Association of major cell type frequencies with BMI status classified as normal (BMI<25) and overweight/obese (BMI > 25) in the n=62 women with significance using Wilcox test indicated. **f,** Cell states significantly associated with BMI status determined by Fisher exact test. **g,** Association of major cell type frequencies with parity status (nulliparous, parous) in the n=62 women with significance using Wilcox test indicated. **h,** Cell states significantly associated with parity status determined by Fisher exact test. **i,** Pie chart showing distribution of ethnicity for the women in the breast atlas study.

### Cell Types and Cell States in the Adult Human Breast

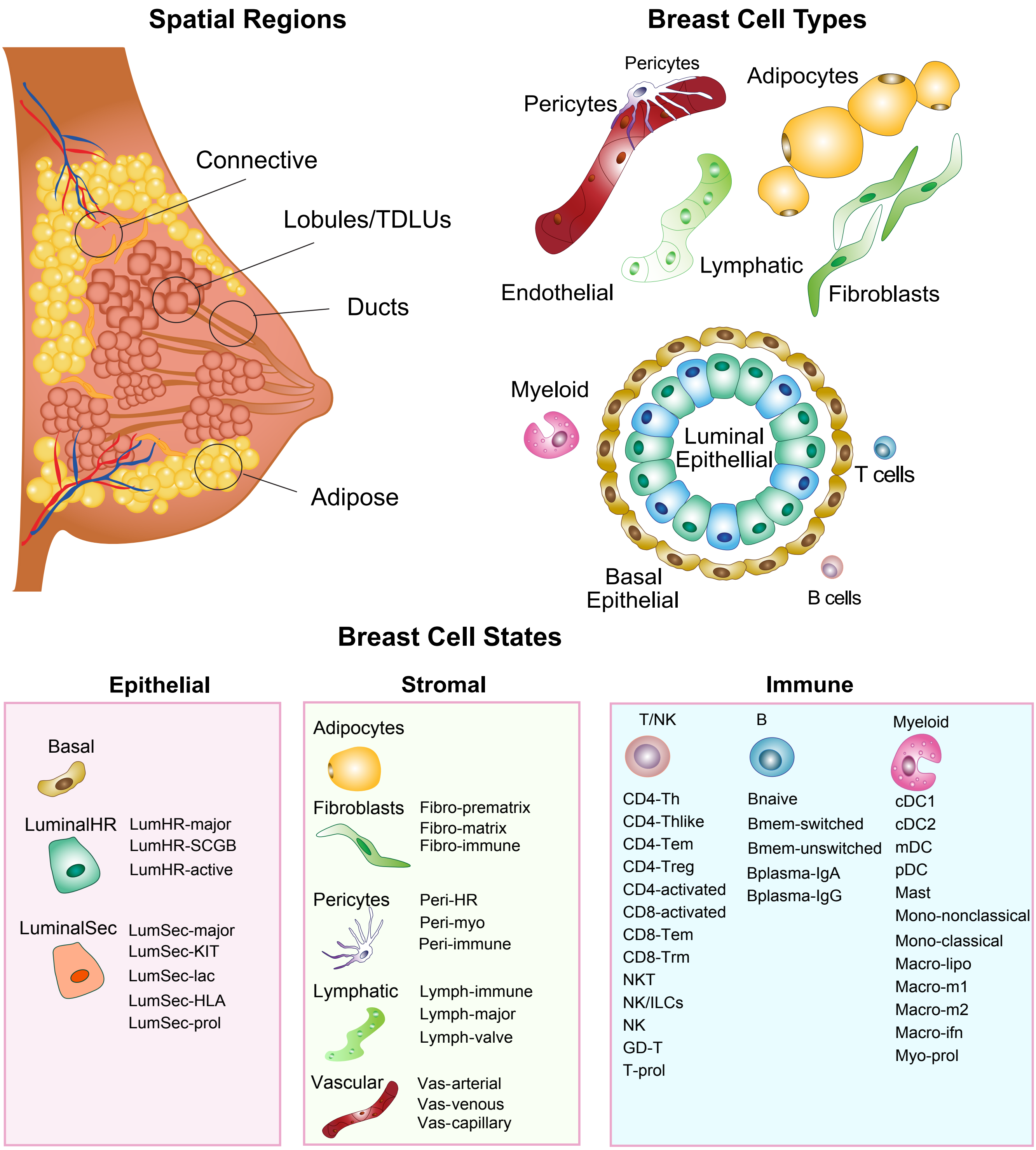

**Extended Data Fig. 12 – Summary of Cell Types in Cell States in Breast Tissues**
This figure summarizes all of the breast cell types and cell states that were identified in the HBCA study.

**Supplementary Table 1: Breast Atlas Tissue Samples and Clinical Metadata**

| Sample ID | Patient ID | Institution | Tissue Source | Age | Ethnicity | Parity | Menopause | BMI | BMI Group | Digestion Protocol (1hr/6hr/24hrs) |
|---|---|---|---|---|---|---|---|---|---|---|
| hbca_c01 | P1 | MDA | CM | 71 | white | 0 | post | 27.11 | Overweight | 1hr |
| hbca_c02 | P2 | MDA | CM | 36 | white | 1 | pre | 24.59 | Normal | 1hr |
| hbca_c03 | P3 | MDA | CM | 62 | white | unknown | post | 37.01 | Obese | 1hr |
| hbca_c04 | P4 | MDA | CM | 64 | white | 1 | post | 24.16 | Normal | 1hr |
| hbca_c05 | P5 | MDA | CM | 44 | white | 0 | pre | 22.89 | Normal | 1hr |
| hbca_c06 | P6 | MDA | CM | 44 | white | 1 | pre | 30.12 | Obese | 1hr |
| hbca_c07 | P7 | MDA | CM | 41 | white | 0 | pre | 27.69 | Overweight | 1hr |
| hbca_c08 | P8 | MDA | CM | 63 | white | 1 | post | 30.27 | Obese | 1hr |
| hbca_c09 | P9 | MDA | PM | 26 | white | unknown | pre | 23.09 | Normal | 1hr |
| hbca_c10 | P9 | MDA | PM | 26 | white | unknown | pre | 23.09 | Normal | 24hr |
| hbca_c11 | P10 | MDA | CM | 43 | white | 1 | pre | 29.79 | Overweight | 1hr |
| hbca_c12 | P11 | MDA | PM | 37 | hispanic | 0 | pre | 22.75 | Normal | 1hr |
| hbca_c13 | P11 | MDA | PM | 37 | hispanic | 0 | pre | 22.75 | Normal | 24hr |
| hbca_c14 | P12 | UCI | RM | 45 | white | unknown | unknown | 33.40 | Obese | 24hr |
| hbca_c15 | P12 | UCI | RM | 45 | white | unknown | unknown | 33.40 | Obese | 24hr |
| hbca_c16 | P13 | UCI | PM | unknown | unknown | 1 | post | unknown | unknown | 24hr |
| hbca_c17 | P13 | UCI | PM | unknown | unknown | 1 | post | unknown | unknown | 24hr |
| hbca_c18 | P14 | UCI | PM | 38 | unknown | unknown | unknown | unknown | unknown | 24hr |
| hbca_c19 | P15 | UCI | RM | 38 | white | unknown | pre | 28.70 | Overweight | 24hr |
| hbca_c20 | P16 | UCI | RM | 47 | white | unknown | unknown | 27.70 | Overweight | 24hr |
| hbca_c21 | P17 | UCI | PM | unknown | unknown | 0 | pre | unknown | unknown | 24hr |
| hbca_c22 | P18 | UCI | RM | 24 | white | unknown | pre | 32.40 | Obese | 24hr |
| hbca_c23 | P18 | UCI | RM | 24 | white | unknown | pre | 32.40 | Obese | 24hr |
| hbca_c24 | P19 | UCI | RM | 25 | white | unknown | pre | 32.20 | Obese | 24hr |
| hbca_c25 | P20 | MDA | RM | 46 | unknown | 1 | post | unknown | unknown | 1hr |
| hbca_c26 | P20 | MDA | RM | 46 | unknown | 1 | post | unknown | unknown | 24hr |
| hbca_c27 | P21 | MDA | CM | 31 | white | 1 | pre | 24.45 | Normal | 24hr |
| hbca_c28 | P22 | MDA | CM | 35 | white | 0 | pre | 29.52 | Overweight | 1hr |
| hbca_c29 | P23 | MDA | CM | 40 | white | 1 | pre | 27.92 | Overweight | 1hr |
| hbca_c30 | P23 | MDA | CM | 40 | white | 1 | pre | 27.92 | Overweight | 24hr |
| hbca_c31 | P24 | MDA | RM | unknown | unknown | unknown | unknown | unknown | unknown | 6hr |
| hbca_c32 | P24 | MDA | RM | unknown | unknown | unknown | unknown | unknown | unknown | 24hr |
| hbca_c33 | P25 | MDA | CM | 63 | black | unknown | post | 25.43 | Overweight | 24hr |
| hbca_c34 | P26 | MDA | CM | 45 | white | 1 | pre | 37.07 | Obese | 24hr |
| hbca_c35 | P27 | MDA | CM | 62 | white | 1 | post | 22.97 | Normal | 1hr |
| hbca_c36 | P28 | MDA | CM | 65 | white | unknown | post | 32.29 | Obese | 24hr |
| hbca_c37 | P29 | MDA | CM | 56 | white | unknown | post | 38.43 | Obese | 24hr |
| hbca_c38 | P30 | MDA | PM | 47 | white | unknown | post | 25.64 | Overweight | 24hr |
| hbca_c39 | P31 | MDA | CM | 61 | white | 0 | post | 24.19 | Normal | 24hr |
| hbca_c40 | P37 | MDA | CM | 68 | white | unknown | post | 25.45 | Overweight | 1hr |
| hbca_c41 | P32 | MDA | CM | 42 | white | 1 | pre | 26.98 | Overweight | 24hr |
| hbca_c42 | P33 | MDA | CM | 38 | white | 1 | pre | 26.76 | Overweight | 24hr |
| hbca_c43 | P34 | MDA | CM | 36 | white | 1 | pre | 20.80 | Normal | 24hr |
| hbca_c44 | P35 | MDA | CM | 46 | white | 1 | pre | 24.71 | Normal | 24hr |
| hbca_c45 | P36 | MDA | CM | 43 | black | 0 | pre | 33.84 | Obese | 24hr |
| hbca_c46 | P42 | MDA | CM | 29 | unknown | 1 | pre | 29.27 | Overweight | 1hr |
| hbca_c47 | P43 | MDA | CM | 45 | unknown | 1 | pre | 22.66 | Normal | 1hr |
| hbca_c48 | P43 | MDA | CM | 45 | unknown | 1 | pre | 22.66 | Normal | 6hr |
| hbca_c49 | P43 | MDA | CM | 45 | unknown | 1 | pre | 22.66 | Normal | 24hr |
| hbca_c50 | P44 | BCM | RM | 51 | white | 0 | post | 29.30 | Overweight | 24hr |
| hbca_c51 | P44 | BCM | RM | 51 | white | 0 | post | 29.30 | Overweight | 24hr |
| hbca_c52 | P45 | BCM | RM | 26 | white | 1 | pre | 25.06 | Overweight | 6hr |
| hbca_c53 | P46 | BCM | RM | 34 | white | 1 | pre | 42.51 | Obese | 6hr |
| hbca_c54 | P46 | BCM | RM | 34 | white | 1 | pre | 42.51 | Obese | 6hr |
| hbca_c55 | P47 | BCM | RM | 49 | black | 1 | pre | 29.79 | Overweight | 6hr |
| hbca_c56 | P47 | BCM | RM | 49 | black | 1 | pre | 29.79 | Overweight | 6hr |
| hbca_c57 | P48 | BCM | RM | 38 | black | 1 | pre | 34.70 | Obese | 6hr |
| hbca_c58 | P48 | BCM | RM | 38 | black | 1 | pre | 34.70 | Obese | 6hr |
| hbca_c59 | P49 | BCM | RM | 36 | black | 1 | pre | 31.37 | Obese | 6hr |
| hbca_c60 | P49 | BCM | RM | 36 | black | 1 | pre | 31.37 | Obese | 6hr |
| hbca_c61 | P49 | BCM | RM | 36 | black | 1 | pre | 31.37 | Obese | 6hr |
| hbca_c62 | P50 | BCM | RM | 26 | black | 0 | pre | 31.18 | Obese | 6hr |
| hbca_c63 | P50 | BCM | RM | 26 | black | 0 | pre | 31.18 | Obese | 6hr |
| hbca_c64 | P50 | BCM | RM | 26 | black | 0 | pre | 31.18 | Obese | 6hr |
| hbca_c65 | P50 | BCM | RM | 26 | black | 0 | pre | 31.18 | Obese | 6hr |
| hbca_c66 | P51 | BCM | RM | 29 | black | unknown | pre | 30.61 | Obese | 6hr |

| Sample ID | Patient ID | Institution | Tissue Source | Age | Ethnicity | Parity | Menopause | BMI | BMI Group | Digestion Protocol (1hr/6hr/24hrs) |
|---|---|---|---|---|---|---|---|---|---|---|
| hbca_c67 | P51 | BCM | RM | 29 | black | unknown | pre | 30.61 | Obese | 6hr |
| hbca_c68 | P51 | BCM | RM | 29 | black | unknown | pre | 30.61 | Obese | 6hr |
| hbca_c69 | P51 | BCM | RM | 29 | black | unknown | pre | 30.61 | Obese | 6hr |
| hbca_c70 | P52 | BCM | RM | 45 | black | 1 | pre | 39.89 | Obese | 6hr |
| hbca_c71 | P52 | BCM | RM | 45 | black | 1 | pre | 39.89 | Obese | 6hr |
| hbca_c72 | P53 | BCM | RM | 51 | black | 1 | pre | 24.93 | normal | 6hr |
| hbca_c73 | P53 | BCM | RM | 51 | black | 1 | pre | 24.93 | normal | 6hr |
| hbca_c74 | P54 | BCM | RM | 30 | black | 1 | pre | 35.92 | Obese | 6hr |
| hbca_c75 | P54 | BCM | RM | 30 | black | 1 | pre | 35.92 | Obese | 6hr |
| hbca_c76 | P55 | BCM | RM | 37 | white | 0 | pre | 28.13 | Overweight | 6hr |
| hbca_c77 | P55 | BCM | RM | 37 | white | 0 | pre | 28.13 | Overweight | 6hr |
| hbca_c78 | P56 | BCM | RM | 18 | black | 0 | pre | 34.00 | Obese | 6hr |
| hbca_c79 | P56 | BCM | RM | 18 | black | 0 | pre | 34.00 | Obese | 6hr |
| hbca_c80 | P57 | BCM | RM | 38 | black | 1 | pre | 43.95 | Obese | 6hr |
| hbca_c81 | P57 | BCM | RM | 38 | black | 1 | pre | 43.95 | Obese | 6hr |
| hbca_c82 | P38 | BCM | RM | 54 | white | 1 | post | 27.64 | Overweight | 6hr |
| hbca_c83 | P38 | BCM | RM | 54 | white | 1 | post | 27.64 | Overweight | 6hr |
| hbca_c84 | P39 | BCM | RM | 19 | black | unknown | pre | 30.79 | Obese | 6hr |
| hbca_c85 | P39 | BCM | RM | 19 | black | unknown | pre | 30.79 | Obese | 6hr |
| hbca_c86 | P40 | BCM | RM | 29 | black | 0 | pre | 38.09 | Obese | 6hr |
| hbca_c87 | P40 | BCM | RM | 29 | black | 0 | pre | 38.09 | Obese | 6hr |
| hbca_c88 | P58 | BCM | RM | 24 | black | 0 | pre | 35.19 | Obese | 6hr |
| hbca_c89 | P58 | BCM | RM | 24 | black | 0 | pre | 35.19 | Obese | 6hr |
| hbca_c90 | P41 | BCM | RM | 23 | black | 1 | pre | 34.41 | Obese | 6hr |
| hbca_c91 | P41 | BCM | RM | 23 | black | 1 | pre | 34.41 | Obese | 6hr |
| hbca_c92 | P59 | BCM | RM | 63 | hispanic | unknown | post | 31.50 | Obese | 6hr |
| hbca_c93 | P59 | BCM | RM | 63 | hispanic | unknown | post | 31.50 | Obese | 6hr |
| hbca_c94 | P60 | BCM | RM | 52 | hispanic | unknown | unknown | 39.00 | Obese | 6hr |
| hbca_c95 | P60 | BCM | RM | 52 | hispanic | unknown | unknown | 39.00 | Obese | 6hr |
| hbca_c96 | P61 | BCM | RM | 50 | black | unknown | unknown | 42.16 | Obese | 6hr |
| hbca_c97 | P61 | BCM | RM | 50 | black | unknown | unknown | 42.16 | Obese | 6hr |
| hbca_c98 | P62 | BCM | RM | 26 | black | unknown | pre | 38.04 | Obese | 6hr |
| hbca_c99 | P62 | BCM | RM | 26 | black | unknown | pre | 38.04 | Obese | 6hr |
| hbca_c100 | P59 | BCM | RM | 63 | hispanic | unknown | post | 31.50 | Obese | 6hr |

**Supplementary Table 1: Breast Atlas Tissue Samples and Clinical Metadata**
This table lists the n=100 tissue samples and clinical metadata for the 62 women included in the breast atlas project. The columns list from left to right the sample identifier (Sample ID), the patient identifier (Patient ID), the institution where the tissue sample was collected from MD Anderson (MDA), UC Irvine (UCI) or Baylor College of Medicine (BCM), the procedure from which the tissue source was collected reduction mammoplasties (RM=30), prophylactic mastectomies (PM=6), and contralateral mastectomies (CM=26), the age of the women, the ethnicity the women self-identifies as, the parity status as positive (1), negative (0) or unknown, the menopause status (pre/post), the Body Mass Index (BMI), the BMI Group classification and digestion protocol time use during sample dissociation (1hr, 6hr, or 24hr).

**Supplementary Table 2: Single Cell RNA-seq Quality Control Metrics**

| Sample ID | Number of Cells | Mean Reads per Cell | Median Genes per Cell | Number of Reads | Fraction Reads in Cells | Genes Detected | Median UMI Counts per Cell |
|---|---|---|---|---|---|---|---|
| hbca_c01 | 4,091 | 79,318 | 353 | 324,492,736 | 75.10% | 20,828 | 1,981 |
| hbca_c02 | 9,577 | 36,403 | 923 | 348,639,606 | 72.30% | 23,775 | 2,336 |
| hbca_c03 | 5,450 | 65,569 | 1,032 | 357,352,385 | 70.60% | 22,024 | 2,564 |
| hbca_c04 | 8,924 | 39,111 | 800 | 349,034,669 | 69.80% | 23,140 | 1,705 |
| hbca_c05 | 5,786 | 57,183 | 709 | 330,866,587 | 70.10% | 22,209 | 1,604 |
| hbca_c06 | 3,593 | 89,184 | 1,178 | 320,441,257 | 83.50% | 22,069 | 2,948 |
| hbca_c07 | 5,250 | 60,928 | 953 | 319,872,861 | 76.80% | 22,364 | 2,250 |
| hbca_c08 | 2,533 | 137,485 | 1,055 | 348,249,946 | 69.50% | 20,626 | 2,570 |
| hbca_c09 | 7,894 | 42,387 | 823 | 334,609,974 | 71.80% | 23,308 | 1,922 |
| hbca_c10 | 5,682 | 63,430 | 771 | 360,412,188 | 87.90% | 21,623 | 1,645 |
| hbca_c11 | 4,052 | 84,470 | 787 | 342,274,452 | 76.60% | 21,264 | 1,729 |
| hbca_c12 | 2,322 | 141,601 | 872 | 328,799,276 | 55.00% | 20,459 | 2,177 |
| hbca_c13 | 4,127 | 83,670 | 1,099 | 345,309,344 | 86.80% | 21,564 | 2,690 |
| hbca_c25 | 6,401 | 59,231 | 2,582 | 379,140,265 | 84.70% | 24,813 | 7,353 |
| hbca_c26 | 7,845 | 50,538 | 2,020 | 396,476,992 | 90.10% | 25,280 | 6,664 |
| hbca_c27 | 5,944 | 59,127 | 1,755 | 351,455,269 | 78.10% | 23,853 | 5,445 |
| hbca_c28 | 2,526 | 154,806 | 1,449 | 391,041,072 | 70.40% | 22,713 | 4,255 |
| hbca_c29 | 6,425 | 59,692 | 1,509 | 383,526,635 | 74.70% | 24,811 | 4,744 |
| hbca_c30 | 12,399 | 30,024 | 1,654 | 372,270,308 | 78.70% | 25,997 | 4,849 |
| hbca_c31 | 7,134 | 44,541 | 2,003 | 317,759,854 | 85.30% | 24,105 | 6,488 |
| hbca_c32 | 11,034 | 32,093 | 1,506 | 354,120,495 | 81.50% | 25,021 | 4,758 |
| hbca_c33 | 6,839 | 58,653 | 2,054 | 401,131,543 | 88.30% | 24,780 | 7,035 |
| hbca_c34 | 8,930 | 55,014 | 1,918 | 491,282,666 | 87.30% | 25,091 | 6,060 |
| hbca_c35 | 5,694 | 76,943 | 1,203 | 438,114,228 | 55.10% | 23,846 | 3,606 |
| hbca_c36 | 7,582 | 58,284 | 2,382 | 441,911,150 | 92.10% | 23,845 | 7,912 |
| hbca_c37 | 10,318 | 44,250 | 1,460 | 456,573,803 | 85.50% | 25,256 | 5,031 |
| hbca_c38 | 8,639 | 43,128 | 2,012 | 372,584,898 | 88.20% | 24,854 | 5,860 |
| hbca_c39 | 5,917 | 66,216 | 2,055 | 391,801,107 | 85.90% | 24,007 | 7,619 |
| hbca_c40 | 4,570 | 90,273 | 1,377 | 412,548,145 | 78.40% | 23,685 | 3,944 |
| hbca_c41 | 8,420 | 47,239 | 1,529 | 397,758,089 | 85.60% | 25,743 | 4,501 |
| hbca_c42 | 6,864 | 56,666 | 2,001 | 388,956,766 | 86.00% | 25,564 | 5,956 |
| hbca_c43 | 10,061 | 39,595 | 2,277 | 398,365,868 | 90.40% | 25,700 | 7,654 |
| hbca_c44 | 12,386 | 37,439 | 1,850 | 463,722,721 | 87.80% | 25,490 | 6,745 |
| hbca_c45 | 16,520 | 25,592 | 1,183 | 422,790,667 | 74.80% | 25,474 | 2,830 |
| hbca_c46 | 11,913 | 34,346 | 1,207 | 409,172,629 | 71.50% | 25,136 | 2,945 |
| hbca_c47 | 9,126 | 39,254 | 1,125 | 358,234,249 | 70.80% | 24,276 | 2,895 |
| hbca_c48 | 8,627 | 46,106 | 1,391 | 397,758,601 | 87.00% | 25,214 | 4,322 |
| hbca_c49 | 8,126 | 58,187 | 2,040 | 472,831,839 | 89.70% | 24,266 | 6,953 |
| hbca_c50 | 5,018 | 87,632 | 1,703 | 439,739,114 | 89.70% | 26,109 | 5,064 |
| hbca_c51 | 6,129 | 76,466 | 2,255 | 468,665,048 | 90.60% | 25,517 | 7,893 |
| hbca_c52 | 5,805 | 82,968 | 808 | 481,631,031 | 59.30% | 23,909 | 2,252 |
| hbca_c53 | 10,090 | 44,704 | 1,337 | 451,065,448 | 86.70% | 25,902 | 3,777 |
| hbca_c54 | 8,900 | 53,864 | 1,946 | 479,396,431 | 87.50% | 26,149 | 5,733 |
| hbca_c55 | 11,025 | 33,129 | 1,976 | 365,251,856 | 88.40% | 25,648 | 6,164 |
| hbca_c56 | 13,292 | 27,174 | 1,745 | 361,197,190 | 86.40% | 25,562 | 5,157 |
| hbca_c57 | 12,807 | 28,873 | 1,244 | 369,783,072 | 82.10% | 25,986 | 3,855 |
| hbca_c58 | 12,035 | 37,459 | 1,444 | 450,819,388 | 85.10% | 25,941 | 4,758 |
| hbca_c59 | 18,664 | 21,815 | 1,582 | 407,158,167 | 87.60% | 26,701 | 4,431 |
| hbca_c60 | 12,740 | 29,815 | 1,730 | 379,843,424 | 87.20% | 26,172 | 5,275 |
| hbca_c61 | 9,448 | 42,433 | 1,160 | 400,915,804 | 71.40% | 24,937 | 3,589 |
| hbca_c62 | 11,795 | 29,324 | 1,706 | 345,879,461 | 85.10% | 25,732 | 4,892 |

| Sample ID | Number of Cells | Mean Reads per Cell | Median Genes per Cell | Number of Reads | Fraction Reads in Cells | Genes Detected | Median UMI Counts per Cell |
|---|---|---|---|---|---|---|---|
| hbca_c63 | 11,648 | 32,884 | 1,715 | 383,040,911 | 84.00% | 25,098 | 5,211 |
| hbca_c64 | 10,615 | 46,004 | 2,228 | 488,341,054 | 84.70% | 25,395 | 7,493 |
| hbca_c65 | 13,521 | 31,221 | 1,864 | 422,151,595 | 82.80% | 25,085 | 6,331 |
| hbca_c66 | 13,547 | 31,444 | 1,608 | 425,972,057 | 81.80% | 25,651 | 4,667 |
| hbca_c67 | 10,884 | 44,418 | 1,747 | 483,454,528 | 83.70% | 25,936 | 5,098 |
| hbca_c68 | 15,150 | 31,424 | 1,619 | 476,074,866 | 81.40% | 25,869 | 4,586 |
| hbca_c69 | 10,582 | 42,935 | 1,566 | 454,341,839 | 78.70% | 26,038 | 4,470 |
| hbca_c70 | 9,916 | 39,706 | 2,072 | 393,728,950 | 87.30% | 25,873 | 6,800 |
| hbca_c71 | 11,964 | 32,298 | 1,887 | 386,422,651 | 86.10% | 25,966 | 5,908 |
| hbca_c72 | 11,502 | 32,651 | 1,974 | 375,557,112 | 83.20% | 26,308 | 5,548 |
| hbca_c73 | 11,647 | 32,187 | 2,044 | 374,889,239 | 85.40% | 26,033 | 6,289 |
| hbca_c74 | 13,610 | 29,180 | 1,643 | 397,144,206 | 85.50% | 25,747 | 4,829 |
| hbca_c75 | 16,502 | 21,757 | 1,373 | 359,038,086 | 83.90% | 25,739 | 3,648 |
| hbca_c76 | 11,679 | 36,674 | 2,152 | 428,317,510 | 85.70% | 26,425 | 6,040 |
| hbca_c77 | 11,403 | 36,082 | 2,210 | 411,454,267 | 87.80% | 26,681 | 6,083 |
| hbca_c78 | 12,214 | 28,491 | 1,775 | 347,997,546 | 80.30% | 25,107 | 5,101 |
| hbca_c79 | 12,305 | 29,823 | 1,654 | 366,983,410 | 76.00% | 25,296 | 4,695 |
| hbca_c80 | 13,289 | 28,285 | 1,563 | 375,892,640 | 81.70% | 25,546 | 4,511 |
| hbca_c81 | 13,122 | 28,146 | 1,645 | 369,333,714 | 83.40% | 25,945 | 4,718 |
| hbca_c82 | 7,533 | 42,551 | 1,564 | 320,540,361 | 77.10% | 24,934 | 4,602 |
| hbca_c83 | 9,969 | 36,167 | 1,336 | 360,555,401 | 75.70% | 25,155 | 3,819 |
| hbca_c84 | 12,466 | 38,226 | 1,372 | 476,527,092 | 72.80% | 25,280 | 3,941 |
| hbca_c85 | 11,774 | 41,479 | 1,700 | 488,380,226 | 78.70% | 25,568 | 5,555 |
| hbca_c86 | 6,133 | 58,994 | 1,322 | 361,813,963 | 80.10% | 28,001 | 2,090 |
| hbca_c87 | 6,947 | 47,397 | 1,327 | 329,269,846 | 84.30% | 28,492 | 2,176 |
| hbca_c88 | 7,041 | 58,303 | 1,985 | 410,512,392 | 86.40% | 24,851 | 6,021 |
| hbca_c89 | 6,718 | 57,121 | 2,108 | 383,742,211 | 87.90% | 24,654 | 6,480 |
| hbca_c90 | 9,161 | 44,766 | 1,635 | 410,105,314 | 87.90% | 24,367 | 4,604 |
| hbca_c91 | 9,469 | 28,450 | 1,403 | 269,397,441 | 87.60% | 24,043 | 3,557 |
| hbca_c92 | 8,476 | 51,364 | 2,060 | 435,365,199 | 84.00% | 26,145 | 6,471 |
| hbca_c93 | 6,618 | 62,898 | 2,086 | 416,264,504 | 84.30% | 25,696 | 6,561 |
| hbca_c94 | 8,431 | 44,139 | 2,135 | 372,138,533 | 84.10% | 24,879 | 6,451 |
| hbca_c95 | 8,086 | 42,646 | 1,720 | 344,840,516 | 76.80% | 24,895 | 5,008 |
| hbca_c96 | 10,184 | 27,403 | 1,499 | 279,075,635 | 84.60% | 24,581 | 4,325 |
| hbca_c97 | 8,411 | 38,493 | 1,855 | 323,768,135 | 86.00% | 25,019 | 5,764 |
| hbca_c98 | 7,145 | 55,077 | 2,082 | 393,531,047 | 86.30% | 25,265 | 6,755 |
| hbca_c99 | 7,950 | 50,287 | 2,220 | 399,788,791 | 87.60% | 25,471 | 7,480 |
| hbca_c100 | 8,927 | 50,041 | 2,485 | 446,716,152 | 84.20% | 25,675 | 8,324 |

**Supplementary Table 2: Single Cell RNA-seq Quality Control Metrics**

This table lists the quality control metrics for the fresh breast tissue samples sequenced with single cell RNA sequencing using the 10X Genomics Chromium Platform before any filtering was applied. The columns listed from left to right indicate the Sample ID, Number of Cells, Mean Reads per Cell, Median Genes per Cell, Number of Reads, Fraction Reads in Cells, Total Genes Detected, Median UMI Counts per Cell.

#### Supplementary Table 3: Top Marker Genes Expressed in Major Breast Cell Types

| Cell Type | Gene | Average LogFC | pct1 | pct2 |
|---|---|---|---|---|
| Basal | KRT14 | 3.834 | 0.936 | 0.120 |
| | KRT17 | 3.568 | 0.938 | 0.156 |
| | SAA1 | 2.778 | 0.807 | 0.190 |
| | KRT5 | 2.527 | 0.876 | 0.057 |
| | DST | 2.446 | 0.895 | 0.403 |
| | ACTA2 | 2.235 | 0.783 | 0.134 |
| | TAGLN | 2.196 | 0.895 | 0.265 |
| | SFN | 2.061 | 0.803 | 0.111 |
| | MT1X | 1.996 | 0.834 | 0.464 |
| | ACTG2 | 1.963 | 0.628 | 0.022 |
| | MYLK | 1.890 | 0.709 | 0.093 |
| | TPM2 | 1.748 | 0.917 | 0.329 |
| | KRT6B | 1.611 | 0.490 | 0.057 |
| | FBXO32 | 1.543 | 0.623 | 0.131 |
| | S100A2 | 1.506 | 0.354 | 0.044 |
| | MT2A | 1.488 | 0.972 | 0.814 |
| | CNN1 | 1.425 | 0.569 | 0.060 |
| | TPM1 | 1.409 | 0.868 | 0.434 |
| | FBXO2 | 1.291 | 0.508 | 0.061 |
| | CRYAB | 1.290 | 0.758 | 0.250 |
| LumHR | AREG | 3.000 | 0.783 | 0.206 |
| | MUCL1 | 2.801 | 0.371 | 0.118 |
| | AZGP1 | 2.702 | 0.927 | 0.174 |
| | ANKRD30A | 2.471 | 0.765 | 0.023 |
| | KRT18 | 2.319 | 0.941 | 0.218 |
| | PIP | 2.274 | 0.354 | 0.069 |
| | KRT8 | 2.000 | 0.916 | 0.218 |
| | KRT19 | 1.904 | 0.866 | 0.191 |
| | S100A14 | 1.840 | 0.842 | 0.125 |
| | AGR2 | 1.759 | 0.588 | 0.020 |
| | TFF1 | 1.741 | 0.314 | 0.018 |
| | TCIM | 1.716 | 0.740 | 0.129 |
| | CD24 | 1.659 | 0.873 | 0.191 |
| | STC2 | 1.554 | 0.677 | 0.071 |
| | SYTL2 | 1.525 | 0.678 | 0.060 |
| | SPINT2 | 1.486 | 0.906 | 0.215 |
| | ELF3 | 1.454 | 0.803 | 0.123 |
| | CLDN4 | 1.386 | 0.841 | 0.169 |
| | AGR3 | 1.381 | 0.481 | 0.011 |
| | CLDN7 | 1.304 | 0.804 | 0.103 |
| LumSec | WFDC2 | 2.516 | 0.576 | 0.189 |
| | SCGB2A2 | 2.425 | 0.395 | 0.106 |
| | PI3 | 2.202 | 0.288 | 0.030 |
| | FDCSP | 2.199 | 0.174 | 0.046 |
| | SLPI | 2.195 | 0.719 | 0.098 |
| | LTF | 2.176 | 0.559 | 0.077 |
| | KRT15 | 2.146 | 0.715 | 0.101 |
| | S100A9 | 1.907 | 0.477 | 0.069 |
| | MMP7 | 1.783 | 0.496 | 0.053 |
| | CLDN4 | 1.733 | 0.802 | 0.186 |
| | SCGB3A1 | 1.679 | 0.351 | 0.094 |
| | KRT19 | 1.644 | 0.822 | 0.208 |
| | KRT7 | 1.582 | 0.870 | 0.277 |
| | KRT23 | 1.564 | 0.561 | 0.057 |
| | CD24 | 1.509 | 0.851 | 0.207 |
| | S100A8 | 1.493 | 0.191 | 0.032 |
| | TACSTD2 | 1.490 | 0.832 | 0.305 |
| | CCL28 | 1.470 | 0.643 | 0.042 |
| | PIGR | 1.452 | 0.555 | 0.035 |
| | RARRES1 | 1.405 | 0.432 | 0.118 |

| Cell Type | Gene | Average LogFC | pct1 | pct2 |
|---|---|---|---|---|
| Vascular | SELE | 2.432 | 0.341 | 0.020 |
| | FABP4 | 2.421 | 0.631 | 0.150 |
| | ACKR1 | 2.361 | 0.435 | 0.018 |
| | CSF3 | 2.139 | 0.405 | 0.059 |
| | CLDN5 | 2.096 | 0.689 | 0.047 |
| | STC1 | 2.065 | 0.476 | 0.054 |
| | ANGPT2 | 1.946 | 0.629 | 0.075 |
| | TM4SF1 | 1.946 | 0.930 | 0.456 |
| | SPARCL1 | 1.853 | 0.823 | 0.366 |
| | ADAMTS9 | 1.812 | 0.665 | 0.079 |
| | IFI27 | 1.781 | 0.853 | 0.296 |
| | SERPINE1 | 1.751 | 0.584 | 0.274 |
| | ADGRL4 | 1.729 | 0.719 | 0.014 |
| | C2CD4B | 1.712 | 0.436 | 0.033 |
| | RBP7 | 1.709 | 0.499 | 0.060 |
| | GNG11 | 1.653 | 0.807 | 0.227 |
| | PECAM1 | 1.615 | 0.664 | 0.043 |
| | CD93 | 1.585 | 0.673 | 0.043 |
| | PGF | 1.576 | 0.337 | 0.085 |
| | AQP1 | 1.560 | 0.383 | 0.039 |
| Pericytes | C11orf96 | 2.633 | 0.896 | 0.321 |
| | RGS5 | 2.159 | 0.494 | 0.014 |
| | MT1A | 2.059 | 0.754 | 0.270 |
| | CCL20 | 1.588 | 0.143 | 0.046 |
| | PDK4 | 1.563 | 0.590 | 0.239 |
| | ADAMTS4 | 1.499 | 0.653 | 0.206 |
| | STEAP4 | 1.449 | 0.355 | 0.069 |
| | MYL9 | 1.442 | 0.584 | 0.307 |
| | EDNRB | 1.434 | 0.576 | 0.203 |
| | ADAMTS1 | 1.414 | 0.690 | 0.262 |
| | IGFBP7 | 1.408 | 0.940 | 0.460 |
| | TAGLN | 1.352 | 0.683 | 0.308 |
| | IL6 | 1.342 | 0.448 | 0.187 |
| | IGFBP5 | 1.313 | 0.534 | 0.210 |
| | MCAM | 1.300 | 0.551 | 0.092 |
| | ADIRF | 1.278 | 0.751 | 0.461 |
| | NR2F2 | 1.254 | 0.578 | 0.216 |
| | MYH11 | 1.252 | 0.311 | 0.050 |
| | GJA4 | 1.231 | 0.402 | 0.008 |
| | CRYAB | 1.226 | 0.561 | 0.286 |
| B cells | IGKC | 5.473 | 0.846 | 0.354 |
| | IGLC2 | 5.200 | 0.486 | 0.118 |
| | IGHA1 | 5.151 | 0.737 | 0.272 |
| | IGLC3 | 4.987 | 0.403 | 0.088 |
| | JCHAIN | 4.889 | 0.643 | 0.090 |
| | IGHA2 | 4.357 | 0.479 | 0.078 |
| | IGHM | 4.093 | 0.368 | 0.029 |
| | IGHG1 | 3.705 | 0.202 | 0.017 |
| | IGLL5 | 2.783 | 0.144 | 0.002 |
| | IGHG3 | 2.758 | 0.179 | 0.009 |
| | MZB1 | 2.173 | 0.555 | 0.013 |
| | IGHD | 2.088 | 0.154 | 0.004 |
| | SSR4 | 1.856 | 0.755 | 0.639 |
| | IGHG4 | 1.835 | 0.106 | 0.005 |
| | CD79A | 1.644 | 0.621 | 0.005 |
| | CD37 | 1.453 | 0.418 | 0.083 |
| | DERL3 | 1.452 | 0.473 | 0.008 |
| | HERPUD1 | 1.348 | 0.778 | 0.527 |
| | SEC11C | 1.295 | 0.561 | 0.202 |
| | CYBA | 1.264 | 0.838 | 0.437 |

| Cell Type | Gene | Average LogFC | pct1 | pct2 |
|---|---|---|---|---|
| Fibroblasts | DCN | 3.569 | 0.963 | 0.262 |
| | CFD | 3.281 | 0.792 | 0.275 |
| | APOD | 2.928 | 0.929 | 0.307 |
| | LUM | 2.787 | 0.879 | 0.090 |
| | TNFAIP6 | 2.634 | 0.740 | 0.111 |
| | COL1A2 | 2.604 | 0.785 | 0.094 |
| | COL1A1 | 2.458 | 0.643 | 0.074 |
| | GSN | 2.399 | 0.911 | 0.474 |
| | MMP3 | 2.392 | 0.204 | 0.078 |
| | COL3A1 | 2.331 | 0.584 | 0.069 |
| | MEG3 | 2.259 | 0.831 | 0.087 |
| | CCDC80 | 2.254 | 0.778 | 0.098 |
| | FBLN1 | 2.136 | 0.782 | 0.062 |
| | IGFBP6 | 2.101 | 0.675 | 0.085 |
| | SFRP2 | 2.043 | 0.639 | 0.036 |
| | COL6A2 | 1.987 | 0.906 | 0.199 |
| | MMP2 | 1.912 | 0.797 | 0.071 |
| | C1R | 1.882 | 0.831 | 0.138 |
| | C1S | 1.878 | 0.847 | 0.102 |
| | SERPINF1 | 1.849 | 0.777 | 0.084 |
| Lymphatic | CCL21 | 4.336 | 0.893 | 0.020 |
| | TFF3 | 3.067 | 0.864 | 0.070 |
| | MMRN1 | 2.840 | 0.860 | 0.009 |
| | CAVIN2 | 2.261 | 0.705 | 0.061 |
| | CLDN5 | 2.080 | 0.830 | 0.101 |
| | LYVE1 | 1.974 | 0.638 | 0.027 |
| | TFPI | 1.954 | 0.937 | 0.418 |
| | PPFIBP1 | 1.880 | 0.832 | 0.246 |
| | ANGPT2 | 1.784 | 0.592 | 0.125 |
| | ECSCR | 1.782 | 0.700 | 0.062 |
| | GNG11 | 1.776 | 0.850 | 0.278 |
| | CD9 | 1.744 | 0.889 | 0.438 |
| | FABP5 | 1.612 | 0.763 | 0.202 |
| | PROX1 | 1.597 | 0.564 | 0.033 |
| | FABP4 | 1.571 | 0.793 | 0.190 |
| | CAV1 | 1.444 | 0.863 | 0.411 |
| | IGFBP7 | 1.393 | 0.952 | 0.477 |
| | AKAP12 | 1.379 | 0.776 | 0.327 |
| | RGS16 | 1.359 | 0.557 | 0.157 |
| | RAMP2 | 1.356 | 0.600 | 0.150 |

| Cell Type | Gene | Average LogFC | pct1 | pct2 |
|---|---|---|---|---|
| T cells | IL7R | 3.085 | 0.732 | 0.024 |
| | CCL5 | 2.727 | 0.607 | 0.018 |
| | PTPRC | 2.538 | 0.882 | 0.058 |
| | CXCR4 | 2.471 | 0.875 | 0.100 |
| | GNLY | 2.400 | 0.186 | 0.008 |
| | NKG7 | 2.021 | 0.305 | 0.007 |
| | CD2 | 1.998 | 0.671 | 0.005 |
| | KLRB1 | 1.988 | 0.442 | 0.004 |
| | CD69 | 1.905 | 0.575 | 0.015 |
| | CCL4 | 1.827 | 0.328 | 0.052 |
| | CD52 | 1.807 | 0.564 | 0.024 |
| | CD3D | 1.793 | 0.604 | 0.004 |
| | CD7 | 1.750 | 0.525 | 0.013 |
| | SRGN | 1.722 | 0.904 | 0.271 |
| | TRBC2 | 1.716 | 0.539 | 0.006 |
| | ARHGDIB | 1.655 | 0.738 | 0.173 |
| | STK4 | 1.654 | 0.732 | 0.189 |
| | SARAF | 1.608 | 0.849 | 0.562 |
| | BTG1 | 1.607 | 0.972 | 0.761 |
| | TRBC1 | 1.603 | 0.426 | 0.004 |
| Myeloid | HLA-DRA | 3.412 | 0.921 | 0.177 |
| | HLA-DPA1 | 2.955 | 0.840 | 0.130 |
| | HLA-DPB1 | 2.840 | 0.820 | 0.184 |
| | RNASE1 | 2.828 | 0.525 | 0.064 |
| | CD74 | 2.788 | 0.934 | 0.299 |
| | HLA-DRB1 | 2.781 | 0.864 | 0.164 |
| | C1QA | 2.703 | 0.726 | 0.019 |
| | C1QB | 2.691 | 0.727 | 0.016 |
| | IL1B | 2.631 | 0.411 | 0.019 |
| | CCL3 | 2.607 | 0.477 | 0.040 |
| | TYROBP | 2.481 | 0.892 | 0.026 |
| | FCER1G | 2.458 | 0.862 | 0.033 |
| | HLA-DQA1 | 2.359 | 0.656 | 0.043 |
| | C1QC | 2.316 | 0.689 | 0.009 |
| | LYZ | 2.307 | 0.594 | 0.014 |
| | CD163 | 2.298 | 0.626 | 0.014 |
| | HLA-DQB1 | 2.168 | 0.687 | 0.084 |
| | CTSB | 2.167 | 0.837 | 0.337 |
| | AIF1 | 2.158 | 0.758 | 0.013 |
| | MS4A7 | 2.055 | 0.753 | 0.021 |

**Supplementary Table 3: Top Marker Genes Expressed in Major Breast Cell Types**
This table lists the top 20 marker genes expressed in the scRNA-seq data for each major cell type cluster based on average log fold-change and specificity of the gene expressed in the indicated cell type (pct1) compared to the other cell types (pct2). The columns listed (from left to right) indicate the name of the major cell type, the gene names, the average log-fold change (Average LogFC), the pct1 value indicating the fraction of cells within the cluster expressing the gene, the pct2 value indicating the fraction of cells in other clusters expressing the gene.

**Supplementary Table 4: Top Marker Genes Expressed in Major Breast Cell Types from Nuclei data**

| Nuclei Type | Gene | Average LogFC | pct1 | pct2 |
|---|---|---|---|---|
| Basal | KRT14 | 3.834 | 0.936 | 0.120 |
| | KRT17 | 3.568 | 0.938 | 0.156 |
| | SAA1 | 2.778 | 0.807 | 0.190 |
| | KRT5 | 2.527 | 0.876 | 0.057 |
| | DST | 2.446 | 0.895 | 0.403 |
| | ACTA2 | 2.235 | 0.783 | 0.134 |
| | TAGLN | 2.196 | 0.895 | 0.265 |
| | SFN | 2.061 | 0.803 | 0.111 |
| | MT1X | 1.996 | 0.834 | 0.464 |
| | ACTG2 | 1.963 | 0.628 | 0.022 |
| | MYLK | 1.890 | 0.709 | 0.093 |
| | TPM2 | 1.748 | 0.917 | 0.329 |
| | KRT6B | 1.611 | 0.490 | 0.057 |
| | FBXO32 | 1.543 | 0.623 | 0.131 |
| | S100A2 | 1.506 | 0.354 | 0.044 |
| | MT2A | 1.488 | 0.972 | 0.814 |
| | CNN1 | 1.425 | 0.569 | 0.060 |
| | TPM1 | 1.409 | 0.868 | 0.434 |
| | FBXO2 | 1.291 | 0.508 | 0.061 |
| | CRYAB | 1.290 | 0.758 | 0.250 |
| LumHR | AREG | 3.000 | 0.783 | 0.206 |
| | MUCL1 | 2.801 | 0.371 | 0.118 |
| | AZGP1 | 2.702 | 0.927 | 0.174 |
| | ANKRD30A | 2.471 | 0.765 | 0.023 |
| | KRT18 | 2.319 | 0.941 | 0.218 |
| | PIP | 2.274 | 0.354 | 0.069 |
| | KRT8 | 2.000 | 0.916 | 0.218 |
| | KRT19 | 1.904 | 0.866 | 0.191 |
| | S100A14 | 1.840 | 0.842 | 0.125 |
| | AGR2 | 1.759 | 0.588 | 0.020 |
| | TFF1 | 1.741 | 0.314 | 0.018 |
| | TCIM | 1.716 | 0.740 | 0.129 |
| | CD24 | 1.659 | 0.873 | 0.191 |
| | STC2 | 1.554 | 0.677 | 0.071 |
| | SYTL2 | 1.525 | 0.678 | 0.060 |
| | SPINT2 | 1.486 | 0.906 | 0.215 |
| | ELF3 | 1.454 | 0.803 | 0.123 |
| | CLDN4 | 1.386 | 0.841 | 0.169 |
| | AGR3 | 1.381 | 0.481 | 0.011 |
| | CLDN7 | 1.304 | 0.804 | 0.103 |
| LumSec | WFDC2 | 2.516 | 0.576 | 0.189 |
| | SCGB2A2 | 2.425 | 0.395 | 0.106 |
| | PI3 | 2.202 | 0.288 | 0.030 |
| | FDCSP | 2.199 | 0.174 | 0.046 |
| | SLPI | 2.195 | 0.719 | 0.098 |
| | LTF | 2.176 | 0.559 | 0.077 |
| | KRT15 | 2.146 | 0.715 | 0.101 |
| | S100A9 | 1.907 | 0.477 | 0.069 |
| | MMP7 | 1.783 | 0.496 | 0.053 |
| | CLDN4 | 1.733 | 0.802 | 0.186 |
| | SCGB3A1 | 1.679 | 0.351 | 0.094 |
| | KRT19 | 1.644 | 0.822 | 0.208 |
| | KRT7 | 1.582 | 0.870 | 0.277 |
| | KRT23 | 1.564 | 0.561 | 0.057 |
| | CD24 | 1.509 | 0.851 | 0.207 |
| | S100A8 | 1.493 | 0.191 | 0.032 |
| | TACSTD2 | 1.490 | 0.832 | 0.305 |
| | CCL28 | 1.470 | 0.643 | 0.042 |
| | PIGR | 1.452 | 0.555 | 0.035 |
| | RARRES1 | 1.405 | 0.432 | 0.118 |
| Vascular | SELE | 2.432 | 0.341 | 0.020 |
| | FABP4 | 2.421 | 0.631 | 0.150 |
| | ACKR1 | 2.361 | 0.435 | 0.018 |
| | CSF3 | 2.139 | 0.405 | 0.059 |
| | CLDN5 | 2.096 | 0.689 | 0.047 |
| | STC1 | 2.065 | 0.476 | 0.054 |
| | ANGPT2 | 1.946 | 0.629 | 0.075 |
| | TM4SF1 | 1.946 | 0.930 | 0.456 |
| | SPARCL1 | 1.853 | 0.823 | 0.366 |
| | ADAMTS9 | 1.812 | 0.665 | 0.079 |
| | IFI27 | 1.781 | 0.853 | 0.296 |
| | SERPINE1 | 1.751 | 0.584 | 0.274 |
| | ADGRL4 | 1.729 | 0.719 | 0.014 |
| | C2CD4B | 1.712 | 0.436 | 0.033 |
| | RBP7 | 1.709 | 0.499 | 0.060 |
| | GNG11 | 1.653 | 0.807 | 0.227 |
| | PECAM1 | 1.615 | 0.664 | 0.043 |
| | CD93 | 1.585 | 0.673 | 0.043 |
| | PGF | 1.576 | 0.337 | 0.085 |
| | AQP1 | 1.560 | 0.383 | 0.039 |
| Pericytes | C11orf96 | 2.633 | 0.896 | 0.321 |
| | RGS5 | 2.159 | 0.494 | 0.014 |
| | MT1A | 2.059 | 0.754 | 0.270 |
| | CCL20 | 1.588 | 0.143 | 0.046 |
| | PDK4 | 1.563 | 0.590 | 0.239 |
| | ADAMTS4 | 1.499 | 0.653 | 0.206 |
| | STEAP4 | 1.449 | 0.355 | 0.069 |
| | MYL9 | 1.442 | 0.584 | 0.307 |
| | EDNRB | 1.434 | 0.576 | 0.203 |
| | ADAMTS1 | 1.414 | 0.690 | 0.262 |
| | IGFBP7 | 1.408 | 0.940 | 0.460 |
| | TAGLN | 1.352 | 0.683 | 0.308 |
| | IL6 | 1.342 | 0.448 | 0.187 |
| | IGFBP5 | 1.313 | 0.534 | 0.210 |
| | MCAM | 1.300 | 0.551 | 0.092 |
| | ADIRF | 1.278 | 0.751 | 0.461 |
| | NR2F2 | 1.254 | 0.578 | 0.216 |
| | MYH11 | 1.252 | 0.311 | 0.050 |
| | GJA4 | 1.231 | 0.402 | 0.008 |
| | CRYAB | 1.226 | 0.561 | 0.286 |
| B cells | IGKC | 5.473 | 0.846 | 0.354 |
| | IGLC2 | 5.200 | 0.486 | 0.118 |
| | IGHA1 | 5.151 | 0.737 | 0.272 |
| | IGLC3 | 4.987 | 0.403 | 0.088 |
| | JCHAIN | 4.889 | 0.643 | 0.090 |
| | IGHA2 | 4.357 | 0.479 | 0.078 |
| | IGHM | 4.093 | 0.368 | 0.029 |
| | IGHG1 | 3.705 | 0.202 | 0.017 |
| | IGLL5 | 2.783 | 0.144 | 0.002 |
| | IGHG3 | 2.758 | 0.179 | 0.009 |
| | MZB1 | 2.173 | 0.555 | 0.013 |
| | IGHD | 2.088 | 0.154 | 0.004 |
| | SSR4 | 1.856 | 0.755 | 0.639 |
| | IGHG4 | 1.835 | 0.106 | 0.005 |
| | CD79A | 1.644 | 0.621 | 0.005 |
| | CD37 | 1.453 | 0.418 | 0.083 |
| | DERL3 | 1.452 | 0.473 | 0.008 |
| | HERPUD1 | 1.348 | 0.778 | 0.527 |
| | SEC11C | 1.295 | 0.561 | 0.202 |
| | CYBA | 1.264 | 0.838 | 0.437 |

| Nuclei Type | Gene | Average LogFC | pct1 | pct2 |
|---|---|---|---|---|
| Fibroblasts | DCN | 3.569 | 0.963 | 0.262 |
| | CFD | 3.281 | 0.792 | 0.275 |
| | APOD | 2.928 | 0.929 | 0.307 |
| | LUM | 2.787 | 0.879 | 0.090 |
| | TNFAIP6 | 2.634 | 0.740 | 0.111 |
| | COL1A2 | 2.604 | 0.785 | 0.094 |
| | COL1A1 | 2.458 | 0.643 | 0.074 |
| | GSN | 2.399 | 0.911 | 0.474 |
| | MMP3 | 2.392 | 0.204 | 0.078 |
| | COL3A1 | 2.331 | 0.584 | 0.069 |
| | MEG3 | 2.259 | 0.831 | 0.087 |
| | CCDC80 | 2.254 | 0.778 | 0.098 |
| | FBLN1 | 2.136 | 0.782 | 0.062 |
| | IGFBP6 | 2.101 | 0.675 | 0.085 |
| | SFRP2 | 2.043 | 0.639 | 0.036 |
| | COL6A2 | 1.987 | 0.906 | 0.199 |
| | MMP2 | 1.912 | 0.797 | 0.071 |
| | C1R | 1.882 | 0.831 | 0.138 |
| | C1S | 1.878 | 0.847 | 0.102 |
| | SERPINF1 | 1.849 | 0.777 | 0.084 |
| Lymphatic | CCL21 | 4.336 | 0.893 | 0.020 |
| | TFF3 | 3.067 | 0.864 | 0.070 |
| | MMRN1 | 2.840 | 0.860 | 0.009 |
| | CAVIN2 | 2.261 | 0.705 | 0.061 |
| | CLDN5 | 2.080 | 0.830 | 0.101 |
| | LYVE1 | 1.974 | 0.638 | 0.027 |
| | TFPI | 1.954 | 0.937 | 0.418 |
| | PPFIBP1 | 1.880 | 0.832 | 0.246 |
| | ANGPT2 | 1.784 | 0.592 | 0.125 |
| | ECSCR | 1.782 | 0.700 | 0.062 |
| | GNG11 | 1.776 | 0.850 | 0.278 |
| | CD9 | 1.744 | 0.889 | 0.438 |
| | FABP5 | 1.612 | 0.763 | 0.202 |
| | PROX1 | 1.597 | 0.564 | 0.033 |
| | FABP4 | 1.571 | 0.793 | 0.190 |
| | CAV1 | 1.444 | 0.863 | 0.411 |
| | IGFBP7 | 1.393 | 0.952 | 0.477 |
| | AKAP12 | 1.379 | 0.776 | 0.327 |
| | RGS16 | 1.359 | 0.557 | 0.157 |
| | RAMP2 | 1.356 | 0.600 | 0.150 |

| Nuclei Type | Gene | Average LogFC | pct1 | pct2 |
|---|---|---|---|---|
| T cells | IL7R | 3.085 | 0.732 | 0.024 |
| | CCL5 | 2.727 | 0.607 | 0.018 |
| | PTPRC | 2.538 | 0.882 | 0.058 |
| | CXCR4 | 2.471 | 0.875 | 0.100 |
| | GNLY | 2.400 | 0.186 | 0.008 |
| | NKG7 | 2.021 | 0.305 | 0.007 |
| | CD2 | 1.998 | 0.671 | 0.005 |
| | KLRB1 | 1.988 | 0.442 | 0.004 |
| | CD69 | 1.905 | 0.575 | 0.015 |
| | CCL4 | 1.827 | 0.328 | 0.052 |
| | CD52 | 1.807 | 0.564 | 0.024 |
| | CD3D | 1.793 | 0.604 | 0.004 |
| | CD7 | 1.750 | 0.525 | 0.013 |
| | SRGN | 1.722 | 0.904 | 0.271 |
| | TRBC2 | 1.716 | 0.539 | 0.006 |
| | ARHGDIB | 1.655 | 0.738 | 0.173 |
| | STK4 | 1.654 | 0.732 | 0.189 |
| | SARAF | 1.608 | 0.849 | 0.562 |
| | BTG1 | 1.607 | 0.972 | 0.761 |
| | TRBC1 | 1.603 | 0.426 | 0.004 |
| Myeloid | HLA-DRA | 3.412 | 0.921 | 0.177 |
| | HLA-DPA1 | 2.955 | 0.840 | 0.130 |
| | HLA-DPB1 | 2.840 | 0.820 | 0.184 |
| | RNASE1 | 2.828 | 0.525 | 0.064 |
| | CD74 | 2.788 | 0.934 | 0.299 |
| | HLA-DRB1 | 2.781 | 0.864 | 0.164 |
| | C1QA | 2.703 | 0.726 | 0.019 |
| | C1QB | 2.691 | 0.727 | 0.016 |
| | IL1B | 2.631 | 0.411 | 0.019 |
| | CCL3 | 2.607 | 0.477 | 0.040 |
| | TYROBP | 2.481 | 0.892 | 0.026 |
| | FCER1G | 2.458 | 0.862 | 0.033 |
| | HLA-DQA1 | 2.359 | 0.656 | 0.043 |
| | C1QC | 2.316 | 0.689 | 0.009 |
| | LYZ | 2.307 | 0.594 | 0.014 |
| | CD163 | 2.298 | 0.626 | 0.014 |
| | HLA-DQB1 | 2.168 | 0.687 | 0.084 |
| | CTSB | 2.167 | 0.837 | 0.337 |
| | AIF1 | 2.158 | 0.758 | 0.013 |
| | MS4A7 | 2.055 | 0.753 | 0.021 |

**Supplementary Table 4: Top Marker Genes Expressed in Major Breast Cell Types from Nuclei data**

This table lists the top 20 marker genes expressed in the scRNA-seq data for each major cell type cluster based on average log fold-change and specificity of the gene expressed in the indicated cell type (pct1) compared to the other cell types (pc2). The columns listed (from left to right) indicate the name of the major cell type, the gene names, the average log-fold change (Average LogFC), the pct1 value indicating the fraction of cells within the cluster expressing the gene, the pct2 value indicating the fraction of cells in other clusters expressing the gene.

Supplementary Table 5: Quality Control Metrics for Spatial Transcriptomics Data

| Patient ID | Number of spots under tissue | Number of reads | Median UMI per spot | Median Genes per spot | Total genes detected | Reads mapped to exonic regions | Reads mapped to intergenic regions | Reads mapped to intronic regions | Reads mapped to transcriptome |
|---|---|---|---|---|---|---|---|---|---|
| P10 | 2741 | 461115960 | 3031 | 1546 | 23050 | 0.836 | 0.022 | 0.032 | 0.816 |
| P35 | 1896 | 478182981 | 2223 | 1184 | 22295 | 0.878 | 0.019 | 0.026 | 0.855 |
| P46 | 3129 | 404340693 | 2348 | 1315 | 22442 | 0.823 | 0.026 | 0.052 | 0.803 |
| P47 | 3449 | 362966308 | 1108 | 797 | 22121 | 0.837 | 0.027 | 0.06 | 0.817 |

**Supplementary Table 5: Quality Control Metrics for Spatial Transcriptomics Data**

This table lists the quality control metrics for breast tissue samples sequenced with the Spatial Transcriptomics platform (Visium, 10X Genomics). The columns listed (from left to right) indicate the Patient ID, Number of spots under tissue, Number of reads, Median UMI per spot, Median Genes per spot, Total genes detected, Reads mapping rate (exonic, intergenic, intronic regions and transcriptome).

**Supplementary Table 6: Custom Targeted Gene Panel for smFISH**

| Cell Type | Gene | Cell Annotation (module score) |
|---|---|---|
| Basal | KRT5 | 1 |
| | ACTG2 | 1 |
| | TUBB2B | 0 |
| | COL17A1 | 1 |
| | KRT6B | 0 |
| | LAMA3 | 1 |
| LumHR | AR | 1 |
| | ESR1 | 1 |
| | PGR | 1 |
| | AREG | 1 |
| | OXTR | 0 |
| | ANKRD30A | 1 |
| | AGR3 | 1 |
| | AGR2 | 1 |
| | TMC5 | 1 |
| | DNAJC12 | 1 |
| | RASEF | 1 |
| LumSec | SLPI | 1 |
| | LTF | 1 |
| | KRT15 | 1 |
| | MMP7 | 1 |
| | CCL28 | 1 |
| | ALDH1A3 | 1 |
| | PIGR | 1 |
| Fibroblasts | LUM | 1 |
| | TNFAIP6 | 0 |
| | COL1A2 | 1 |
| | COL1A1 | 1 |
| | FBLN1 | 1 |
| | MMP2 | 1 |
| | SERPINF1 | 1 |
| Pericytes | MCAM | 0 |
| | RGS5 | 1 |
| | GJA4 | 1 |
| | NDUFA4L2 | 1 |
| | SSTR2 | 1 |
| | AVPR1A | 0 |
| | PLN | 1 |
| | EDNRA | 1 |
| Lymphatic | CCL21 | 1 |
| | MMRN1 | 0 |
| | PROX1 | 1 |
| | SCN3B | 1 |
| | PKHD1L1 | 0 |
| | TBX1 | 1 |
| | PGM5 | 1 |
| Vascular | VWF | 0 |
| | SELE | 0 |
| | ACKR1 | 1 |
| | CSF3 | 0 |
| | ADGRL4 | 0 |
| | RBP7 | 1 |
| | PGF | 1 |
| | AQP1 | 1 |

| Cell Type | Gene | Cell Annotation (module score) |
|---|---|---|
| Myeloid | CD14 | 1 |
| | CD68 | 1 |
| | C1QA | 1 |
| | C1QB | 1 |
| | FCER1G | 0 |
| | C1QC | 1 |
| | LYZ | 1 |
| | CD163 | 1 |
| | MSR1 | 1 |
| Mast | IL18R1 | 0 |
| | HPGD | 0 |
| | HDC | 0 |
| | SLC18A2 | 0 |
| | CPA3 | 0 |
| | CD69 | 0 |
| | HPGDS | 0 |
| T cells | CD3D | 0 |
| | CD3E | 0 |
| | CD4 | 1 |
| | CD8A | 1 |
| | IL7R | 1 |
| | CCL5 | 1 |
| | NKG7 | 1 |
| | GZMB | 1 |
| | CD2 | 1 |
| B cells | CD27 | 0 |
| | DERL3 | 0 |
| | JCHAIN | 1 |
| | IGHM | 1 |
| | TNFRSF17 | 0 |
| | MZB1 | 1 |
| | CD79A | 0 |
| Adipocytes | PLIN1 | 0 |
| | ADIPOQ | 0 |
| | PLIN4 | 0 |
| | GPD1 | 0 |
| | LPL | 0 |
| | LIPE | 0 |
| Epithelial | EPCAM | 0 |
| Endothelial | PECAM1 | 0 |
| Proliferation | MKI67 | 0 |
| Immune | PTPRC | 0 |
| Immune | TRAC | 0 |
| Mis | TP63 | 0 |
| Luminal | KRT19 | 0 |
| B cells | IGHA2 | Probe Failure |

**Supplementary Table 6: Custom Targeted Gene Panel for smFISH**

This table lists the top expressed genes selected for each breast cell type from the scRNA-seq data to generate a custom 100-gene panel for smFISH analysis (Resolve Biosciences). The entire panel of 100 genes was used to profile each of the breast tissue samples, to determine the spatial distribution of the breast cell types in the tissue sections. The table shows the cell type, the genes used for classification and cell annotation module score (if the gene was used for annotating the cell, 1 = yes, 0 = no, not included).

#### Supplementary Table 7: Quality Control Metrics for smFISH Data

| Section ID | Patient ID | Total Counts (raw) | Total Counts (post filter) | Total Cells (raw) | Filtered Cells (post filter) |
|---|---|---|---|---|---|
| P46-S1 | P46 | 1491813 | 1301494 | 11929 | 9879 |
| P35-S1 | P35 | 913325 | 702059 | 7604 | 6683 |
| P47-S1 | P47 | 667314 | 615735 | 6872 | 6050 |
| P64-S1 | P64 | 491862 | 330047 | 4276 | 3390 |
| P64-S2 | P64 | 580993 | 157526 | 4754 | 4018 |
| P46-S2 | P46 | 460881 | 319725 | 5847 | 3961 |
| P63-S1 | P63 | 113031 | 75978 | 1244 | 868 |
| P64-S3 | P64 | 1549406 | 853581 | 5441 | 4591 |
| P46-S3 | P46 | 1991213 | 1018173 | 11788 | 8989 |
| P46-S4 | P46 | 703202 | 343803 | 3635 | 2906 |
| P63-S2 | P63 | 208133 | 95192 | 1290 | 928 |
| P63-S3 | P63 | 1019583 | 545389 | 4597 | 3372 |

**Supplementary Table 7: Quality Control Metrics for smFISH Data**

The table lists the quality control metrics for the 12 breast tissue samples profiled with the custom 100-gene panel for smFISH analysis (Resolve Biosciences). The columns listed (from left to right) indicate the Section ID, Patient ID, the number of transcripts captured under tissue, the number of transcripts after filtering, the number of cells detected and the number of cells after filtering.

#### Supplementary Table 8: Samples for CODEX

| Sample | Patient ID | Age | Ethnicity | BMI | Prior Cancer History | Cells profiled |
|---|---|---|---|---|---|---|
| p65_s1 | P65 | 24 | W | 32.4 | endocrine, thyroid | 78219 |
| p66_s1 | P66 | 19 | W | 30.6 | NA | 62302 |
| p67_s1 | P67 | 29 | B | 40.1 | NA | 40781 |
| p68_s1 | P68 | 24 | W | 30.5 | NA | 49985 |

**Supplementary Table 8: Samples for CODEX**

This table lists the 4 breast tissue samples that were used for CODEX analysis, along with the clinical metadata for each woman, and total number of cells that were profiled and analyzed.

#### Supplementary Table 9: CODEX 34-Antibody Panel Design

| Cycle # | Antibody | Clone | | Reporter | Ratio | Ab vendor |
|---|---|---|---|---|---|---|
| Cycle1 | blank | | | | | |
| | blank | | | | | |
| | blank | | | | | |
| Cycle2 | Keratin19 | A53-B/A2 | BX025 | RX025-A750 | 1:50 | Biolegend |
| | CD8 | C8/144B | BX026 | BX026-ATTO550 | 1:200 | Akoya |
| | PCNA | PC10 | BX020 | RX020-Cy5 | 1:50 | Biolegend |
| Cycle3 | Vimentin | RV202 | BX034 | RX034-A750 | 1:50 | BD |
| | CD31 | EP3095 | BX001 | RX001-ATTO550 | 1:200 | Akoya |
| | CD3e | EP449E | BX045 | RX045-Cy5 | 1:150 | Akoya |
| Cycle4 | Keratin7 | W16155A | BX019 | RX019-A750 | 1:50 | Biolegend |
| | Ki67 | B56 | BX047 | RX047-ATTO550 | 1:200 | Akoya |
| | CD4 | EPR6855 | BX003 | RX003-Cy5 | 1:200 | Akoya |
| Cycle5 | CD227 | HMPV | BX004 | BX004-A750 | 1:50 | BD |
| | Perlecan | 5D7-2E4 | BX017 | RX017-ATTO550 | 1:50 | BD |
| | CD45 | 2D1 | BX021 | RX021-Cy5 | 1:100 | Akoya |
| Cycle6 | Keratin17 | W16131A | BX022 | RX022-A750 | 1:50 | Biolegend |
| | Podoplanin | NC-08 | BX023 | BX023-ATTO550 | 1:200 | Akoya |
| | CD68 | KP1 | BX015 | RX015-Cy5 | 1:200 | Akoya |
| Cycle7 | Empty | | | | | |
| | CD14 | EPR3653 | BX037 | RX037-ATTO550 | 1:50 | Abcam |
| | CollagenIV | EPR20966 | BX042 | RX042-Cy5 | 1:50 | Abcam |
| Cycle8 | Keratin18 | DA-7 | BX049 | RX049-A750 | 1:50 | Biolegend |
| | HLA-DPB1 | EPR11226 | BX035 | RX035-ATTO550 | 1:50 | Abcam |
| | CD11c | 118/A5 | BX024 | RX024-Cy5 | 1:200 | Akoya |
| Cycle9 | Empty | | | | | |
| | E-Cadherin | 4A2C7 | BX014 | RX014-ATTO550 | 1:200 | Akoya |
| | PR | KMC912 | BX007 | RX007-Cy5 | 1:50 | ThermoFisher |
| Cycle10 | Keratin8 | 1E8 | BX040 | RX040-A750 | 1:50 | Biolegend |
| | Keratin14 | Poly19053 | BX032 | BX032-ATTO550 | 1:300 | Akoya |
| | TP63 | W15093A | BX033 | RX033-Cy5 | 1:50 | Biolegend |
| Cycle11 | Empty | | | | | |
| | Empty | | | | | |
| | SMA | 1A4 | BX030 | RX030-Cy5 | 1:50 | Abcam |
| Cycle12 | Empty | | | | | |
| | Empty | | | | | |
| | Runx3 | R3-5G4 | BX036 | RX036-Cy5 | 1:50 | Biolegend |
| Cycle13 | Empty | | | | | |
| | Empty | | | | | |
| | CD66e | BSB-13 | BX016 | RX016-Cy5 | 1:50 | Biolegend |
| | Empty | | | | | |

| | | | | | | |
|---|---|---|---|---|---|---|
| Cycle14 | Empty | | | | | |
| | BCL6 | K112-91 | BX041 | RX041-Cy5 | 1:50 | BD |
| Cycle15 | Empty | | | | | |
| | Empty | | | | | |
| | Foxp3 | 259D/C7 | BX027 | RX027-Cy5 | 1:40 | BD |
| Cycle16 | Empty | | | | | |
| | Empty | | | | | |
| | LIF | M1506B09 | BX006 | RX006-Cy5 | 1:200 | Akoya |
| Cycle17 | Empty | | | | | |
| | Empty | | | | | |
| | GranzymeB | D6E9W | BX046 | RX046-Cy5 | 1:50 | CST |
| Cycle18 | Empty | | | | | |
| | Empty | | | | | |
| | BCL2 | N46-467 | BX029 | RX029-Cy5 | 1:50 | BD |
| Cycle19 | Empty | | | | | |
| | Empty | | | | | |
| | Keratin5 | Poly19055 | BX005 | RX005-Cy5 | 1:50 | Biolegend |
| Cycle20 | blank | | | | | |
| | blank | | | | | |
| | blank | | | | | |

**Supplementary Table 9: CODEX 34-Antibody Panel Design**

This table lists the cycle order and description of the 34 antibodies run in the CODEX pane. The columns listed (from left to right) indicate, the cycle, the antibody, the clone, the reporter, the dilution ratio use and antibody vendor.

Supplementary Table 10: Top Gene Markers Expressed for Breast Cell States

| Cell state | Gene | Average LogFC | pct1 | pct2 | Adjusted P-value |
|---|---|---|---|---|---|
| **Epithelial** | | | | | |
| Basal | KRT14 | 3.441 | 0.962 | 0.256 | 0 |
| | TAGLN | 3.065 | 0.917 | 0.147 | 0 |
| | MT2A | 2.856 | 0.983 | 0.831 | 0 |
| | ACTA2 | 2.752 | 0.819 | 0.067 | 0 |
| | KRT17 | 2.665 | 0.967 | 0.388 | 0 |
| | MT1X | 2.629 | 0.879 | 0.474 | 0 |
| | TPM2 | 2.593 | 0.945 | 0.109 | 0 |
| | DST | 2.357 | 0.927 | 0.579 | 0 |
| | KRT5 | 2.341 | 0.917 | 0.163 | 0 |
| | MYLK | 2.125 | 0.758 | 0.032 | 0 |
| | ACTG2 | 2.007 | 0.665 | 0.022 | 0 |
| | CALD1 | 1.947 | 0.945 | 0.351 | 0 |
| | MT1E | 1.801 | 0.714 | 0.403 | 0 |
| | CAV1 | 1.701 | 0.757 | 0.135 | 0 |
| | NNMT | 1.696 | 0.911 | 0.391 | 0 |
| | SAA1 | 1.660 | 0.844 | 0.417 | 0 |
| | APOE | 1.600 | 0.578 | 0.125 | 0 |
| | CXCL14 | 1.596 | 0.375 | 0.043 | 0 |
| | THBS1 | 1.582 | 0.594 | 0.234 | 0 |
| | CNN1 | 1.559 | 0.613 | 0.023 | 0 |
| LumHR-major | AREG | 2.034 | 0.799 | 0.393 | 0 |
| | TCIM | 1.550 | 0.792 | 0.122 | 0 |
| | ANKRD30A | 1.540 | 0.787 | 0.102 | 0 |
| | KRT18 | 1.419 | 0.982 | 0.561 | 0 |
| | AGR2 | 1.380 | 0.630 | 0.102 | 0 |
| | SYTL2 | 1.358 | 0.734 | 0.097 | 0 |
| | HSPB1 | 1.285 | 0.798 | 0.358 | 0 |
| | AZGP1 | 1.262 | 0.968 | 0.455 | 0 |
| | TM4SF1 | 1.207 | 0.965 | 0.789 | 0 |
| | AGR3 | 1.200 | 0.516 | 0.075 | 0 |
| | TFF1 | 1.199 | 0.315 | 0.065 | 0 |
| | AQP3 | 1.137 | 0.358 | 0.177 | 0 |
| | ADIRF | 1.038 | 0.907 | 0.385 | 0 |
| | TMC5 | 1.031 | 0.672 | 0.093 | 0 |
| | FAM107B | 1.030 | 0.675 | 0.256 | 0 |
| | STC1 | 1.003 | 0.241 | 0.053 | 0 |
| | KRT8 | 0.989 | 0.967 | 0.744 | 0 |
| | XBP1 | 0.987 | 0.944 | 0.689 | 0 |
| | DNAJC12 | 0.986 | 0.670 | 0.093 | 0 |
| | C15orf48 | 0.972 | 0.580 | 0.143 | 0 |
| LumHR-active | CXCL13 | 2.052 | 0.459 | 0.023 | 0 |
| | DIO2 | 1.776 | 0.397 | 0.048 | 0 |
| | EREG | 1.547 | 0.537 | 0.084 | 0 |
| | TFF3 | 1.518 | 0.772 | 0.172 | 0 |
| | PTHLH | 1.355 | 0.642 | 0.134 | 0 |
| | MYBPC1 | 1.209 | 0.677 | 0.061 | 0 |
| | TCIM | 1.195 | 0.917 | 0.317 | 0 |
| | CA2 | 1.191 | 0.556 | 0.144 | 0 |
| | ANKRD30A | 1.189 | 0.923 | 0.301 | 0 |
| | SPINK1 | 1.144 | 0.179 | 0.045 | 0 |
| | EFHD1 | 1.141 | 0.874 | 0.329 | 0 |
| | TFF1 | 1.140 | 0.528 | 0.131 | 0 |
| | ALB | 1.126 | 0.130 | 0.037 | 5.04E-231 |
| | HSPB1 | 1.015 | 0.849 | 0.488 | 0 |
| | AREG | 0.983 | 0.927 | 0.509 | 0 |
| | FASN | 0.944 | 0.683 | 0.235 | 0 |
| | AZGP1 | 0.943 | 0.988 | 0.608 | 0 |
| | SLC26A3 | 0.913 | 0.321 | 0.045 | 0 |
| | RAB11FIP1 | 0.901 | 0.872 | 0.403 | 0 |
| | XBP1 | 0.886 | 0.966 | 0.764 | 0 |
| LumHR-SCGB | PIP | 3.343 | 0.916 | 0.179 | 0 |
| | SCGB1D2 | 2.850 | 0.375 | 0.102 | 0 |
| | MUCL1 | 2.583 | 0.725 | 0.226 | 0 |
| | SCGB2A2 | 2.425 | 0.588 | 0.216 | 0 |
| | SERPINA1 | 2.146 | 0.800 | 0.131 | 0 |
| | SCGB3A1 | 2.067 | 0.642 | 0.187 | 0 |
| | TFPI2 | 1.697 | 0.607 | 0.063 | 0 |
| | CA2 | 1.530 | 0.433 | 0.153 | 0 |
| | AZGP1 | 1.276 | 0.994 | 0.613 | 0 |
| | HMGB3 | 1.212 | 0.420 | 0.112 | 0 |
| | CP | 1.209 | 0.326 | 0.059 | 0 |
| | SAT1 | 1.160 | 0.977 | 0.881 | 0 |
| | CST3 | 1.034 | 0.946 | 0.623 | 0 |
| | HSPB1 | 0.989 | 0.826 | 0.494 | 0 |
| | KRT23 | 0.967 | 0.639 | 0.273 | 0 |
| | TNFSF10 | 0.886 | 0.659 | 0.290 | 0 |
| | CYP4X1 | 0.885 | 0.471 | 0.040 | 0 |
| | TIMP1 | 0.874 | 0.846 | 0.543 | 0 |
| | STC2 | 0.868 | 0.848 | 0.417 | 0 |
| | CYB5A | 0.835 | 0.916 | 0.510 | 0 |

| Cell state | Gene | Average LogFC | pct1 | pct2 | Adjusted P-value |
|---|---|---|---|---|---|
| **Epithelial** | | | | | |
| LumSec-major | S100A9 | 2.202 | 0.621 | 0.091 | 0 |
| | PI3 | 2.138 | 0.494 | 0.073 | 0 |
| | WFDC2 | 2.119 | 0.866 | 0.637 | 0 |
| | SLPI | 2.103 | 0.921 | 0.191 | 0 |
| | FDCSP | 2.037 | 0.246 | 0.077 | 0 |
| | SCGB2A2 | 1.872 | 0.544 | 0.157 | 0 |
| | RARRES1 | 1.834 | 0.657 | 0.138 | 0 |
| | S100A8 | 1.831 | 0.296 | 0.036 | 0 |
| | LTF | 1.822 | 0.799 | 0.124 | 0 |
| | KRT15 | 1.726 | 0.780 | 0.195 | 0 |
| | MMP7 | 1.703 | 0.606 | 0.078 | 0 |
| | LCN2 | 1.552 | 0.616 | 0.058 | 0 |
| | ALDH1A3 | 1.499 | 0.784 | 0.105 | 0 |
| | PTN | 1.419 | 0.441 | 0.218 | 0 |
| | KRT23 | 1.336 | 0.727 | 0.186 | 0 |
| | CCL28 | 1.237 | 0.708 | 0.083 | 0 |
| | PIGR | 1.226 | 0.647 | 0.061 | 0 |
| | SLC25A37 | 1.166 | 0.864 | 0.331 | 0 |
| | AKR1C3 | 1.161 | 0.342 | 0.068 | 0 |
| | RGS2 | 1.091 | 0.408 | 0.073 | 0 |
| LumSec-KIT | PLCG2 | 5.212 | 0.901 | 0.174 | 0 |
| | SCGB3A1 | 1.801 | 0.492 | 0.194 | 0 |
| | MAFB | 1.593 | 0.485 | 0.156 | 0 |
| | SNORC | 1.519 | 0.554 | 0.133 | 0 |
| | ELF5 | 1.496 | 0.549 | 0.074 | 0 |
| | JUN | 1.453 | 0.911 | 0.831 | 0 |
| | KRT15 | 1.452 | 0.808 | 0.290 | 0 |
| | CCL28 | 1.357 | 0.666 | 0.186 | 0 |
| | MTRNR2L12 | 1.307 | 0.727 | 0.516 | 0 |
| | AL355075.4 | 1.295 | 0.331 | 0.061 | 0 |
| | GOLGA8A | 1.251 | 0.455 | 0.111 | 0 |
| | NR4A1 | 1.205 | 0.454 | 0.185 | 0 |
| | PIK3C2G | 1.166 | 0.449 | 0.108 | 0 |
| | FOSB | 1.158 | 0.758 | 0.514 | 0 |
| | GABRP | 1.103 | 0.464 | 0.131 | 0 |
| | PLA2R1 | 1.062 | 0.350 | 0.103 | 0 |
| | PIGR | 1.036 | 0.474 | 0.160 | 0 |
| | KIT | 1.027 | 0.418 | 0.063 | 0 |
| | FOS | 1.016 | 0.819 | 0.539 | 0 |
| | AC025164.1 | 0.960 | 0.237 | 0.023 | 0 |
| LumSec-HLA | CCL20 | 3.480 | 0.544 | 0.045 | 0 |
| | PI3 | 2.992 | 0.725 | 0.145 | 0 |
| | SLPI | 2.564 | 0.988 | 0.318 | 0 |
| | CD74 | 2.096 | 0.941 | 0.353 | 0 |
| | S100A9 | 1.484 | 0.786 | 0.183 | 0 |
| | CRABP2 | 1.441 | 0.972 | 0.427 | 0 |
| | HLA-DRA | 1.428 | 0.732 | 0.141 | 0 |
| | ANXA1 | 1.423 | 0.999 | 0.632 | 0 |
| | HLA-DRB1 | 1.371 | 0.739 | 0.121 | 0 |
| | HIST1H1C | 1.259 | 0.834 | 0.306 | 0 |
| | B2M | 1.224 | 1.000 | 0.977 | 0 |
| | CHI3L2 | 1.202 | 0.823 | 0.129 | 0 |
| | LTF | 1.180 | 0.866 | 0.242 | 0 |
| | IL32 | 1.158 | 0.796 | 0.275 | 0 |
| | TNFAIP2 | 1.074 | 0.682 | 0.178 | 0 |
| | S100A1 | 1.038 | 0.826 | 0.170 | 0 |
| | RBP1 | 1.024 | 0.924 | 0.355 | 0 |
| | HLA-A | 1.013 | 0.998 | 0.829 | 0 |
| | PIGR | 1.001 | 0.813 | 0.162 | 0 |
| | CXCL17 | 0.971 | 0.643 | 0.071 | 0 |
| LumSec-lac | LALBA | 5.326 | 0.524 | 0.009 | 0 |
| | CSN1S1 | 4.279 | 0.548 | 0.014 | 0 |
| | CSN2 | 4.093 | 0.405 | 0.005 | 0 |
| | CSN3 | 3.842 | 0.506 | 0.010 | 0 |
| | LYZ | 2.895 | 0.186 | 0.013 | 0 |
| | LTF | 2.310 | 0.574 | 0.242 | 0 |
| | IGKC | 2.280 | 0.554 | 0.268 | 4.37E-260 |
| | SPP1 | 2.006 | 0.307 | 0.106 | 9.82E-167 |
| | FDCSP | 1.943 | 0.322 | 0.105 | 4.28E-202 |
| | IGHA1 | 1.822 | 0.466 | 0.196 | 1.96E-236 |
| | KRT15 | 1.642 | 0.720 | 0.295 | 0 |
| | PIGR | 1.633 | 0.429 | 0.163 | 2.23E-236 |
| | S100A1 | 1.603 | 0.484 | 0.170 | 0 |
| | ZG16B | 1.377 | 0.283 | 0.265 | 5.17E-08 |
| | FABP3 | 1.338 | 0.250 | 0.009 | 0 |
| | PLIN2 | 1.290 | 0.338 | 0.182 | 9.47E-95 |
| | S100A9 | 1.289 | 0.406 | 0.184 | 4.49E-147 |
| | CLU | 1.260 | 0.542 | 0.674 | 5.48E-21 |
| | HLA-DRA | 1.174 | 0.336 | 0.143 | 6.67E-140 |
| | SCGB3A1 | 1.171 | 0.314 | 0.199 | 5.72E-48 |
| LumSec-prol | PCLAF | 1.443 | 0.554 | 0.011 | 0 |
| | UBE2C | 1.402 | 0.472 | 0.004 | 0 |
| | TYMS | 1.392 | 0.562 | 0.017 | 0 |
| | TOP2A | 1.387 | 0.441 | 0.007 | 0 |
| | NUSAP1 | 1.357 | 0.503 | 0.025 | 0 |
| | CRABP1 | 1.236 | 0.503 | 0.047 | 0 |
| | CDK1 | 1.150 | 0.416 | 0.003 | 0 |
| | TK1 | 1.042 | 0.424 | 0.026 | 0 |
| | BIRC5 | 1.011 | 0.370 | 0.005 | 0 |
| | CENPF | 0.987 | 0.259 | 0.020 | 0 |
| | MAD2L1 | 0.959 | 0.397 | 0.023 | 0 |
| | PBK | 0.865 | 0.312 | 0.001 | 0 |
| | TPX2 | 0.842 | 0.284 | 0.006 | 0 |
| | CCNB2 | 0.832 | 0.279 | 0.004 | 0 |
| | UBE2T | 0.832 | 0.337 | 0.026 | 0 |
| | MKI67 | 0.805 | 0.249 | 0.001 | 0 |
| | FAM111B | 0.726 | 0.220 | 0.009 | 0 |
| | ZWINT | 0.711 | 0.308 | 0.016 | 0 |
| | CDC20 | 0.698 | 0.216 | 0.009 | 0 |
| | CENPM | 0.690 | 0.284 | 0.015 | 0 |

**Supplementary Table 10: Top Gene Markers Expressed for Breast Cell States**

| Cell state | Gene | Average LogFC | pct1 | pct2 | Adjusted P-value |
|---|---|---|---|---|---|
| | | **Fibroblasts** | | | |
| | POSTN | 1.925 | 0.759 | 0.158 | 0 |
| | COL3A1 | 1.849 | 0.947 | 0.491 | 0 |
| | COL1A1 | 1.639 | 0.928 | 0.556 | 0 |
| | PENK | 1.509 | 0.227 | 0.038 | 0 |
| | IGF1 | 1.354 | 0.876 | 0.528 | 0 |
| | TNC | 1.185 | 0.487 | 0.074 | 0 |
| | ADAM12 | 1.103 | 0.725 | 0.183 | 0 |
| | IGFBP2 | 1.088 | 0.560 | 0.122 | 0 |
| | COL5A2 | 1.071 | 0.804 | 0.382 | 0 |
| Fibro-matrix | SPARC | 1.041 | 0.943 | 0.642 | 0 |
| | TAC1 | 1.009 | 0.624 | 0.187 | 0 |
| | COL1A2 | 0.998 | 0.970 | 0.737 | 0 |
| | DLK1 | 0.977 | 0.569 | 0.150 | 0 |
| | HGF | 0.952 | 0.649 | 0.266 | 0 |
| | VCAN | 0.889 | 0.870 | 0.572 | 0 |
| | HSPA6 | 0.815 | 0.145 | 0.058 | 0 |
| | MXRA5 | 0.807 | 0.657 | 0.288 | 0 |
| | ECEL1 | 0.804 | 0.312 | 0.006 | 0 |
| | FN1 | 0.785 | 0.748 | 0.510 | 0 |
| | COL15A1 | 0.785 | 0.698 | 0.375 | 0 |
| | FOS | 1.994 | 0.654 | 0.365 | 0 |
| | CFD | 1.824 | 0.996 | 0.723 | 0 |
| | GPX3 | 1.673 | 0.895 | 0.316 | 0 |
| | GSN | 1.517 | 0.984 | 0.896 | 0 |
| | WISP2 | 1.439 | 0.796 | 0.202 | 0 |
| | CXCL14 | 1.429 | 0.657 | 0.360 | 0 |
| | PLA2G2A | 1.426 | 0.373 | 0.030 | 0 |
| | ADH1B | 1.417 | 0.688 | 0.155 | 0 |
| | GPC3 | 1.386 | 0.943 | 0.560 | 0 |
| Fibro-prematrix | JUN | 1.360 | 0.840 | 0.623 | 0 |
| | MGST1 | 1.345 | 0.911 | 0.306 | 0 |
| | PCOLCE2 | 1.254 | 0.612 | 0.042 | 0 |
| | FBLN2 | 1.200 | 0.803 | 0.357 | 0 |
| | IGFBP3 | 1.180 | 0.595 | 0.233 | 0 |
| | S100A4 | 1.172 | 0.916 | 0.674 | 0 |
| | PDK4 | 1.137 | 0.592 | 0.176 | 0 |
| | MFAP5 | 1.072 | 0.618 | 0.209 | 0 |
| | SVEP1 | 1.057 | 0.665 | 0.231 | 0 |
| | MYOC | 1.047 | 0.324 | 0.015 | 0 |
| | NTRK2 | 1.042 | 0.446 | 0.042 | 0 |
| | MMP3 | 1.237 | 0.333 | 0.176 | 0 |
| | CXCL8 | 1.158 | 0.444 | 0.316 | 0 |
| | CXCL1 | 1.140 | 0.606 | 0.337 | 0 |
| | CXCL3 | 1.081 | 0.459 | 0.253 | 0 |
| | IL6 | 0.976 | 0.429 | 0.263 | 0 |
| | CXCL2 | 0.935 | 0.637 | 0.402 | 0 |
| | MMP1 | 0.865 | 0.113 | 0.060 | 3.96E-208 |
| | SAT1 | 0.756 | 0.932 | 0.843 | 0 |
| | SERPINE2 | 0.701 | 0.569 | 0.413 | 0 |
| | GEM | 0.680 | 0.891 | 0.662 | 0 |
| Fibro-immune | CEBPB | 0.608 | 0.935 | 0.889 | 0 |
| | KDM6B | 0.605 | 0.719 | 0.569 | 0 |
| | MMP10 | 0.604 | 0.113 | 0.059 | 1.04E-209 |
| | RSPO3 | 0.596 | 0.356 | 0.211 | 0 |
| | ARSG | 0.589 | 0.342 | 0.187 | 0 |
| | TNFAIP6 | 0.584 | 0.928 | 0.727 | 0 |
| | CREM | 0.573 | 0.404 | 0.238 | 0 |
| | THBS1 | 0.570 | 0.534 | 0.354 | 0 |
| | IER3 | 0.566 | 0.711 | 0.553 | 0 |
| | SFRP4 | 0.560 | 0.191 | 0.134 | 2.53E-164 |

| Cell state | Gene | Average LogFC | pct1 | pct2 | Adjusted P-value |
|---|---|---|---|---|---|
| | | **Vascular cells** | | | |
| | IGFBP3 | 2.086 | 0.678 | 0.170 | 0 |
| | CXCL3 | 1.644 | 0.438 | 0.223 | 0 |
| | HEY1 | 1.502 | 0.622 | 0.107 | 0 |
| | CLDN14 | 1.333 | 0.386 | 0.059 | 0 |
| | TCIM | 1.293 | 0.426 | 0.196 | 0 |
| | RASD1 | 1.232 | 0.440 | 0.146 | 0 |
| | CREM | 1.228 | 0.616 | 0.419 | 0 |
| | CD55 | 1.224 | 0.689 | 0.587 | 0 |
| | PLPP1 | 1.211 | 0.585 | 0.224 | 0 |
| Vas-arterial | GADD45B | 1.197 | 0.619 | 0.510 | 5.11E-192 |
| | SAT1 | 1.130 | 0.939 | 0.878 | 0 |
| | FN1 | 1.116 | 0.410 | 0.123 | 0 |
| | FOS | 1.050 | 0.506 | 0.364 | 2.57E-195 |
| | SERPINE2 | 1.046 | 0.272 | 0.041 | 0 |
| | CXCL8 | 1.036 | 0.590 | 0.445 | 9.46E-221 |
| | RND3 | 1.032 | 0.391 | 0.130 | 0 |
| | DUSP1 | 1.029 | 0.720 | 0.548 | 0 |
| | HIST1H1C | 1.012 | 0.278 | 0.116 | 1.69E-298 |
| | SRGN | 0.986 | 0.923 | 0.782 | 0 |
| | CXCL12 | 0.937 | 0.376 | 0.145 | 0 |
| | RGCC | 1.145 | 0.575 | 0.177 | 0 |
| | COL4A1 | 1.143 | 0.772 | 0.442 | 0 |
| | BTNL9 | 1.120 | 0.356 | 0.067 | 0 |
| | SOX18 | 1.102 | 0.602 | 0.225 | 0 |
| | CA4 | 1.017 | 0.233 | 0.039 | 0 |
| | ADGRF5 | 0.955 | 0.646 | 0.221 | 0 |
| | FABP4 | 0.920 | 0.689 | 0.599 | 5.14E-246 |
| | RBP7 | 0.902 | 0.632 | 0.462 | 0 |
| | C11orf96 | 0.892 | 0.444 | 0.239 | 0 |
| Vas-capillary | APOD | 0.886 | 0.588 | 0.385 | 0 |
| | COL4A2 | 0.882 | 0.692 | 0.380 | 0 |
| | CD36 | 0.869 | 0.465 | 0.363 | 3.01E-221 |
| | MCAM | 0.706 | 0.598 | 0.434 | 0 |
| | CD300LG | 0.704 | 0.218 | 0.043 | 0 |
| | RFLNB | 0.689 | 0.415 | 0.106 | 0 |
| | PXDN | 0.667 | 0.536 | 0.259 | 0 |
| | MT1M | 0.643 | 0.297 | 0.218 | 5.04E-120 |
| | F2RL3 | 0.633 | 0.437 | 0.240 | 0 |
| | SPARC | 0.622 | 0.807 | 0.660 | 0 |
| | GSN | 0.609 | 0.771 | 0.624 | 0 |
| | SELE | 2.345 | 0.730 | 0.160 | 0 |
| | ACKR1 | 2.209 | 0.883 | 0.216 | 0 |
| | IL6 | 1.780 | 0.653 | 0.222 | 0 |
| | ADIRF | 1.398 | 0.880 | 0.451 | 0 |
| | CSF3 | 1.354 | 0.672 | 0.286 | 0 |
| | CLU | 1.318 | 0.799 | 0.382 | 0 |
| | G0S2 | 1.223 | 0.401 | 0.194 | 0 |
| | VCAM1 | 1.076 | 0.381 | 0.053 | 0 |
| | HLA-DRA | 1.024 | 0.890 | 0.501 | 0 |
| Vas-venous | IL1R1 | 1.016 | 0.641 | 0.193 | 0 |
| | AQP1 | 0.993 | 0.734 | 0.232 | 0 |
| | SELP | 0.937 | 0.518 | 0.060 | 0 |
| | CYP1B1 | 0.921 | 0.283 | 0.141 | 0 |
| | MED24 | 0.906 | 0.451 | 0.165 | 0 |
| | CD74 | 0.901 | 0.957 | 0.771 | 0 |
| | PLCG2 | 0.888 | 0.304 | 0.274 | 1.21E-10 |
| | CST3 | 0.885 | 0.886 | 0.671 | 0 |
| | S100A10 | 0.857 | 0.923 | 0.641 | 0 |
| | CNKSR3 | 0.855 | 0.620 | 0.240 | 0 |
| | HLA-DPB1 | 0.818 | 0.789 | 0.409 | 0 |

#### Supplementary Table 10: Top Gene Markers Expressed for Breast Cell States

| Cell state | Gene | Average LogFC | pct1 | pct2 | Adjusted P-value |
|---|---|---|---|---|---|
| | | Lymphatic cells | | | |
| **Lymph-major** | CCL21 | 2.074 | 0.937 | 0.465 | 3.23E-185 |
| | FABP4 | 2.056 | 0.867 | 0.277 | 2.02E-181 |
| | LYVE1 | 1.928 | 0.736 | 0.081 | 2.44E-159 |
| | FABP5 | 1.344 | 0.848 | 0.434 | 5.80E-124 |
| | CYR61 | 1.272 | 0.554 | 0.251 | 2.21E-48 |
| | PRSS23 | 1.207 | 0.671 | 0.222 | 7.81E-95 |
| | NRP2 | 1.028 | 0.697 | 0.367 | 1.30E-71 |
| | IGF1 | 1.007 | 0.450 | 0.130 | 1.15E-47 |
| | MMRN1 | 1.002 | 0.929 | 0.680 | 1.61E-125 |
| | RGS16 | 0.979 | 0.622 | 0.373 | 2.65E-41 |
| | PLCG2 | 0.837 | 0.399 | 0.344 | 1 |
| | LOX | 0.832 | 0.383 | 0.072 | 6.81E-45 |
| | PDPN | 0.815 | 0.656 | 0.320 | 1.00E-57 |
| | FN1 | 0.814 | 0.520 | 0.259 | 1.66E-34 |
| | GNG11 | 0.781 | 0.910 | 0.832 | 2.00E-67 |
| | TM4SF18 | 0.754 | 0.497 | 0.212 | 8.57E-38 |
| | CLEC2B | 0.749 | 0.489 | 0.242 | 1.48E-29 |
| | CTGF | 0.739 | 0.392 | 0.253 | 1.24E-10 |
| | CYTOR | 0.734 | 0.492 | 0.253 | 9.26E-30 |
| | MRC1 | 0.716 | 0.524 | 0.266 | 4.25E-34 |
| **Lymph-valve** | SCG3 | 2.520 | 0.586 | 0.043 | 0 |
| | CD24 | 1.698 | 0.458 | 0.111 | 5.77E-100 |
| | HGF | 1.190 | 0.384 | 0.019 | 2.06E-303 |
| | GJA4 | 1.153 | 0.411 | 0.032 | 2.01E-245 |
| | SLC41A1 | 1.145 | 0.618 | 0.267 | 2.84E-72 |
| | ODC1 | 1.145 | 0.783 | 0.415 | 1.22E-88 |
| | CALM1 | 1.134 | 0.906 | 0.769 | 5.45E-66 |
| | TTN | 1.045 | 0.557 | 0.179 | 5.48E-81 |
| | CLDN11 | 1.038 | 0.451 | 0.027 | 0 |
| | ADAMTS1 | 0.846 | 0.320 | 0.104 | 2.26E-38 |
| | SBSPON | 0.833 | 0.399 | 0.059 | 8.42E-140 |
| | NTS | 0.816 | 0.224 | 0.102 | 3.84E-11 |
| | SOCS2 | 0.814 | 0.722 | 0.389 | 7.13E-63 |
| | GSTM3 | 0.809 | 0.490 | 0.168 | 1.37E-66 |
| | ADM | 0.802 | 0.633 | 0.433 | 5.66E-19 |
| | FOXC2 | 0.791 | 0.515 | 0.235 | 6.06E-42 |
| | PROX1 | 0.772 | 0.865 | 0.607 | 1.98E-54 |
| | NUDT9 | 0.752 | 0.426 | 0.131 | 1.45E-64 |
| | MYLIP | 0.750 | 0.512 | 0.170 | 9.71E-71 |
| | KLF2 | 0.737 | 0.845 | 0.585 | 1.58E-43 |
| **Lymph-immune** | NTS | 2.817 | 0.748 | 0.095 | 3.19E-153 |
| | CXCL1 | 2.124 | 0.483 | 0.159 | 9.55E-24 |
| | CXCL2 | 2.110 | 0.714 | 0.285 | 1.25E-37 |
| | CLU | 2.069 | 0.925 | 0.516 | 5.15E-48 |
| | IGFBP5 | 2.066 | 0.313 | 0.092 | 1.83E-16 |
| | CXCL8 | 2.016 | 0.551 | 0.268 | 1.21E-14 |
| | CXCL3 | 1.753 | 0.395 | 0.118 | 4.39E-21 |
| | PTGS2 | 1.669 | 0.578 | 0.031 | 9.25E-239 |
| | CSF3 | 1.611 | 0.429 | 0.063 | 6.48E-65 |
| | CCL2 | 1.589 | 0.633 | 0.352 | 8.90E-13 |
| | PLAT | 1.551 | 0.551 | 0.074 | 3.99E-99 |
| | TGM2 | 1.485 | 0.653 | 0.066 | 5.92E-158 |
| | SERPINB1 | 1.370 | 0.612 | 0.167 | 5.45E-49 |
| | AKR1C2 | 1.314 | 0.551 | 0.243 | 5.57E-17 |
| | IL6 | 1.261 | 0.279 | 0.055 | 9.34E-27 |
| | MT1X | 1.211 | 0.741 | 0.335 | 3.88E-25 |
| | SAT1 | 1.166 | 0.830 | 0.639 | 1.13E-13 |
| | PRELP | 1.138 | 0.401 | 0.027 | 5.22E-130 |
| | THBS4 | 1.104 | 0.442 | 0.048 | 1.34E-93 |
| | SRGN | 1.095 | 0.531 | 0.091 | 7.73E-69 |

| Cell state | Gene | Average LogFC | pct1 | pct2 | Adjusted P-value |
|---|---|---|---|---|---|
| | | Pericytes | | | |
| **Peri-immune** | SERPINE1 | 1.925 | 0.759 | 0.158 | 0 |
| | CXCL3 | 1.849 | 0.947 | 0.491 | 0 |
| | CYP1B1 | 1.639 | 0.928 | 0.556 | 0 |
| | COL6A3 | 1.509 | 0.227 | 0.038 | 0 |
| | CXCL8 | 1.354 | 0.876 | 0.528 | 0 |
| | CCL20 | 1.185 | 0.487 | 0.074 | 0 |
| | AKR1C1 | 1.103 | 0.725 | 0.183 | 0 |
| | PLIN2 | 1.088 | 0.56 | 0.122 | 0 |
| | CTSC | 1.071 | 0.804 | 0.382 | 0 |
| | EDNRB | 1.041 | 0.943 | 0.642 | 0 |
| | MEDAG | 1.009 | 0.624 | 0.187 | 0 |
| | TFPI | 0.998 | 0.97 | 0.737 | 0 |
| | PAPPA | 0.977 | 0.569 | 0.15 | 0 |
| | SOD2 | 0.952 | 0.649 | 0.266 | 0 |
| | HMOX1 | 0.889 | 0.87 | 0.572 | 0 |
| | TIMP1 | 0.815 | 0.145 | 0.058 | 0 |
| | FTH1 | 0.807 | 0.657 | 0.288 | 0 |
| | MARCKS | 0.804 | 0.312 | 0.006 | 0 |
| | FGF7 | 0.785 | 0.748 | 0.51 | 0 |
| | COL4A1 | 0.785 | 0.698 | 0.375 | 0 |
| **Peri-HR** | CREM | 1.847 | 0.878 | 0.284 | 0 |
| | CMSS1 | 1.725 | 0.698 | 0.266 | 0 |
| | GCNA | 1.334 | 0.128 | 0.034 | 5.22E-196 |
| | C11orf96 | 1.304 | 0.975 | 0.855 | 0 |
| | IL24 | 1.293 | 0.149 | 0.016 | 0 |
| | CNST | 1.186 | 0.28 | 0.104 | 0 |
| | ATP1B3 | 1.150 | 0.805 | 0.547 | 0 |
| | PDK4 | 1.118 | 0.74 | 0.514 | 0 |
| | CRYAB | 1.109 | 0.81 | 0.433 | 0 |
| | EIF4A3 | 1.088 | 0.752 | 0.454 | 0 |
| | PNP | 1.074 | 0.623 | 0.148 | 0 |
| | PGAP1 | 1.069 | 0.481 | 0.071 | 0 |
| | PTP4A1 | 1.023 | 0.689 | 0.265 | 0 |
| | SRL | 1.001 | 0.415 | 0.034 | 0 |
| | SLC25A4 | 0.998 | 0.607 | 0.217 | 0 |
| | CHMP1B | 0.980 | 0.545 | 0.228 | 0 |
| | HIST1H1C | 0.976 | 0.373 | 0.131 | 0 |
| | YBX3 | 0.971 | 0.952 | 0.798 | 0 |
| | BCAS2 | 0.944 | 0.633 | 0.249 | 0 |
| | ZNF331 | 0.940 | 0.541 | 0.239 | 0 |
| **Peri-myo** | PLCG2 | 2.142 | 0.364 | 0.241 | 9.09E-191 |
| | MYL9 | 1.614 | 0.853 | 0.444 | 0 |
| | ACTA2 | 1.552 | 0.758 | 0.35 | 0 |
| | ZFP36 | 1.526 | 0.747 | 0.488 | 0 |
| | FOS | 1.489 | 0.775 | 0.477 | 0 |
| | FABP4 | 1.472 | 0.711 | 0.393 | 0 |
| | TAGLN | 1.443 | 0.896 | 0.569 | 0 |
| | JUN | 1.443 | 0.839 | 0.74 | 0 |
| | S100A4 | 1.398 | 0.76 | 0.371 | 0 |
| | RGS16 | 1.395 | 0.566 | 0.244 | 0 |
| | ADIRF | 1.349 | 0.901 | 0.673 | 0 |
| | HSPA1B | 1.337 | 0.565 | 0.316 | 0 |
| | TPM1 | 1.294 | 0.816 | 0.448 | 0 |
| | DNAJB1 | 1.282 | 0.574 | 0.334 | 0 |
| | PLAC9 | 1.248 | 0.635 | 0.248 | 0 |
| | EGR1 | 1.245 | 0.61 | 0.195 | 0 |
| | HSPA1A | 1.223 | 0.688 | 0.522 | 0 |
| | CRIP1 | 1.206 | 0.471 | 0.139 | 0 |
| | SPARCL1 | 1.193 | 0.876 | 0.646 | 0 |
| | FOSB | 1.177 | 0.622 | 0.324 | 0 |

**Supplementary Table 10: Top Gene Markers Expressed for Breast Cell States**

| Cell state | Gene | Average LogFC | pct1 | pct2 | Adjusted P-value |
|---|---|---|---|---|---|
| | | B cells | | | |
| Bnaive | HLA-DRA | 1.557 | 0.971 | 0.317 | 0 |
| | CD37 | 1.519 | 0.911 | 0.336 | 0 |
| | HLA-DRB1 | 1.514 | 0.929 | 0.275 | 0 |
| | HLA-DPB1 | 1.489 | 0.930 | 0.254 | 0 |
| | CD74 | 1.352 | 0.984 | 0.831 | 0 |
| | HLA-DPA1 | 1.322 | 0.891 | 0.298 | 0 |
| | HLA-DQB1 | 1.319 | 0.799 | 0.259 | 0 |
| | LAPTM5 | 1.273 | 0.862 | 0.319 | 0 |
| | SMCHD1 | 1.261 | 0.753 | 0.453 | 0 |
| | YBX3 | 1.217 | 0.411 | 0.226 | 5.02E-106 |
| | REL | 1.187 | 0.796 | 0.227 | 0 |
| | ATP1B3 | 1.183 | 0.562 | 0.426 | 6.01E-120 |
| | HLA-DQA2 | 1.129 | 0.407 | 0.082 | 2.24E-242 |
| | BCL11A | 1.128 | 0.531 | 0.117 | 0 |
| | BTG1 | 1.123 | 0.979 | 0.858 | 0 |
| | LTB | 1.115 | 0.737 | 0.182 | 0 |
| | HLA-DQA1 | 1.103 | 0.710 | 0.185 | 0 |
| | CEMIP2 | 1.087 | 0.417 | 0.128 | 1.99E-198 |
| | CREM | 1.067 | 0.506 | 0.338 | 8.14E-113 |
| | LYN | 1.050 | 0.559 | 0.204 | 4.11E-272 |
| Bmem-switched | S100A4 | 1.249 | 0.550 | 0.292 | 3.28E-88 |
| | CD52 | 0.985 | 0.867 | 0.362 | 1.19E-193 |
| | LTB | 0.859 | 0.801 | 0.321 | 3.71E-155 |
| | TMSB4X | 0.843 | 0.983 | 0.883 | 1.74E-137 |
| | MARCH1 | 0.816 | 0.544 | 0.118 | 4.82E-194 |
| | ARL6IP5 | 0.806 | 0.547 | 0.233 | 2.04E-107 |
| | RPL31 | 0.801 | 0.960 | 0.866 | 1.94E-115 |
| | COTL1 | 0.784 | 0.527 | 0.131 | 1.76E-167 |
| | RPS10 | 0.781 | 0.904 | 0.760 | 5.97E-127 |
| | RPL36A | 0.770 | 0.656 | 0.508 | 2.47E-39 |
| | SH3BGRL3 | 0.754 | 0.776 | 0.557 | 2.90E-95 |
| | MS4A1 | 0.728 | 0.758 | 0.298 | 7.74E-133 |
| | POU2F2 | 0.726 | 0.586 | 0.265 | 1.95E-89 |
| | RPS17 | 0.708 | 0.593 | 0.503 | 1.44E-16 |
| | RPL27A | 0.707 | 0.994 | 0.967 | 7.80E-178 |
| | BANK1 | 0.700 | 0.711 | 0.290 | 3.52E-120 |
| | RPLP2 | 0.692 | 0.999 | 0.983 | 7.55E-192 |
| | PFN1 | 0.691 | 0.730 | 0.560 | 8.41E-67 |
| | RPL13A | 0.689 | 1.000 | 0.988 | 2.63E-178 |
| | SSPN | 0.687 | 0.312 | 0.116 | 2.17E-60 |
| Bmem-unswitched | DUSP2 | 1.852 | 0.737 | 0.123 | 2.04E-181 |
| | CD83 | 1.816 | 0.947 | 0.266 | 3.32E-158 |
| | BCL2A1 | 1.776 | 0.728 | 0.082 | 6.51E-244 |
| | MYC | 1.707 | 0.700 | 0.149 | 7.92E-139 |
| | EGR3 | 1.672 | 0.576 | 0.031 | 0 |
| | CD69 | 1.565 | 0.765 | 0.194 | 4.16E-114 |
| | MIR155HG | 1.497 | 0.601 | 0.226 | 6.91E-55 |
| | ZFP36L1 | 1.467 | 0.918 | 0.437 | 2.77E-88 |
| | NFKBID | 1.435 | 0.728 | 0.197 | 5.38E-104 |
| | NR4A1 | 1.420 | 0.613 | 0.160 | 1.47E-89 |
| | REL | 1.370 | 0.963 | 0.407 | 4.25E-102 |
| | HLA-DQA1 | 1.253 | 0.877 | 0.351 | 4.95E-79 |
| | CCR7 | 1.219 | 0.704 | 0.228 | 1.28E-70 |
| | HSP90AB1 | 1.197 | 0.959 | 0.863 | 1.20E-76 |
| | NFKB1 | 1.167 | 0.720 | 0.287 | 2.58E-64 |
| | FABP5 | 1.159 | 0.383 | 0.101 | 4.25E-45 |
| | NR4A3 | 1.156 | 0.387 | 0.045 | 2.22E-115 |
| | KDM6B | 1.146 | 0.691 | 0.296 | 5.95E-57 |
| | NR4A2 | 1.124 | 0.535 | 0.169 | 2.20E-53 |
| | DDX21 | 1.101 | 0.827 | 0.465 | 6.06E-61 |

| Cell state | Gene | Average LogFC | pct1 | pct2 | Adjusted P-value |
|---|---|---|---|---|---|
| | | B cells | | | |
| Bplasma-IgA | IGHA2 | 3.559 | 0.824 | 0.193 | 0 |
| | IGHA1 | 3.430 | 0.953 | 0.538 | 0 |
| | JCHAIN | 3.346 | 0.946 | 0.333 | 0 |
| | IGHM | 2.161 | 0.258 | 0.541 | 4.65E-122 |
| | IGLL5 | 1.717 | 0.295 | 0.060 | 4.65E-149 |
| | IGLC2 | 1.691 | 0.641 | 0.427 | 7.75E-62 |
| | GADD45A | 1.576 | 0.683 | 0.164 | 0 |
| | HSPA5 | 1.506 | 0.900 | 0.332 | 0 |
| | IGLC3 | 1.430 | 0.520 | 0.356 | 5.75E-37 |
| | ANKRD28 | 1.392 | 0.925 | 0.173 | 0 |
| | ERN1 | 1.348 | 0.785 | 0.126 | 0 |
| | VIM | 1.342 | 0.980 | 0.697 | 0 |
| | IGKC | 1.312 | 0.889 | 0.831 | 2.48E-45 |
| | SSR4 | 1.295 | 0.985 | 0.583 | 0 |
| | DERL3 | 1.211 | 0.905 | 0.167 | 0 |
| | CCL4 | 1.206 | 0.111 | 0.029 | 1.51E-40 |
| | NEAT1 | 1.200 | 0.983 | 0.763 | 0 |
| | HSP90B1 | 1.192 | 0.971 | 0.548 | 0 |
| | CYTOR | 1.101 | 0.864 | 0.164 | 0 |
| | CCL3 | 1.078 | 0.135 | 0.031 | 1.24E-55 |
| Bplasma-IgG | IGHG1 | 4.451 | 0.874 | 0.175 | 0 |
| | IGHG4 | 3.900 | 0.749 | 0.068 | 0 |
| | IGHG3 | 3.713 | 0.832 | 0.151 | 0 |
| | IGHG2 | 3.375 | 0.579 | 0.072 | 0 |
| | IGHGP | 2.532 | 0.375 | 0.010 | 0 |
| | IGKC | 1.127 | 0.894 | 0.849 | 5.56E-39 |
| | MZB1 | 1.110 | 0.901 | 0.458 | 1.06E-141 |
| | PRDX4 | 0.976 | 0.785 | 0.416 | 3.86E-122 |
| | IGLC3 | 0.951 | 0.462 | 0.416 | 1 |
| | DNAAF1 | 0.904 | 0.351 | 0.140 | 8.16E-49 |
| | JSRP1 | 0.883 | 0.549 | 0.202 | 3.59E-112 |
| | SSR4 | 0.812 | 0.982 | 0.713 | 4.29E-122 |
| | FKBP11 | 0.806 | 0.855 | 0.461 | 3.85E-109 |
| | XBP1 | 0.760 | 0.806 | 0.483 | 1.64E-80 |
| | IGKV4-1 | 0.744 | 0.155 | 0.062 | 1.21E-16 |
| | ITM2C | 0.664 | 0.599 | 0.350 | 4.99E-49 |
| | IGLV3-1 | 0.653 | 0.303 | 0.113 | 2.29E-46 |
| | IGLL5 | 0.648 | 0.197 | 0.148 | 1 |
| | SLC3A2 | 0.635 | 0.715 | 0.494 | 3.20E-43 |
| | TYMP | 0.603 | 0.675 | 0.430 | 4.26E-50 |

##### Supplementary Table 10: Top Gene Markers Expressed for Breast Cell States

| Cell state | Gene | Average LogFC | pct1 | pct2 | Adjusted P-value |
|---|---|---|---|---|---|
| | | T cells | | | |
| CD4-Th | CCL20 | 0.261 | 0.033 | 0.000 | 0 |
| | KLRB1 | 0.752 | 0.422 | 0.000 | 0 |
| | CCR6 | 0.405 | 0.144 | 0.000 | 0 |
| | IL7R | 0.991 | 0.726 | 0.000 | 0 |
| | ODF2L | 0.546 | 0.321 | 0.000 | 0 |
| | SESN1 | 0.285 | 0.116 | 0.000 | 0 |
| | RORA | 0.760 | 0.535 | 0.000 | 0 |
| | PTPN13 | 0.187 | 0.021 | 0.000 | 0 |
| | RPLP0 | 0.982 | 0.922 | 0.000 | 0 |
| | CEBPD | 0.353 | 0.187 | 0.000 | 5.02E-106 |
| | RPS17 | 0.502 | 0.375 | 0.000 | 0 |
| | FURIN | 0.278 | 0.095 | 0.000 | 6.01E-120 |
| | MCAM | 0.188 | 0.036 | 0.000 | 2.24E-242 |
| | RPLP1 | 1.000 | 0.997 | 0.000 | 0 |
| | TNFRSF25 | 0.279 | 0.105 | 0.000 | 0 |
| | GPR171 | 0.427 | 0.282 | 0.000 | 0 |
| | TNFSF13B | 0.138 | 0.016 | 0.000 | 0 |
| | ERN1 | 0.416 | 0.233 | 0.000 | 1.99E-198 |
| | IL4I1 | 0.143 | 0.012 | 0.000 | 8.14E-113 |
| | RPL36A | 0.618 | 0.532 | 0.000 | 4.11E-272 |
| CD4-Thlike | MAF | 1.045 | 0.538 | 0.229 | 0 |
| | PGAP1 | 0.885 | 0.324 | 0.074 | 0 |
| | CH25H | 0.808 | 0.166 | 0.029 | 0 |
| | SOCS3 | 0.772 | 0.486 | 0.179 | 0 |
| | CMSS1 | 0.766 | 0.325 | 0.137 | 5.34E-302 |
| | FTH1 | 0.723 | 0.994 | 0.978 | 0 |
| | CRYBG1 | 0.713 | 0.689 | 0.410 | 0 |
| | ATP1B3 | 0.696 | 0.766 | 0.520 | 0 |
| | SRGN | 0.692 | 0.997 | 0.933 | 0 |
| | GPR183 | 0.692 | 0.665 | 0.370 | 0 |
| | SESN3 | 0.680 | 0.336 | 0.128 | 0 |
| | ARID5B | 0.670 | 0.675 | 0.359 | 0 |
| | NR3C1 | 0.646 | 0.728 | 0.439 | 0 |
| | SLC2A3 | 0.628 | 0.630 | 0.354 | 0 |
| | DDIT4 | 0.608 | 0.794 | 0.573 | 0 |
| | CREM | 0.589 | 0.882 | 0.580 | 0 |
| | GLIPR1 | 0.578 | 0.687 | 0.418 | 0 |
| | ZFP36 | 0.576 | 0.819 | 0.631 | 0 |
| | UGP2 | 0.563 | 0.524 | 0.286 | 0 |
| | AP3M2 | 0.543 | 0.358 | 0.135 | 0 |
| CD4-naive | LTB | 1.086 | 0.800 | 0.308 | 0 |
| | KLF2 | 1.037 | 0.816 | 0.275 | 0 |
| | SELL | 0.792 | 0.393 | 0.058 | 0 |
| | CCR7 | 0.691 | 0.406 | 0.110 | 0 |
| | LDHB | 0.639 | 0.794 | 0.421 | 0 |
| | SESN3 | 0.612 | 0.400 | 0.144 | 1.66E-274 |
| | RPS8 | 0.499 | 0.999 | 0.988 | 0 |
| | RIPOR2 | 0.471 | 0.460 | 0.190 | 4.87E-235 |
| | RPL23 | 0.463 | 0.976 | 0.884 | 0 |
| | LEF1 | 0.453 | 0.260 | 0.046 | 0 |
| | NOSIP | 0.449 | 0.412 | 0.169 | 5.75E-210 |
| | RPL18A | 0.437 | 0.999 | 0.990 | 0 |
| | RPL29 | 0.430 | 0.996 | 0.976 | 0 |
| | RPS12 | 0.427 | 1.000 | 0.994 | 0 |
| | RPS6 | 0.417 | 0.999 | 0.991 | 0 |
| | RPL13 | 0.413 | 1.000 | 0.998 | 0 |
| | MAL | 0.410 | 0.246 | 0.046 | 0 |
| | RPL18 | 0.408 | 0.997 | 0.985 | 0 |
| | EEF1A1 | 0.406 | 1.000 | 0.998 | 0 |
| | RPS13 | 0.404 | 0.999 | 0.982 | 0 |

| Cell state | Gene | Average LogFC | pct1 | pct2 | Adjusted P-value |
|---|---|---|---|---|---|
| | | T cells | | | |
| CD4-Tem | HSPA1A | 3.559 | 0.824 | 0.193 | 0 |
| | FOS | 3.430 | 0.953 | 0.538 | 0 |
| | LMNA | 3.346 | 0.946 | 0.333 | 0 |
| | KLF6 | 2.161 | 0.258 | 0.541 | 4.65E-122 |
| | FOSB | 1.717 | 0.295 | 0.060 | 4.65E-149 |
| | HSPA1B | 1.691 | 0.641 | 0.427 | 7.75E-62 |
| | AHNAK | 1.576 | 0.683 | 0.164 | 0 |
| | CRIP1 | 1.506 | 0.900 | 0.332 | 0 |
| | ANXA1 | 1.430 | 0.520 | 0.356 | 5.75E-37 |
| | DNAJB1 | 1.392 | 0.925 | 0.173 | 0 |
| | ANKRD28 | 1.348 | 0.785 | 0.126 | 0 |
| | TNFAIP3 | 1.342 | 0.980 | 0.697 | 0 |
| | JUN | 1.312 | 0.889 | 0.831 | 2.48E-45 |
| | RGCC | 1.295 | 0.985 | 0.583 | 0 |
| | IGLC2 | 1.211 | 0.905 | 0.167 | 0 |
| | IGKC | 1.206 | 0.111 | 0.029 | 1.51E-40 |
| | PPP1R15A | 1.200 | 0.983 | 0.763 | 0 |
| | MYADM | 1.192 | 0.971 | 0.548 | 0 |
| | TSC22D3 | 1.101 | 0.864 | 0.164 | 0 |
| | S100A4 | 1.078 | 0.135 | 0.031 | 1.24E-55 |
| CD4-Treg | TIGIT | 1.349 | 0.634 | 0.128 | 0 |
| | RTKN2 | 1.337 | 0.415 | 0.022 | 0 |
| | CTLA4 | 1.299 | 0.549 | 0.063 | 0 |
| | CARD16 | 1.213 | 0.568 | 0.131 | 1.44E-286 |
| | TBC1D4 | 1.195 | 0.512 | 0.062 | 0 |
| | BATF | 1.142 | 0.588 | 0.197 | 8.09E-182 |
| | HPGD | 1.124 | 0.301 | 0.062 | 3.49E-155 |
| | LTB | 1.072 | 0.694 | 0.342 | 1.02E-123 |
| | PMAIP1 | 1.067 | 0.455 | 0.113 | 1.07E-193 |
| | IL32 | 1.063 | 0.947 | 0.673 | 1.83E-176 |
| | IKZF2 | 1.005 | 0.427 | 0.043 | 0 |
| | TNFRSF18 | 0.958 | 0.408 | 0.105 | 4.73E-159 |
| | FOXP3 | 0.939 | 0.334 | 0.003 | 0 |
| | IL2RA | 0.921 | 0.338 | 0.044 | 2.83E-295 |
| | UGP2 | 0.906 | 0.670 | 0.321 | 8.77E-121 |
| | TNFRSF4 | 0.895 | 0.304 | 0.089 | 3.04E-89 |
| | CD27 | 0.889 | 0.454 | 0.116 | 4.00E-180 |
| | DUSP4 | 0.865 | 0.469 | 0.174 | 2.60E-105 |
| | TNFRSF9 | 0.854 | 0.342 | 0.050 | 6.34E-258 |
| | LINC01943 | 0.847 | 0.329 | 0.050 | 4.21E-237 |
| CD4-activated | PLCG2 | 3.032 | 0.936 | 0.193 | 0 |
| | HIST1H4C | 1.525 | 0.439 | 0.346 | 2.75E-08 |
| | LTB | 1.165 | 0.710 | 0.344 | 5.20E-91 |
| | HSPA1B | 1.160 | 0.226 | 0.102 | 5.49E-17 |
| | RMRP | 1.130 | 0.295 | 0.016 | 0 |
| | TRAF3IP3 | 0.923 | 0.525 | 0.195 | 2.40E-78 |
| | LINC00324 | 0.909 | 0.188 | 0.046 | 5.73E-43 |
| | INPP4B | 0.902 | 0.373 | 0.116 | 6.79E-69 |
| | CD40LG | 0.855 | 0.392 | 0.106 | 1.46E-88 |
| | TRAC | 0.769 | 0.727 | 0.481 | 8.97E-54 |
| | SLFN5 | 0.756 | 0.572 | 0.333 | 4.50E-38 |
| | CD52 | 0.734 | 0.794 | 0.583 | 2.43E-50 |
| | ATM | 0.712 | 0.557 | 0.328 | 2.68E-35 |
| | CD3G | 0.691 | 0.732 | 0.456 | 6.15E-54 |
| | LINC00861 | 0.688 | 0.255 | 0.080 | 3.00E-40 |
| | MTRNR2L12 | 0.685 | 0.743 | 0.571 | 3.92E-32 |
| | CHURC1 | 0.671 | 0.446 | 0.225 | 6.39E-34 |
| | N4BP2L2 | 0.665 | 0.818 | 0.643 | 3.41E-50 |
| | SCML4 | 0.653 | 0.306 | 0.123 | 4.12E-32 |
| | GSTK1 | 0.648 | 0.521 | 0.323 | 6.65E-30 |

**Supplementary Table 10: Top Gene Markers Expressed for Breast Cell States**

| Cell state | Gene | Average LogFC | pct1 | pct2 | Adjusted P-value |
|---|---|---|---|---|---|
| | | T cells | | | |
| | PLCG2 | 2.947 | 0.908 | 0.183 | 0 |
| | CCL5 | 1.425 | 0.984 | 0.615 | 0 |
| | IGKC | 1.330 | 0.575 | 0.385 | 2.57E-60 |
| | HIST1H4C | 1.289 | 0.415 | 0.345 | 6.45E-17 |
| | ZNF683 | 1.122 | 0.306 | 0.029 | 0 |
| | ITGA1 | 1.074 | 0.428 | 0.139 | 3.43E-167 |
| | TRG-AS1 | 1.006 | 0.360 | 0.090 | 4.81E-198 |
| | RMRP | 0.970 | 0.256 | 0.013 | 0 |
| | TRAF3IP3 | 0.952 | 0.499 | 0.191 | 2.20E-163 |
| CD8-activated | HSPA1B | 0.937 | 0.193 | 0.101 | 8.05E-21 |
| | CD8B | 0.920 | 0.462 | 0.176 | 1.31E-139 |
| | TRAC | 0.916 | 0.716 | 0.478 | 1.99E-133 |
| | LINC00324 | 0.892 | 0.188 | 0.044 | 1.39E-99 |
| | TRGC2 | 0.866 | 0.398 | 0.171 | 2.73E-91 |
| | CD3G | 0.854 | 0.702 | 0.453 | 2.61E-131 |
| | GPR174 | 0.851 | 0.342 | 0.111 | 2.70E-127 |
| | JCHAIN | 0.844 | 0.141 | 0.097 | 0.00479643 |
| | KLRC1 | 0.838 | 0.299 | 0.133 | 1.04E-53 |
| | CD2 | 0.828 | 0.873 | 0.696 | 9.22E-167 |
| | KLRC4 | 0.816 | 0.232 | 0.046 | 6.24E-156 |
| | KLRC1 | 1.091 | 0.396 | 0.068 | 0 |
| | CD8A | 1.075 | 0.727 | 0.184 | 0 |
| | ITGA1 | 0.884 | 0.415 | 0.075 | 0 |
| | SYTL3 | 0.744 | 0.812 | 0.526 | 0 |
| | AUTS2 | 0.728 | 0.424 | 0.155 | 0 |
| | XCL1 | 0.727 | 0.246 | 0.083 | 0 |
| | SPRY1 | 0.675 | 0.311 | 0.093 | 0 |
| | CD8B | 0.667 | 0.435 | 0.116 | 0 |
| | PGK1 | 0.667 | 0.674 | 0.461 | 0 |
| CD8-Trm | LINC01871 | 0.653 | 0.545 | 0.246 | 0 |
| | METRNL | 0.634 | 0.619 | 0.343 | 0 |
| | LDLRAD4 | 0.613 | 0.277 | 0.103 | 0 |
| | GABARAPL1 | 0.602 | 0.570 | 0.318 | 0 |
| | CD7 | 0.575 | 0.765 | 0.503 | 0 |
| | TSPYL2 | 0.562 | 0.535 | 0.366 | 1.94E-248 |
| | PARP8 | 0.538 | 0.618 | 0.400 | 0 |
| | RGS1 | 0.531 | 0.597 | 0.367 | 0 |
| | SLA2 | 0.527 | 0.276 | 0.093 | 0 |
| | PARD6G | 0.520 | 0.202 | 0.032 | 0 |
| | SYTL2 | 0.511 | 0.366 | 0.183 | 1.77E-296 |
| | GZMK | 1.735 | 0.794 | 0.131 | 0 |
| | HLA-DRB1 | 1.143 | 0.444 | 0.145 | 0 |
| | TNFRSF9 | 1.105 | 0.266 | 0.036 | 0 |
| | CRTAM | 1.047 | 0.336 | 0.068 | 0 |
| | SH2D1A | 0.939 | 0.484 | 0.171 | 0 |
| | CMC1 | 0.917 | 0.438 | 0.161 | 0 |
| | CST7 | 0.907 | 0.822 | 0.456 | 0 |
| | CD74 | 0.847 | 0.601 | 0.340 | 0 |
| | IGLC3 | 0.791 | 0.109 | 0.085 | 0.01151389 |
| CD8-Tem | HLA-DRA | 0.770 | 0.308 | 0.124 | 1.46E-196 |
| | HLA-DPB1 | 0.750 | 0.488 | 0.209 | 0 |
| | TIGIT | 0.745 | 0.349 | 0.119 | 7.78E-299 |
| | LYST | 0.736 | 0.419 | 0.163 | 0 |
| | DUSP2 | 0.702 | 0.732 | 0.474 | 3.60E-295 |
| | TUBA4A | 0.667 | 0.605 | 0.438 | 2.31E-157 |
| | HLA-DPA1 | 0.633 | 0.385 | 0.171 | 1.30E-219 |
| | CEMIP2 | 0.629 | 0.627 | 0.451 | 1.54E-168 |
| | EOMES | 0.613 | 0.226 | 0.041 | 0 |
| | CCL4L2 | 0.602 | 0.226 | 0.113 | 7.13E-73 |
| | HLA-DQB1 | 0.591 | 0.236 | 0.078 | 1.89E-196 |

| Cell state | Gene | Average LogFC | pct1 | pct2 | Adjusted P-value |
|---|---|---|---|---|---|
| | | T cells | | | |
| | NKG7 | 1.883 | 0.873 | 0.126 | 0 |
| | FGFBP2 | 1.623 | 0.977 | 0.298 | 0 |
| | GNLY | 1.482 | 0.697 | 0.068 | 0 |
| | CCL4 | 1.333 | 0.838 | 0.168 | 0 |
| | CD52 | 1.160 | 0.811 | 0.275 | 0 |
| | GZMA | 1.082 | 0.857 | 0.565 | 0 |
| | ZEB2 | 1.009 | 0.775 | 0.276 | 0 |
| | GZMB | 0.995 | 0.641 | 0.177 | 0 |
| | KLRD1 | 0.966 | 0.769 | 0.158 | 0 |
| NKT | KLRG1 | 0.917 | 0.768 | 0.251 | 0 |
| | TRGC2 | 0.893 | 0.446 | 0.079 | 0 |
| | C12orf75 | 0.893 | 0.507 | 0.153 | 0 |
| | HCST | 0.869 | 0.550 | 0.162 | 0 |
| | ACTB | 0.854 | 0.821 | 0.518 | 0 |
| | CCL5 | 0.826 | 0.942 | 0.869 | 1.85E-283 |
| | HLA-DPB1 | 0.802 | 0.976 | 0.599 | 0 |
| | EFHD2 | 0.801 | 0.595 | 0.207 | 0 |
| | FCGR3A | 0.769 | 0.618 | 0.248 | 0 |
| | CTSW | 0.769 | 0.420 | 0.074 | 0 |
| | S100A4 | 0.756 | 0.652 | 0.235 | 0 |
| | AREG | 1.702 | 0.811 | 0.311 | 1.06E-299 |
| | XCL2 | 1.685 | 0.671 | 0.115 | 0 |
| | TYROBP | 1.587 | 0.832 | 0.105 | 0 |
| | XCL1 | 1.503 | 0.676 | 0.105 | 0 |
| | FCER1G | 1.438 | 0.827 | 0.079 | 0 |
| | GNLY | 1.368 | 0.629 | 0.205 | 1.77E-199 |
| | TNFRSF18 | 1.294 | 0.568 | 0.101 | 0 |
| | CMC1 | 1.251 | 0.581 | 0.177 | 1.10E-203 |
| | CCL3 | 1.148 | 0.247 | 0.087 | 1.83E-53 |
| NK/ILCs | CTSW | 1.062 | 0.720 | 0.253 | 4.77E-220 |
| | REL | 1.038 | 0.706 | 0.444 | 7.00E-87 |
| | NFKB1 | 1.009 | 0.565 | 0.242 | 9.45E-122 |
| | KLRC1 | 1.000 | 0.558 | 0.128 | 5.25E-254 |
| | CSF2 | 0.980 | 0.132 | 0.004 | 0 |
| | TRDC | 0.978 | 0.414 | 0.051 | 0 |
| | KLRD1 | 0.946 | 0.737 | 0.277 | 3.82E-191 |
| | NFKBIA | 0.945 | 0.803 | 0.557 | 4.93E-83 |
| | GZMK | 0.912 | 0.596 | 0.180 | 1.96E-184 |
| | CCL4 | 0.888 | 0.417 | 0.309 | 2.62E-10 |
| | IER2 | 0.857 | 0.714 | 0.468 | 2.57E-74 |
| | GNLY | 2.416 | 0.957 | 0.149 | 0 |
| | CCL3 | 2.230 | 0.477 | 0.057 | 0 |
| | NKG7 | 2.132 | 0.982 | 0.289 | 0 |
| | GZMB | 2.066 | 0.910 | 0.138 | 0 |
| | TYROBP | 2.029 | 0.831 | 0.059 | 0 |
| | FCGR3A | 1.971 | 0.779 | 0.038 | 0 |
| | FCER1G | 1.966 | 0.740 | 0.039 | 0 |
| | FGFBP2 | 1.937 | 0.738 | 0.056 | 0 |
| | SPON2 | 1.877 | 0.697 | 0.094 | 0 |
| NK | PRF1 | 1.805 | 0.839 | 0.188 | 0 |
| | KLRF1 | 1.774 | 0.647 | 0.023 | 0 |
| | CCL4 | 1.767 | 0.833 | 0.266 | 0 |
| | KLRD1 | 1.695 | 0.924 | 0.231 | 0 |
| | CLIC3 | 1.617 | 0.614 | 0.077 | 0 |
| | CMC1 | 1.554 | 0.652 | 0.145 | 0 |
| | CTSW | 1.545 | 0.771 | 0.219 | 0 |
| | HBB | 1.448 | 0.138 | 0.041 | 1.41E-119 |
| | GZMA | 1.359 | 0.817 | 0.266 | 0 |
| | CCL4L2 | 1.356 | 0.359 | 0.102 | 0 |
| | CD247 | 1.263 | 0.757 | 0.301 | 0 |

**Supplementary Table 10: Top Gene Markers Expressed for Breast Cell States**

| Cell state | Gene | Average LogFC | pct1 | pct2 | Adjusted P-value |
|---|---|---|---|---|---|
| | | T cells | | | |
| GD-T | XCL1 | 1.105 | 0.372 | 0.116 | 3.99E-29 |
| | CEMIP2 | 1.028 | 0.783 | 0.465 | 5.12E-37 |
| | GPCPD1 | 1.007 | 0.588 | 0.162 | 7.60E-73 |
| | CCDC57 | 0.992 | 0.438 | 0.097 | 1.63E-69 |
| | METRNL | 0.947 | 0.836 | 0.398 | 4.34E-56 |
| | LITAF | 0.919 | 0.805 | 0.411 | 2.92E-47 |
| | SMC4 | 0.909 | 0.332 | 0.136 | 6.33E-16 |
| | REL | 0.894 | 0.832 | 0.447 | 5.95E-42 |
| | SOX4 | 0.874 | 0.226 | 0.066 | 3.06E-18 |
| | FTH1 | 0.847 | 0.996 | 0.981 | 9.35E-33 |
| | CD55 | 0.823 | 0.717 | 0.346 | 7.00E-37 |
| | CREM | 0.798 | 0.916 | 0.632 | 1.04E-37 |
| | CAST | 0.794 | 0.827 | 0.520 | 2.59E-32 |
| | EZR | 0.781 | 0.925 | 0.620 | 8.89E-41 |
| | CD7 | 0.774 | 0.934 | 0.556 | 2.90E-48 |
| | IKZF2 | 0.768 | 0.327 | 0.049 | 1.48E-76 |
| | AREG | 0.767 | 0.580 | 0.320 | 5.21E-16 |
| | LMNA | 0.766 | 0.558 | 0.344 | 2.99E-10 |
| | JUND | 0.758 | 0.982 | 0.839 | 1.17E-51 |
| | RIPOR2 | 0.720 | 0.522 | 0.210 | 1.99E-29 |
| T-prol | STMN1 | 2.541 | 0.983 | 0.221 | 1.82E-198 |
| | HIST1H4C | 2.259 | 0.888 | 0.344 | 9.30E-91 |
| | TUBB | 2.190 | 0.961 | 0.233 | 1.24E-162 |
| | MKI67 | 2.105 | 0.899 | 0.004 | 0 |
| | TUBA1B | 2.034 | 1.000 | 0.494 | 9.40E-100 |
| | CENPF | 1.995 | 0.775 | 0.027 | 0 |
| | TOP2A | 1.987 | 0.798 | 0.008 | 0 |
| | HMGB2 | 1.877 | 0.978 | 0.432 | 3.64E-108 |
| | HMGN2 | 1.871 | 0.994 | 0.355 | 1.82E-129 |
| | ASPM | 1.814 | 0.758 | 0.004 | 0 |
| | NUSAP1 | 1.688 | 0.837 | 0.027 | 0 |
| | PCLAF | 1.643 | 0.781 | 0.005 | 0 |
| | TYMS | 1.580 | 0.753 | 0.008 | 0 |
| | SMC4 | 1.478 | 0.865 | 0.133 | 5.77E-193 |
| | PCNA | 1.451 | 0.725 | 0.086 | 4.30E-202 |
| | CKS1B | 1.437 | 0.815 | 0.072 | 0 |
| | DUT | 1.424 | 0.848 | 0.178 | 2.05E-133 |
| | ACTB | 1.388 | 0.994 | 0.873 | 2.76E-55 |
| | UBE2C | 1.360 | 0.635 | 0.001 | 0 |
| | BIRC5 | 1.308 | 0.702 | 0.001 | 0 |

| Cell state | Gene | Average LogFC | pct1 | pct2 | Adjusted P-value |
|---|---|---|---|---|---|
| | | Myeloid cells | | | |
| cDC1 | CPVL | 2.097 | 0.955 | 0.509 | 8.94E-193 |
| | DNASE1L3 | 2.038 | 0.904 | 0.017 | 0 |
| | CPNE3 | 1.802 | 0.958 | 0.206 | 0 |
| | C1orf54 | 1.748 | 0.952 | 0.233 | 0 |
| | TACSTD2 | 1.741 | 0.850 | 0.122 | 0 |
| | S100B | 1.676 | 0.599 | 0.047 | 0 |
| | IDO1 | 1.656 | 0.790 | 0.050 | 0 |
| | LGALS2 | 1.482 | 0.928 | 0.162 | 0 |
| | CLEC9A | 1.386 | 0.868 | 0.035 | 0 |
| | WDFY4 | 1.383 | 0.907 | 0.099 | 0 |
| | HLA-DPB1 | 1.370 | 0.997 | 0.846 | 1.35E-163 |
| | CST3 | 1.356 | 0.982 | 0.945 | 4.35E-126 |
| | SNX3 | 1.343 | 0.979 | 0.730 | 3.21E-159 |
| | RAB11FIP1 | 1.267 | 0.907 | 0.253 | 1.15E-207 |
| | HLA-DPA1 | 1.181 | 0.988 | 0.866 | 1.68E-137 |
| | PPA1 | 1.143 | 0.958 | 0.352 | 3.53E-165 |
| | EEF1B2 | 1.092 | 0.988 | 0.885 | 5.17E-160 |
| | DAPP1 | 1.088 | 0.826 | 0.129 | 0 |
| | C12orf45 | 1.062 | 0.796 | 0.218 | 6.03E-163 |
| | CADM1 | 1.054 | 0.835 | 0.080 | 0 |
| cDC2 | FCER1A | 1.639 | 0.657 | 0.099 | 0 |
| | AREG | 1.422 | 0.686 | 0.305 | 0 |
| | CD1C | 1.348 | 0.576 | 0.032 | 0 |
| | IL1R2 | 1.316 | 0.696 | 0.192 | 0 |
| | CLEC10A | 1.272 | 0.727 | 0.143 | 0 |
| | LYZ | 1.149 | 0.957 | 0.641 | 0 |
| | CST3 | 1.111 | 0.971 | 0.940 | 0 |
| | CCL22 | 1.018 | 0.237 | 0.039 | 0 |
| | HLA-DQA1 | 0.981 | 0.947 | 0.657 | 0 |
| | HLA-DPB1 | 0.881 | 0.989 | 0.823 | 0 |
| | AC020656.1 | 0.877 | 0.638 | 0.255 | 0 |
| | LGALS2 | 0.849 | 0.596 | 0.097 | 0 |
| | CCR7 | 0.829 | 0.332 | 0.105 | 1.58E-257 |
| | HLA-DQB1 | 0.824 | 0.959 | 0.693 | 0 |
| | CFP | 0.819 | 0.619 | 0.147 | 0 |
| | CRIP1 | 0.805 | 0.640 | 0.274 | 0 |
| | CST7 | 0.787 | 0.412 | 0.089 | 0 |
| | HLA-DRA | 0.784 | 0.995 | 0.942 | 0 |
| | HLA-DRB1 | 0.765 | 0.985 | 0.879 | 0 |
| | FCN1 | 0.758 | 0.493 | 0.126 | 0 |
| mDC | CCL22 | 2.962 | 0.770 | 0.062 | 0 |
| | BIRC3 | 2.952 | 0.995 | 0.425 | 3.54E-167 |
| | CCR7 | 2.767 | 0.995 | 0.132 | 0 |
| | IDO1 | 2.441 | 0.761 | 0.054 | 0 |
| | CCL17 | 2.142 | 0.446 | 0.024 | 0 |
| | TXN | 2.002 | 0.991 | 0.782 | 2.69E-99 |
| | LAMP3 | 1.972 | 0.937 | 0.034 | 0 |
| | NUB1 | 1.921 | 0.905 | 0.188 | 2.02E-213 |
| | MARCKSL1 | 1.881 | 0.919 | 0.232 | 4.16E-176 |
| | IL7R | 1.848 | 0.950 | 0.213 | 3.55E-189 |
| | FSCN1 | 1.816 | 0.824 | 0.155 | 7.79E-193 |
| | CST7 | 1.795 | 0.824 | 0.132 | 1.81E-221 |
| | DAPP1 | 1.727 | 0.928 | 0.131 | 0 |
| | GADD45A | 1.701 | 0.766 | 0.165 | 4.07E-151 |
| | RAMP1 | 1.674 | 0.671 | 0.067 | 1.79E-275 |
| | CRIP1 | 1.665 | 0.887 | 0.325 | 1.10E-91 |
| | RAB9A | 1.573 | 0.887 | 0.259 | 3.45E-153 |
| | CCL19 | 1.526 | 0.392 | 0.007 | 0 |
| | EBI3 | 1.509 | 0.649 | 0.199 | 1.06E-77 |
| | DUSP5 | 1.474 | 0.937 | 0.254 | 1.50E-162 |

**Supplementary Table 10: Top Gene Markers Expressed for Breast Cell States**

| Cell state | Gene | Average LogFC | pct1 | pct2 | Adjusted P-value |
|---|---|---|---|---|---|
| Myeloid cells | | | | | |
| pDC | GZMB | 3.653 | 0.969 | 0.009 | 0 |
| | PTGDS | 2.965 | 0.462 | 0.027 | 1.08E-99 |
| | IGKC | 2.673 | 0.954 | 0.376 | 1.00E-35 |
| | JCHAIN | 2.549 | 0.969 | 0.108 | 2.29E-127 |
| | IRF4 | 2.131 | 0.831 | 0.090 | 4.59E-106 |
| | CLIC3 | 2.066 | 0.754 | 0.006 | 0 |
| | PLAC8 | 2.062 | 0.754 | 0.053 | 6.76E-140 |
| | C12orf75 | 1.924 | 0.800 | 0.086 | 8.63E-102 |
| | ITM2C | 1.915 | 0.800 | 0.144 | 6.17E-61 |
| | TSPAN13 | 1.882 | 0.785 | 0.030 | 1.68E-267 |
| | TCL1A | 1.826 | 0.277 | 0.000 | 0 |
| | IRF7 | 1.813 | 0.862 | 0.200 | 3.57E-53 |
| | LILRA4 | 1.775 | 0.692 | 0.017 | 0 |
| | TCF4 | 1.735 | 0.938 | 0.418 | 5.81E-34 |
| | IRF8 | 1.718 | 0.815 | 0.256 | 3.65E-34 |
| | PPP1R14B | 1.716 | 0.954 | 0.513 | 9.96E-32 |
| | CXCR3 | 1.715 | 0.754 | 0.015 | 0 |
| | BCL11A | 1.698 | 0.738 | 0.050 | 5.26E-144 |
| | SOX4 | 1.670 | 0.723 | 0.182 | 2.04E-33 |
| | MZB1 | 1.599 | 0.754 | 0.007 | 0 |
| Mast | TPSAB1 | 3.304 | 0.542 | 0.007 | 0 |
| | TPSB2 | 3.278 | 0.550 | 0.007 | 0 |
| | CTSG | 3.025 | 0.496 | 0.002 | 0 |
| | HPGD | 3.014 | 0.618 | 0.086 | 2.45E-116 |
| | CPA3 | 2.684 | 0.595 | 0.001 | 0 |
| | GATA2 | 2.648 | 0.603 | 0.003 | 0 |
| | HDC | 2.508 | 0.618 | 0.001 | 0 |
| | ATP10D | 2.432 | 0.336 | 0.092 | 1.98E-22 |
| | IL1RL1 | 2.326 | 0.611 | 0.016 | 0 |
| | BIRC3 | 2.121 | 0.626 | 0.429 | 1.15E-15 |
| | ADCYAP1 | 1.950 | 0.176 | 0.001 | 0 |
| | MCTP2 | 1.852 | 0.359 | 0.018 | 2.40E-167 |
| | PRKX | 1.831 | 0.450 | 0.171 | 1.41E-21 |
| | CLC | 1.786 | 0.221 | 0.000 | 0 |
| | GK5 | 1.777 | 0.275 | 0.110 | 5.55085E-08 |
| | KIT | 1.774 | 0.405 | 0.011 | 0 |
| | HPGDS | 1.767 | 0.466 | 0.124 | 3.51E-37 |
| | GPR65 | 1.723 | 0.412 | 0.217 | 1.0128E-09 |
| | MS4A2 | 1.719 | 0.458 | 0.001 | 0 |
| | SELENOK | 1.703 | 0.649 | 0.620 | 3.73E-10 |
| Mono-nonclassical | LST1 | 2.153 | 0.987 | 0.523 | 0 |
| | CD52 | 1.824 | 0.869 | 0.188 | 0 |
| | IFITM2 | 1.658 | 0.956 | 0.549 | 0 |
| | FCGR3A | 1.639 | 0.950 | 0.330 | 0 |
| | SERPINA1 | 1.610 | 0.962 | 0.362 | 0 |
| | COTL1 | 1.598 | 0.985 | 0.661 | 0 |
| | SMIM25 | 1.542 | 0.873 | 0.216 | 0 |
| | FCN1 | 1.517 | 0.817 | 0.160 | 0 |
| | STXBP2 | 1.376 | 0.878 | 0.308 | 0 |
| | LILRA5 | 1.324 | 0.759 | 0.069 | 0 |
| | CORO1A | 1.249 | 0.897 | 0.360 | 0 |
| | LILRB2 | 1.237 | 0.877 | 0.326 | 0 |
| | S100A4 | 1.203 | 0.991 | 0.745 | 1.91E-262 |
| | CD48 | 1.190 | 0.883 | 0.322 | 0 |
| | AIF1 | 1.183 | 0.991 | 0.794 | 2.37E-274 |
| | CDKN1C | 1.159 | 0.581 | 0.104 | 0 |
| | LYPD2 | 1.155 | 0.215 | 0.000 | 0 |
| | RIPOR2 | 1.146 | 0.621 | 0.045 | 0 |
| | SPN | 1.140 | 0.722 | 0.097 | 0 |
| | PLAC8 | 1.138 | 0.531 | 0.038 | 0 |

| Cell state | Gene | Average LogFC | pct1 | pct2 | Adjusted P-value |
|---|---|---|---|---|---|
| Myeloid cells | | | | | |
| Mono-classical | CXCL5 | 2.609 | 0.485 | 0.097 | 0 |
| | SERPINB2 | 2.390 | 0.301 | 0.020 | 0 |
| | EREG | 1.863 | 0.787 | 0.180 | 0 |
| | VCAN | 1.663 | 0.735 | 0.219 | 0 |
| | S100A9 | 1.660 | 0.718 | 0.366 | 1.54E-301 |
| | S100A8 | 1.655 | 0.515 | 0.200 | 2.00E-266 |
| | THBS1 | 1.561 | 0.889 | 0.395 | 0 |
| | CCL20 | 1.398 | 0.549 | 0.171 | 0 |
| | G0S2 | 1.307 | 0.686 | 0.285 | 0 |
| | PTGS2 | 1.301 | 0.500 | 0.181 | 1.78E-294 |
| | TIMP1 | 1.278 | 0.960 | 0.675 | 0 |
| | IL1B | 1.277 | 0.784 | 0.426 | 0 |
| | CXCL3 | 1.241 | 0.856 | 0.451 | 0 |
| | CXCL1 | 1.238 | 0.468 | 0.212 | 5.83E-171 |
| | S100A12 | 1.193 | 0.313 | 0.025 | 0 |
| | TNFAIP6 | 1.085 | 0.366 | 0.128 | 2.48E-205 |
| | LYZ | 1.078 | 0.900 | 0.667 | 3.82E-282 |
| | AQP9 | 1.054 | 0.574 | 0.118 | 0 |
| | FCN1 | 1.015 | 0.564 | 0.141 | 0 |
| | MT1G | 1.011 | 0.137 | 0.066 | 2.35E-30 |
| Macro-lipo | FABP4 | 3.612 | 0.630 | 0.237 | 1.63E-230 |
| | APOC1 | 2.757 | 0.811 | 0.247 | 0 |
| | SPP1 | 2.088 | 0.560 | 0.136 | 5.28E-297 |
| | APOE | 1.984 | 0.861 | 0.467 | 7.55E-271 |
| | LIPA | 1.562 | 0.841 | 0.377 | 0 |
| | ACP5 | 1.552 | 0.797 | 0.199 | 0 |
| | FABP5 | 1.518 | 0.887 | 0.550 | 7.17E-214 |
| | GPNMB | 1.453 | 0.876 | 0.406 | 0 |
| | CSTB | 1.311 | 0.972 | 0.821 | 3.40E-262 |
| | MMP9 | 1.290 | 0.478 | 0.219 | 2.45E-85 |
| | CHIT1 | 1.290 | 0.251 | 0.006 | 0 |
| | LGALS3 | 1.283 | 0.932 | 0.729 | 7.72E-231 |
| | CTSD | 1.259 | 0.955 | 0.739 | 7.03E-254 |
| | CHI3L1 | 1.191 | 0.183 | 0.027 | 7.67E-150 |
| | CD9 | 1.169 | 0.795 | 0.380 | 9.46E-227 |
| | CD52 | 1.168 | 0.595 | 0.194 | 1.87E-200 |
| | CAPG | 1.135 | 0.730 | 0.384 | 1.38E-154 |
| | CYP27A1 | 1.133 | 0.593 | 0.060 | 0 |
| | LPL | 1.067 | 0.531 | 0.134 | 5.83E-255 |
| | CD36 | 1.031 | 0.621 | 0.300 | 2.24E-112 |
| Macro-m1 | C3 | 1.942 | 0.753 | 0.116 | 0 |
| | RGS1 | 1.794 | 0.867 | 0.459 | 0 |
| | FCGBP | 1.382 | 0.405 | 0.091 | 0 |
| | PLCG2 | 1.174 | 0.322 | 0.319 | 1 |
| | OLR1 | 1.107 | 0.546 | 0.177 | 0 |
| | APOE | 1.095 | 0.819 | 0.374 | 0 |
| | SDS | 0.993 | 0.444 | 0.174 | 0 |
| | HERPUD1 | 0.967 | 0.874 | 0.678 | 0 |
| | CXCR4 | 0.950 | 0.751 | 0.510 | 0 |
| | PLXDC2 | 0.943 | 0.740 | 0.338 | 0 |
| | YWHAH | 0.908 | 0.733 | 0.438 | 0 |
| | SGK1 | 0.899 | 0.741 | 0.556 | 0 |
| | ALOX5AP | 0.888 | 0.795 | 0.486 | 0 |
| | ZNF331 | 0.888 | 0.512 | 0.237 | 0 |
| | AXL | 0.876 | 0.623 | 0.194 | 0 |
| | FCGR3A | 0.859 | 0.669 | 0.248 | 0 |
| | SAT1 | 0.816 | 0.998 | 0.979 | 0 |
| | HLA-DPA1 | 0.809 | 0.967 | 0.835 | 0 |
| | RGS2 | 0.807 | 0.742 | 0.475 | 0 |
| | CD81 | 0.798 | 0.834 | 0.663 | 0 |

**Supplementary Table 10: Top Gene Markers Expressed for Breast Cell States**

| Cell state | Gene | Average LogFC | pct1 | pct2 | Adjusted P-value |
|---|---|---|---|---|---|
| | | **Myeloid cells** | | | |
| **Macro-m2** | HMOX1 | 2.214 | 0.850 | 0.524 | 0 |
| | RNASE1 | 2.108 | 0.982 | 0.300 | 0 |
| | SELENOP | 1.869 | 0.894 | 0.273 | 0 |
| | LYVE1 | 1.868 | 0.771 | 0.084 | 0 |
| | EMP1 | 1.631 | 0.801 | 0.319 | 0 |
| | F13A1 | 1.570 | 0.749 | 0.221 | 0 |
| | MRC1 | 1.450 | 0.903 | 0.322 | 0 |
| | PLTP | 1.323 | 0.756 | 0.208 | 0 |
| | LGMN | 1.297 | 0.840 | 0.389 | 0 |
| | FOLR2 | 1.282 | 0.771 | 0.235 | 0 |
| | CCL18 | 1.249 | 0.301 | 0.036 | 0 |
| | CD163 | 1.242 | 0.962 | 0.558 | 0 |
| | CCL2 | 1.238 | 0.474 | 0.177 | 0 |
| | MAN1A1 | 1.208 | 0.786 | 0.299 | 0 |
| | LILRB5 | 1.160 | 0.664 | 0.064 | 0 |
| | PMP22 | 1.100 | 0.677 | 0.203 | 0 |
| | STAB1 | 1.099 | 0.792 | 0.284 | 0 |
| | GCLM | 1.074 | 0.631 | 0.249 | 0 |
| | MARCO | 1.072 | 0.438 | 0.166 | 0 |
| | CCL4 | 1.064 | 0.630 | 0.359 | 0 |
| **Macro-ifn** | CXCL10 | 2.925 | 0.439 | 0.032 | 4.21E-207 |
| | ISG15 | 2.674 | 0.975 | 0.226 | 1.35E-202 |
| | IFIT3 | 2.450 | 0.869 | 0.041 | 0 |
| | IFIT1 | 2.414 | 0.884 | 0.022 | 0 |
| | RSAD2 | 2.365 | 0.803 | 0.027 | 0 |
| | IFIT2 | 2.349 | 0.732 | 0.039 | 0 |
| | CCL8 | 2.299 | 0.495 | 0.052 | 3.67E-161 |
| | MX1 | 2.104 | 0.955 | 0.126 | 0 |
| | IFI6 | 1.683 | 0.904 | 0.214 | 3.20E-152 |
| | IFI44L | 1.661 | 0.894 | 0.069 | 0 |
| | PARP14 | 1.542 | 0.934 | 0.398 | 1.57E-109 |
| | MX2 | 1.430 | 0.869 | 0.171 | 1.38E-179 |
| | RNF213 | 1.413 | 0.924 | 0.465 | 1.02E-85 |
| | CCL2 | 1.354 | 0.682 | 0.278 | 1.19E-44 |
| | EIF2AK2 | 1.350 | 0.879 | 0.204 | 1.15E-156 |
| | IFIH1 | 1.319 | 0.747 | 0.118 | 4.35E-181 |
| | OAS1 | 1.312 | 0.768 | 0.105 | 6.57E-209 |
| | SAMD9L | 1.287 | 0.712 | 0.093 | 1.17E-204 |
| | TNFSF10 | 1.245 | 0.485 | 0.071 | 2.53E-109 |
| | XAF1 | 1.239 | 0.737 | 0.093 | 2.57E-219 |
| **Myeloid-prol** | STMN1 | 1.685 | 0.612 | 0.315 | 2.69E-71 |
| | HIST1H4C | 1.573 | 0.466 | 0.321 | 8.96E-19 |
| | HMGN2 | 1.437 | 0.690 | 0.590 | 1.17E-46 |
| | HMGB2 | 1.296 | 0.531 | 0.383 | 1.02E-24 |
| | TUBA1B | 1.241 | 0.746 | 0.852 | 1.54E-37 |
| | MKI67 | 1.189 | 0.399 | 0.005 | 0 |
| | TOP2A | 1.166 | 0.383 | 0.010 | 0 |
| | PCLAF | 1.121 | 0.399 | 0.014 | 0 |
| | TUBB | 1.115 | 0.634 | 0.661 | 3.79E-19 |
| | H2AFZ | 1.091 | 0.701 | 0.741 | 1.47E-33 |
| | HMGB1 | 1.091 | 0.791 | 0.888 | 5.28E-53 |
| | CKS1B | 1.050 | 0.480 | 0.135 | 5.34E-92 |
| | FCER1A | 1.026 | 0.318 | 0.183 | 2.74E-10 |
| | CENPF | 0.993 | 0.344 | 0.018 | 0 |
| | TYMS | 0.957 | 0.380 | 0.015 | 0 |
| | NUSAP1 | 0.945 | 0.369 | 0.029 | 1.78E-273 |
| | UBE2C | 0.926 | 0.291 | 0.002 | 0 |
| | CDK1 | 0.918 | 0.344 | 0.019 | 0 |
| | AC020656.1 | 0.883 | 0.422 | 0.312 | 0.0004303 |
| | PTTG1 | 0.881 | 0.377 | 0.106 | 8.58E-64 |

**Supplementary Table 10: Top Gene Markers Expressed for Breast Cell States**

This table lists the top marker genes expressed for each major cell state cluster based on the average log fold-change within the respective cell type and specificity of the gene expressed in the indicated cell state (pct1) compared to the other cell states of the same cell type (pc2). The columns listed (from left to right) indicate the cell state name, the gene names, the average log-fold change, the pct1 value indicating the fraction of cells within the cluster expressing the gene, the pct2 value indicating the fraction of cells in other clusters expressing the gene, and the Bonferroni adjusted p-value.
